# Substrate binding and activation mechanism of the essential bacterial septal cell wall synthase FtsW

**DOI:** 10.64898/2026.02.13.705525

**Authors:** Martín Yepes, Nicolás N. Yehya, Amilcar Perez, Ruiwen Xiong, Cole Christensen, Brooke Britton, Guy W. Dayhoff, Xinzhe Ren, Caden Sing Hei Lok, Stephen D. Fried, Jana Shen, Albert Y. Lau, Jie Xiao

## Abstract

Septal peptidoglycan (sPG) synthesis in most bacteria is driven by the essential, highly conserved FtsWIQLB synthase complex, comprising the SEDS (Shape, Elongation, Division, and Sporulation) glycosyltransferase FtsW, the monofunctional transpeptidase FtsI, and the scaffolding and regulatory subcomplex FtsQLB. Although apo structures of FtsWIQLB are known, no substrate-bound structures exist, likely due to the intrinsic flexibility of lipid II (L2) substrates. Here, we combined all-atom molecular dynamics simulations of *E. coli* FtsWIQLB with cysteine-based mutagenesis and cell-based assays to investigate donor and acceptor L2 binding sites in FtsW and to elucidate the complex’s activation mechanism. We show that conserved arginine residues adjacent to membrane-accessible cavities in FtsW coordinate donor and acceptor L2 pyrophosphate groups, stabilizing substrate binding. The FtsWIQLB complex appears to adopt a self-inhibitory architecture in which gating elements and a periplasmic loop prevent the donor and acceptor from approaching FtsW’s catalytic residue D297. Activation by FtsN or a superfission FtsW variant relieves these constraints, stabilizes acceptor-site binding, and reorients both substrates into a catalytically primed state. Long donor glycan chains further stabilize this activated conformation at the acceptor site, possibly promoting processive sPG polymerization. Comparative modeling of ESKAPE pathogen homologs reveals both conserved and divergent features of binding-site engagement and activation. We corroborate these computational findings with cysteine-based mutagenesis and crosslinking experiments. These results establish a mechanistic framework for FtsW substrate recognition and functional activation, highlighting tractable sites for structure- and mechanism-based antibiotic discovery targeting SEDS-class cell wall synthases.

## INTRODUCTION

The bacterial cell wall is composed of cross-linked peptidoglycan (PG). It determines cell shape and provides mechanical strength against intracellular turgor pressure, thereby preventing cell lysis^1^. Consequently, PG synthesis is critical for bacterial growth and survival. PG is polymerized from the universal precursor lipid II (L2, consisting of a disaccharide GlcNAc and MurNAc and a C55 lipid tail) by glycosyltransferases (GTase) and crosslinked by transpeptidases (TPase) to the existing cell wall (**Fig. 1A, Fig. S1A).** Bifunctional class-A penicillin-binding proteins (aPBP) contain both GTase and TPase domains; they were traditionally regarded as the main PG synthases^2,3^. However, recent studies have shown that a new class of PG synthases, formed by a SEDS (Shape, Elongation, Division, and Sporulation) GTase and a monofunctional class-B PBP with only TPase activity (SEDS:bPBP), is responsible for the bulk of cell wall synthesis^4,5^. Because SEDS GTases are more broadly conserved than aPBPs^5^, adopt a unique structural fold^6–8^, and are essential for cell survival^4,5^, they are attractive targets for new antimicrobial development, especially amid the growing β-lactam resistance targeting TPases^2,9,10^

**Figure 1.**
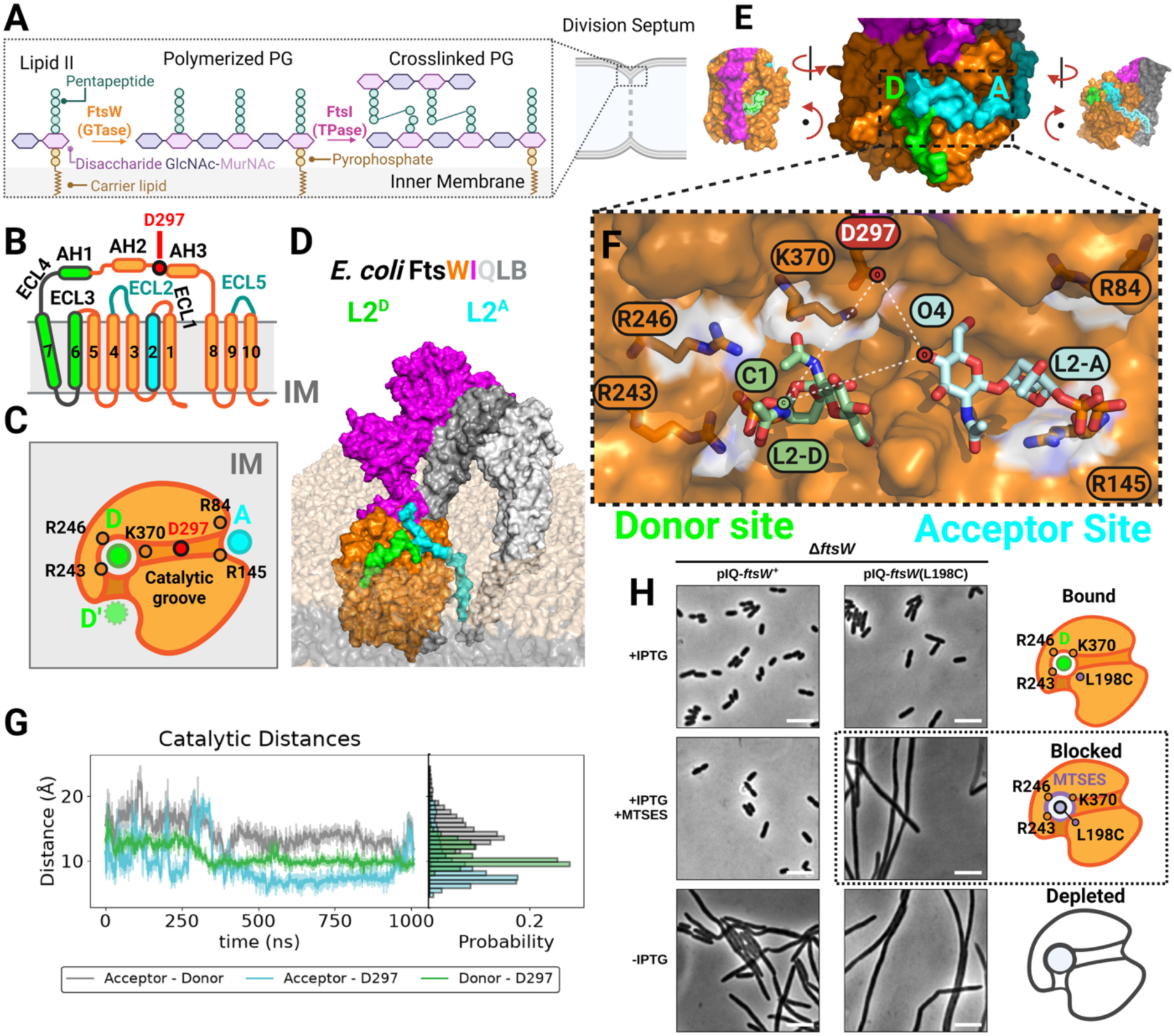
Identifying putative L2 donor and acceptor sites in FtsW by molecular dynamics (MD) simulations and targeted mutagenesis. **(A)** Schematic of lipid II polymerization by glycosyltransferase (GTase) FtsW and crosslinking by transpeptidase (TPase) FtsI. **(B)** Topology diagram of FtsW showing ten transmembrane helices (TM1–TM10), five extracellular loops (ECL1–ECL5), and three amphipathic helices (AH1–AH3) in ECL4. Regions implicated in substrate binding are highlighted in green (donor site), cyan (acceptor site), or teal (contributing to both sites). The putative catalytic residue D297 is shown in red. **(C)** Schematic of the catalytic groove and conserved coordinating residues. The catalytic aspartate D297 (red) lies between a cluster of donor-site arginines (R246, R243) and acceptor-site arginines (R84, R145), defining a shared catalytic groove flanked by two substrate-binding subsites. **(D)** MD structural model of the *E. coli* FtsWIQLB complex with an L2 donor (green) and an L2 acceptor (cyan), shown in surface representation. Inner membrane (IM) lipids are shown as gray space-filled model. **(E)** Periplasmic and side views of the structural model of FtsW in the FtsWIQLB complex with two L2 molecules bound in the donor site and the acceptor site, shown as surfaces. Middle: FtsW catalytic groove viewed from the periplasm. Left: Donor cavity viewed from the inner membrane and rotated from the middle view, with L2 C55 tail carbons shown as sticks in a transparent green surface. Right: Acceptor site viewed from the inner membrane and rotated from the middle view, with L2 C55 tail carbons shown as sticks in a transparent cyan surface. **(F)** FtsW catalytic groove viewed from the periplasm and shown as a semi-transparent surface, with key basic residues (orange), the putative catalytic residue (D297, red), and L2 head groups (green and cyan for donor and acceptor, respectively) shown as sticks. The C1 and O4 atoms of L2^D^ and L2^A^, respectively, are labeled. **(G)** Representative distance time traces and accompanying histograms from 1 µs equilibrium MD simulations. Shown are the shortest distance between the donor reducing-end C1 atom and the acceptor nonreducing-end O4 atom (donor–acceptor, grey), shortest distance between the D297 carboxylate and the acceptor O4 atom (D297–acceptor, cyan), and shortest distance between the D297 carboxylate and the donor reducing-end C1 atom (D297–donor, green). **(H)** Phase contrast micrographs comparing effects of cells expressing *ftsW^+^* (left column, strain XY396), or *ftsW*(L198C) (right column, strain AP852) as sole copies of *ftsW* under control of an IPTG-inducible promoter in M9 at 25 °C. Rows show cells grown with IPTG (top), with IPTG and MTSES (middle), and without IPTG (FtsW depletion, bottom). The cartoons on the right depict the proposed state of the catalytic groove in each case: lipid II bound at the donor site (Bound), MTSES-modified FtsW(L198C) blocking the donor site (Blocked; dotted box marks the condition that produces filamentation), and no FtsW (Depleted). Micrographs are representative of three independent biological replicates. Quantifications of cell lengths were included in **Supplementary Table 4**.

The SEDS protein FtsW belongs to the divisome and was recently identified as the essential septal PG (sPG) GTase during cell division^4^ (**Fig. 1A**). It forms the core bacterial sPG synthesis complex with its cognate TPase partner FtsI (PBP3), a class-B monofunctional PBP, and a tripartite scaffolding and regulatory subcomplex FtsQLB^11–14^. The FtsWIQLB complex is conserved in nearly all bacteria; their septal localization and function are essential for sPG synthesis and cell wall constriction^15,16^

In many gammaproteobacteria, the FtsWIQLB complex is further activated by a highly conserved protein FtsN^17^. FtsN engages with multiple divisome components: its cytoplasmic N-terminus contacts FtsA to relay an activation signal to FtsW^14,18^; genetic evidence suggests that its essential E domain in the periplasm activates FtsW^19^, likely through its binding to FtsI and FtsL at the AWI (Activation of W and I) region to trigger conformational changes^19,20^; and its SPOR domain oligomerizes on denuded glycans (dnG) in the septal cell wall to protect them from degradation by lytic glycosyltransferases (LTs)^21^. These interactions allow FtsN to coordinate sPG synthesis and degradation during septal cell wall constriction^21^.

The activated FtsWIQLB complex exhibits high processivity in sPG synthesis. Single-molecule imaging studies from our group and others have shown that the complex can synthesize septal sPG chains several hundred to over a thousand nanometers long in a single processive run in live cells^22,23,24,25^. The mechanism underlying this remarkable processivity is unclear, but the glycosyltransfer activity of FtsW is likely a central driver.

In a proposed *S_N_*2-like inverting glycosyltransfer reaction mechanism for RodA^8^, the elongasome SEDS paralog of FtsW with a highly similar structural fold^5^, the putative catalytic residue D262 of RodA (D297 in FtsW) deprotonates the 4′-OH of the N-acetylglucosamine (GlcNAc) in the acceptor L2 molecule, rendering it nucleophilic to attack the 1′-carbon of the N-acetylmuramic acid (MurNAc) in the donor L2 or nascent PG chain molecule with an inversion of stereochemistry (**Fig. S1B**). This reaction cleaves off the donor’s undecaprenyl-pyrophosphate C55 lipid tail (Und-PP) and transfers its sugar moiety to the acceptor L2’s nonreducing end^4^, forming the β-1,4 glycosidic bond^26^.

Previously, we used a combination of structural prediction, molecular dynamics (MD) simulations, and mutagenesis to model the structure of the *E. coli* FtsWIQLB complex and the activator protein FtsN^20^. In the model, the heterotrimer FtsQLB scaffolds FtsI in an upright, stable conformation, with the essential domain of FtsN (N^E^) binding to the AWI region of FtsI and FtsL, likely modulating the conformation of FtsI’s anchor domain and the interface with FtsW ECL4^20^. The AlphaFold and MD-derived structural model of the *E. coli* FtsWIQLB complex closely matched the cryoEM-resolved *Pseudomonas aeruginosa* FtsWIQLB structure^7,27^ and the X-ray crystallographic structure of the *E. coli* FtsQLB complex^28^, validating the combined structural prediction and MD modeling approach and underscoring the complex’s high conservation. However, L2 substrates are not visible in cryoEM structures^7,27^, possibly because of their dynamic, flexible, and variable binding modes. The lack of an atomic understanding of how L2 substrates bind to FtsW makes it challenging to define the enzymatic mechanism for potential antibiotic development.

In this work, we combined all-atom MD simulations of *E. coli* FtsWIQLB with cysteine-based mutagenesis and cell-based assays to identify putative donor and acceptor L2 binding sites in FtsW and to elucidate its activation mechanism. We uncovered conserved electrostatic contacts and conformational switches required for function and propose a model linking substrate recognition to FtsW activation and processivity in sPG synthesis. Together, these findings establish a molecular framework that opens new avenues for mechanism-based antibiotic discovery targeting SEDS-class cell wall synthases.

## RESULTS

### Assigning putative L2 donor and acceptor sites in FtsW

To investigate how FtsW binds its L2 substrates for catalysis, we first examined FtsW for potential L2-binding sites. *E. coli* FtsW has ten transmembrane (TM) helices, with the putative catalytic residue D297 located at the start of the amphipathic helix 3 (AH3) within the large extracellular loop ECL4 (**Fig. 1B**). We reasoned that the putative donor and acceptor sites in FtsW with respect to D297 must satisfy the following four criteria. First, key amino acids at the binding sites must be highly conserved across FtsW and related enzymes, as L2 polymerization is a universal reaction in bacterial cell wall synthesis^1^, and the chemical structure of L2 varies only slightly across species^29,30^. Second, the binding pockets must contain solvent-exposed, positively charged residues near D297 to coordinate the negatively charged pyrophosphate group linking the lipid tail to the disaccharide moiety of L2. Third, the geometry of the two binding sites must orient the reactive atoms of the donor and acceptor substrates correctly with respect to D297 to enable catalysis. Fourth, as L2 diffuses freely within the inner membrane, the periplasmic binding sites must lie adjacent to hydrophobic cavities in FtsW that can accommodate the lipid tails of the substrates.

Based on these criteria, we assigned the cavity formed by FtsW AH1, TM6, and TM7 as the putative PG donor site and the cavity formed by FtsW TM2 as the L2 acceptor site (**Fig. 1B**, green and cyan segments for the donor and acceptor sites, respectively). ECL2 and ECL5 encompass both the donor and acceptor sites (**Fig. 1B**, teal segments). The donor site lies within FtsW, while the acceptor site opens directly to the outside (**Fig. 1C**, green and cyan circles).

The putative donor site corresponds to a cavity occupied by a lipid density in the cryoEM structures of the FtsW paralog RodA^8,31^ and WaaL, a GTase belonging to the same GT-C_B_ class as FtsW^31^. The lipid density in these cryoEM structures was attributed to partially ordered Und-PP, which would be found only at the donor site after the glycosyltransfer reaction. The putative acceptor site coincided with a high L2-density-associated cavity identified in a previous MS simulation study of RodA^8^.

Most importantly, the two sites contain several conserved, positively charged arginine residues, as revealed by sequence alignment analysis^32^ across 6859 bacterial species (**Table 1**, Methods). These solvent-exposed arginines likely play a critical role in coordinating the pyrophosphates of the donor and acceptor L2 molecules, respectively. For example, R246 and R243 in AH1 of ECL4 at the putative donor site are highly conserved (∼99% and ∼77%, respectively; **Table 1, Fig. S3).** Both produced a dominant-negative phenotype when mutated to H and L, P, or S, respectively^33^. At the putative acceptor site, R84 in ECL1 is also highly conserved (∼70%, most often replaced by lysine, **Table 1**, **Fig. S3).** R145 in ECL2 is less conserved, but its mutations failed to complement an *E. coli* FtsW temperature-sensitive strain at 42°C^34^. Note that in the GT-C_B_ family of enzymes, the Und-PP-linked substrates are always coordinated by conserved arginines at the donor site, such as R48, R109, and R210 in RodA (corresponding to R84, R145, and R246 in FtsW, respectively, **Fig. S2B)**^8,31^. The sequence alignment analysis also identified the known LPEAHTD motif containing the putative catalytic residue (D297) in FtsW (**Fig. S3**), as previously noted^7,8^. D297 lies in a similar structural location and orientation as the putative catalytic residue D262 in RodA (**Fig. S2B**) and H313 in WaaL (**Fig. S2D**).

**Table 1.** Conserved Positively Charged Residues in FtsW using Clustal W analysis^32^ across 685G bacterial species.

| **Index** | **Residue** | **Conservation Score** | **Solvent Exposed** | **Dominant Negative**<sup>33</sup> |
|---|---|---|---|---|
| 246 | R | 0.9965 | Yes | Yes |
| 287 | K | 0.9799 | No | No |
| 153 | K | 0.9692 | No | No |
| 166 | R | 0.9621 | No | No |
| 137 | R | 0.7807 | Yes | Yes |
| 243 | R | 0.7712 | Yes | Yes |
| 84 | R | 0.7140 | Yes | No |

Structurally, R246 and R243 at the donor site and R84 and R145 at the acceptor site flank D297 with a distance of less than 20 Å, enclosing a periplasmic cavity that we defined as the catalytic groove (**Fig. 1C**, central cleft). Notably, both sites encompass hydrophobic cavities facing the membrane (**Fig. S4A**), making it possible to accommodate L2’s long hydrophobic lipid tail. As such, the structural and chemical properties of the two sites fulfill the criteria proposed for putative donor and acceptor sites in FtsW.

### MD simulations reveal that FtsW accommodates individual lipid II molecules at either putative binding site

To investigate whether each of the two putative binding sites in FtsW could indeed accommodate an L2 molecule, we extracted the atomic coordinates from our previously published MD trajectories of the FtsWIQLB apo system^20^ and examined snapshots where FtsW’s TM7 opens outward to create enough space for the polar headgroups of membrane lipids to approach the catalytic groove unobstructed (**Fig. S4B**). Interestingly, as shown in **Supplementary Movie 1**, a 1-palmitoyl-2-oleoyl-phosphatidylethanolamine (POPE) lipid in the outer leaflet appeared to slither through an opening between FtsW’s TM6 and TM7 to reach the putative donor site. At ∼333 ns, one tail of the POPE lipid partially entered the putative donor cavity, and FtsW R243 and R246 both contacted the POPE phosphate head group.

Using this snapshot, we manually docked an *E. coli* L2 molecule into the putative donor site and rebuilt the protein-ligand system in a lipid bilayer and solvent box (Methods; hereafter referred to as the FtsWIQLB-L2^D^ simulation, **Table 2**). We then performed 52 ns all-atom equilibrium MD simulations at 300K using the NAMD MD software platform^35,36^ to allow the L2 molecule to form favorable contacts or diffuse back into the membrane. Because the putative acceptor site is located at the periphery of FtsW, we repeated the docking process at the Acceptor site using the same MD simulation trajectory snapshot (Methods, FtsWIQLB-L2^A^ simulation, **Table 2**).

**Table 2.** List of MD Simulations and key information.

| **Sim#** | **Name** | **System** | **Simulated Time** | **D** | **A** | **Description** |
|---|---|---|---|---|---|---|
| **1** | FtsWIQLB-L2<sup>D</sup> | WT | 52 ns | L2 | None | 300K, no Mg* |
| **2** | FtsWIQLB-L2<sup>D’</sup> | WT | 52 ns | L2* | None | 300K, no Mg* |
| **2-mtd3** | FtsWIQLB-L2D’ | WT | 188 ns | L2* | None | Biased (well-tempered metadynamics), NAMD, 300K, no Mg* |
| **4** | FtsWIQLB-L2<sup>A</sup> | WT | 52 ns | None | L2 | 300K, no Mg* |
| **8** | FtsWIQLB-L2<sup>D</sup>-L2<sup>A</sup> | WT | 1 µs | L2 | L2 | 300K, no Mg* |
| **14** | FtsWIQLB-L2<sup>D</sup>-L2<sup>A</sup> | WT | 1 µs | L2 | L2 | 310K, with Mg* |
| **14b** | FtsWIQLB-L2<sup>D</sup>-L2<sup>A</sup> | WT | 1 µs | L2 | L2 | Repeat of 14 |
| **15** | FtsW<sup>E289G</sup>IQLB-L2<sup>D</sup>-L2<sup>A</sup> | E289G | 1 µs | L2 | L2 |  |
| **15b** | FtsW<sup>E289G</sup>IQLB-L2<sup>D</sup>-L2<sup>A</sup> | E289G | 1 µs | L2 | L2 | Repeat of 15 |
| **17** | FtsWIQLB-N<sup>58-108</sup>-L2<sup>D</sup>-L2<sup>A</sup> | +FtsN (E) | 4 µs | L2 | L2 | A2 (1 µs) + A3** extension (3 µs) |
| **34** | PAE FtsWIQLB-L2<sup>D</sup>-L2<sup>A</sup> | WT | 100 ns | L2 | L2 |  |
| **35** | BSU FtsWIQLB-L2<sup>D</sup>-L2<sup>A</sup> | WT | 100 ns | L2 | L2 |  |
| **36** | KPN FtsWIQLB-L2<sup>D</sup>-L2<sup>A</sup> | WT | 92 ns | L2 | L2 |  |
| **37** | CCR FtsWIQLB-L2<sup>D</sup>-L2<sup>A</sup> | WT | 100 ns | L2 | L2 |  |
| **42** | FtsWIQLB-N<sup>58-108</sup>-L4<sup>D</sup>-L2<sup>A</sup> | +FtsN (E) | 4 µs | L4 | L2 | A3** |
| **43** | FtsWIQLB-N<sup>58-108</sup>-L32<sup>D</sup>-L2<sup>A</sup> | +FtsN (E) | 2 µs | L32 | L2 | A3** |
| **43b** | FtsWIQLB-N<sup>58-108</sup>-L32<sup>D</sup>-L2<sup>A</sup> | +FtsN (E) | 2 µs | L32 | L2 | A3**, Repeat of 43 |
| **43c** | FtsWIQLB-N<sup>58-108</sup>-L32<sup>D</sup>-L2<sup>A</sup> | +FtsN (E) | 2 µs | L32 | L2 | A3**, Repeat of 43 |
| **53** | FtsW<sup>L198C</sup>IQLB-N<sup>58-108</sup>-L2<sup>D</sup>-L2<sup>A</sup> | L198C + MTSES, +FtsN (E) | 1 µs | L2 | L2 | A3**, 310K, with Mg* |
| **54** | FtsW<sup>S382C</sup>IQLB-N<sup>58-108</sup>-L2<sup>D</sup>-L2<sup>A</sup> | S382C + MTSES, +FtsN (E) | 1 µs | L2 | L2 | A3**, 310K, with Mg* |
| **58** | FtsW<sup>SSCC</sup>IQLB-N<sup>58-108</sup>-L2<sup>D</sup>-L2<sup>A</sup> | SSCC disulfide, +FtsN (E) | 1 µs | L2 | L2 | A3**, 310K, with Mg* |

\*All MD simulations other than 1,2,4,8 were run at 310K in the presence of 10mM Magnesium
\*\* A2 and A3 refer to Anton2 and Anton3, respectively. Simulations 42, 43, 43b, and 43c were run on Anton 3 after an 8 ns equilibration and an 8 ns preproduction run using GROMACS.

In the FtsWIQLB-L2^D^ and FtsWIQLB-L2^A^ simulations, the C55 lipid tail of L2^D^ packed tightly against the FtsW-FtsI interface between FtsW’s TM9 and FtsI’s transmembrane helix (**Fig. S4C**), while the lipid tail of L2^A^ retained considerable conformational freedom throughout the simulation (**Fig. S4D**). Furthermore, the two arginine pairs (R243-R246 and R84-R145) readily coordinated the L2 pyrophosphate group at the donor site **(Fig. S5A)** and the acceptor site **(Fig. S5B),** respectively, with at least one arginine or both from each pair maintaining electrostatic contact with the L2 pyrophosphate (defined as an N-H--O distance < 3.5 Å) for over 90% of the trajectory. These simulations confirmed that the predicted electrostatic interactions could indeed form at their respective sites and provided initial atomic coordinates for subsequent simulations.

### FtsW accommodates concurrent binding of donor and acceptor L2 molecules

Next, we assessed whether two L2 molecules could remain bound simultaneously at their respective sites. We constructed a double-docked simulation system, placing one L2 at the donor site and another at the acceptor site, and performed all-atom MD simulations on the Anton 2 supercomputer^37^ for 1 µs (FtsWIQLB-L2^D^-L2^A^-300K, **Table 2**). This configuration enabled us to examine substrate retention and binding rearrangements on the microsecond timescale.

In the FtsWIQLB-L2^D^-L2^A^-300K simulation (**Fig. 1D, E**), both L2 molecules remained bound, with their C55 lipid tails occupying the hydrophobic cavities adjacent to the donor and acceptor subsites (**Fig. 1E**, left- and right-rotated view angles), as seen in the single-L2 simulations FtsWIQLB-L2^D^ and FtsWIQLB-L2^A^. At the donor site, R243 and R246 maintained electrostatic coordination of the L2ᴰ pyrophosphate for most of the trajectory (94% and 72% of frames, respectively), with transient engagement of L2ᴰ by an additional residue, K370, in the catalytic groove during the first ∼200 ns (21% of frames, **Fig. S5C**). At the acceptor site, R145 coordinated the L2ᴬ pyrophosphate in 96% of frames (**Fig. S5C**), whereas R84, which contacted L2ᴬ in 94% of frames when the acceptor site alone was occupied (**Fig. S5B**), contacted it in only 1% of frames when both sites were occupied (**Fig. S5C**).

Notably, the acceptor-site disaccharide adopted conformations in which the GlcNAc moiety flipped down toward the catalytic groove, bringing donor and acceptor reaction atoms closer together and to D297 (**Fig. 1F**, **Supplementary Movie 2**). To quantify these conformational dynamics, we defined a catalytic triangle (**Fig. 1F**, dashed lines), in which the acceptor–donor distance is measured between the acceptor O4 atom of the hydroxyl group and the donor anomeric carbon (C1) that together form the β-1,4 glycosidic bond during the GT reaction, and the acceptor-D297 and donor-D297 distances as the separations between these atoms and the nearest carboxylate oxygen of the catalytic D297, respectively **(Table 3).** We expect GTase catalysis to require bond-forming acceptor-donor distances of < 4 Å and proximity between D297 and the acceptor O4 atom (which must get deprotonated)^38^, whereas a short donor-D297 distance is not necessary in the pre-catalytic geometry.

**Table 3.** Summary statistics for pairwise catalytic distances.

| Sim | Mean Donor-Acceptor Distance (Å) | Mean D297-Acceptor Distance (Å) | Mean D297-Donor Distance (Å) |
|---|---|---|---|
| **WT (300K)** | 14.4 | 9 | 10.8 |
| **WT (A)** | 35.2 | 29.2 | 11.7 |
| **WT (B)** | 21.8 | 18.6 | 11.0 |
| **E289G (A)** | 14.8 | 10.3 | 9.5 |
| **E289G (B)** | 18.4 | 15 | 11.3 |
| **+FtsN (0-1 µs)** | 14.6 | 13.0 | 9.7 |
| **+FtsN (1-4 µs)** | 16.2 | 14.5 | 9.3 |
| **+FtsN, full** | 15.8 | 14.1 | 9.4 |
| **+FtsN, L4** | 13.1 | 10.8 | 8.9 |
| **+FtsN, L32 (A)** | 11.5 | 9.6 | 9.0 |
| **+FtsN, L32 (B)** | 13.1 | 10.0 | 8.6 |
| **+FtsN, L32 (C)** | 13.7 | 12.3 | 9.1 |

Using this catalytic triangle metric, we observed that the acceptor-donor distance decreased from ∼18 Å to ∼12 Å within the first 400 ns and then remained stable for the remainder of the trajectory (**Fig. 1G**, grey). The Acceptor-D297 **(Fig. 1G**, cyan) and the Donor – D297 **(Fig. 1G**, green) distances decreased similarly to ∼10 Å after 400 ns. Although these reductions did not yet reach bond-forming distances, they are consistent with these sites representing the physiological binding pockets and with a model in which dual substrate binding precedes additional rearrangements required to achieve a true pre-catalytic state.

To test whether these conformational changes also occur at physiological temperature and in the presence of Mg²⁺, which is required for the GTase activity of *Streptococcus thermophilus* FtsW but has not been demonstrated for gram-negative FtsW^4^, we performed two independent 1 µs replicas at 37 °C (FtsWIQLB-L2^D^-L2^A^-310K A and B; **Table 2**). In both replicas, the same conserved residues (R84, R145, R243, R246, and K370) formed sustained electrostatic contacts with the bound substrates (**Fig. S5D**), consistent with a pre-formed binding complex before activation. However, acceptor-site engagement was markedly less stable at 310 K, with frequent partial or complete disengagement of the acceptor disaccharide from the catalytic groove; accordingly, the mean acceptor-D297 distance increased to 29.2 Å (replicate A) and 18.6 Å (replicate B), and the mean donor– acceptor distance increased to 35.2 Å (A) and 21.8 Å (B), respectively (**Table 3**). In contrast, the donor-site geometry remained comparatively constrained under all WT conditions (mean D297–donor distance: 10.8 Å at 300 K; 11.7 Å and 11.0 Å at 310 K; **Table 3**), indicating that the donor site more tightly restricts substrate positioning than the acceptor site. Together, these results support stable dual-substrate binding at the assigned sites but also indicate that robust acceptor-site binding may require additional stabilizing factors under physiological conditions.

### Metadynamics reveals that L2 can access the donor site through the FtsW TM6-TM7 opening

While our MD simulations support the assignment of donor and acceptor sites, they do not explain how an L2 molecule accesses the donor site, which is buried within FtsW (**Fig. 1D**). Although FtsW’s TM9 and FtsI’s transmembrane helix form a cavity that opens to the membrane, direct diffusion of an L2 molecule through this opening is blocked because the AH1 in FtsW’s ECL4 sterically occludes the L2 sugar headgroup (**Fig. S4B**). In contrast, in some simulations FtsW’s TM6 and TM7 transiently widened and opened to the membrane (**Supplementary Movie 1**), suggesting this interface could act as a gated entry pathway that allows an L2 molecule to reach the donor site from the membrane-exposed outer surface of FtsW.

To test this possibility, we quantified the conformational state of this putative “outer gate” by measuring the Cα–Cα distance between L198 (TM6) and L236 (TM7) in apo simulations of FtsWIQLB and in those formed by a superfission variant, FtsW-E289G, which we previously performed^20^. This distance effectively represents the widest gap between the two helices. In the FtsWIQLB apo simulation, this distance is typically ∼10 Å, consistent with a tightly packed TM6–TM7 interface (**Fig. S6A**, left), whereas the superfission FtsW-E289G apo complex shifted the distance to larger values (∼13–15 Å; **Fig. S6A,** middle). An open TM6–TM7 interface was also evident in the RodA cryoEM structure^8^ (PDB: 8TJ3), where the corresponding helices (L160–L202) are 22 Å apart, compared with ∼10 Å in WT FtsWIQLB (**Fig. S6A**, right). Together, these observations supported a model in which the TM6–TM7 interface may indeed function as a dynamic gate that interconverts between closed and open conformations, where an “open” gate is operationally defined as a conformation that permits L2 to enter or exit the donor cavity.

To probe whether L2 can access the donor cavity through this outer gate, we docked an L2 molecule at the TM6-TM7 interface, just outside the donor site (termed D’, **Fig. 1C**), and equilibrated the system for 52 ns (FtsWIQLB-L2^D’^, **Table 2**). During equilibration, the L2 pyrophosphate formed stable electrostatic interactions with FtsW’s R243 **(Fig. S5E),** while the C55 tail packed against the membrane-accessible surface between FtsW TM6 and TM7 (**Fig. S6B**, left). In this ensemble, we observed limited tail penetration into the TM6–TM7 interface; sidechain contacts between TM6 and TM7 sterically blocked full tail entry into the donor cavity.

Because opening of membrane-embedded helices is expected to be a rare event on equilibrium MD timescales, we next applied well-tempered metadynamics, an enhanced-sampling method^39^ using the TM6–TM7 helix–helix separation (centers-of-mass distance) and the position of the C55 tail relative to FtsW TM8 and the FtsI TM as collective variables (WT-MetaD, FtsWIQLB-L2^D’/D^, Methods). Metadynamics is an enhanced-sampling MD method that accelerates conformational sampling by applying Gaussian energetic biases along selected collective variables, thereby driving the system toward otherwise inaccessible conformational states and transition pathways^39,40^.

In the metadynamics trajectory, we observed that L2 moved from D’ into D as the C55 tail threaded between TM6 and TM7, accompanied by new electrostatic contacts with FtsW K370 and R246 (**Fig. S6B**, middle, **Supplementary Movie 3**). The lipid tail fully transitioned from the outer to the inner side of the gate by ∼60 ns (**Fig. S6B**, right), followed by partial gate closure behind the tail at ∼100 ns and complete cavity closure by ∼150 ns **(Fig. S6C,** top). Correspondingly, the L198-L236 distance increased to a maximum of 18.1 Å during the 0-60 ns transition window, likely representing an open conformation of FtsW that permits L2 entry (**Fig. S6C**, bottom). Based on post-processing of the metadynamics data, we defined the lowest-energy path (Methods) and estimated a free energy barrier of ∼5 kcal/mol for opening the TM6-TM7 outer gate (**Fig. S6D**), consistent with reported values for membrane-helix opening transitions and therefore physiologically feasible.

Furthermore, during the D’→D transition, 95% of the simulation trajectory (5^th^ percentile) showed a distance ≥12.7 Å between L198 and L236 (**Fig. S6E**), indicating that the outer gate remained predominantly open. We therefore used L198–L236 separation as a quantitative readout of TM6–TM7 gating and adopted ∼13 Å as the operational threshold for an “open” outer gate (**Fig. S6F**). Based on this threshold, the TM6–TM7 gate of the FtsW^E289G^ apo complex adopted a significantly more open state than the WT FtsW apo complex (**Fig. S6G**), likely increasing the probability of an L2 molecule entering the donor site from the membrane. However, a persistently open outer gate also favors a bound L2 escape from the binding pocket, implying that the superfission mutation is more likely to accelerate substrate turnover than simply increase donor-site binding affinity. In summary, the metadynamics results suggested that L2 entry into the donor site is energetically feasible with relatively modest conformational change in FtsW TM6-TM7 and is facilitated by the same residues (R243, R246, K370) that stabilize FtsWIQLB-L2^D^ binding.

### Chemical blockade of the donor site of FtsW inhibits cell division

The simulations above indicate that the FtsW donor site relies on a combination of electrostatic coordination and precise steric gating to achieve stable, specific L2 binding. We reasoned that if these interactions are essential for FtsW function, disrupting them should impair FtsW activity and inhibit cell growth. To test this idea, we introduced an FtsW L198C mutation, which should be covalently modified by the cysteine-reactive reagent MTSES (2-sulfonatoethyl methanethiosulfonate) in a solvent-accessible periplasmic environment^42^. In our simulations, L198 packs against the donor L2 and lies in close proximity to the conserved basic residues (R243, R246, K370). Therefore, introducing a bulky, negatively charged MTSES adduct at this position could block inward movement of the L2 donor and compete for these key electrostatic contacts.

In the absence of MTSES, cells expressing FtsW(L198C) in an FtsW-depletion background (AP852, **Supplementary Table 1; plasmids and primers, Supplementary Tables 2and 3)** showed mostly rod-shaped cell morphology, but a small fraction of cells was elongated (6.6% of cells, range 5.2–7.8% across replicates, versus 0.6%, range 0–1.9%, for cells expressing WT FtsW in M9 medium; elongated cells are defined in Methods; **Fig. 1H**, top right; **Fig. S7A**, 1st and 4th groups, n = 3 biological replicates, 169 cells; Supplementary **Table 4**), suggesting partial function of FtsW(L198C). The addition of MTSES to FtsW(L198C), but not wild-type FtsW (**Fig. S7A**, 2^nd^ and 5^th^ groups), blocked cell division in most cells and produced severely filamented cells (**Fig. 1H**, middle right, **Fig. S7A**, 5th group; 81.6% of cells elongated, range 73.0–90.6%; n = 3 biological replicates, 143 cells; Supplementary **Table 4**), approaching the proportion seen in cells lacking FtsW entirely (**Fig. 1H**, bottom left, Fig. S7A, 6th group; 92.6%, range 86.2–97.3%; n = 3, 113 cells) or a previous *ftsW*(I302C) mutant upon MTSES treatment^22^. As *wildtype* (*wt*) cells grew normally in the presence of MTSES (**Fig. 1H**, middle left), this result suggested that the filamentous phenotype was FtsW(L198C)-specific rather than a general sensitivity of the cell to MTSES.

A filamentous phenotype alone cannot distinguish direct inhibition of FtsW-mediated sPG synthesis from indirect effects such as protein destabilization or defective divisome assembly. We therefore asked whether the engineered FtsW(L198C) variant was expressed normally and properly recruited to the divisome under the inhibitory condition. We constructed an FtsW(L198C)-Halo fusion protein expressed from a plasmid in the presence of the native chromosomal *ftsW* gene (AP896, **Supplementary Table 1),** labeled it with a saturating level of Halo-JF646 dye, and observed that it localized to the midcell in the presence of MTSES (**Fig. S8A**), with the ensemble fluorescence level at ∼81% of the WT FtsW-Halo signal (**Fig. S8B**). When further induced at 100 µM IPTG, FtsW(L198C)-Halo produced a dominant-negative, filamentous phenotype (**Fig. S9**); 35.7% of cells were elongated without MTSES (range 21.8–50.0%) and 60.1% with MTSES (range 53.1–68.5%), compared with 0% for the matched FtsW-Halo control (n = 3 biological replicates; 143 and 161 cells, respectively; **Fig. S9A,C**). We noted that direct probing of MTSES modification at L198C (as well as other mutations below) using mass spectrometry on HaloTag-enriched FtsW-Halo from cells grown with and without MTSES was inconclusive, as we did not detect any peptide containing the L198 position, either native or modified, which was likely due to challenges working with lowly expressed membrane proteins (Methods, **Fig. S10).** Nevertheless, our results (MTSES-specific defects, similar midcell localization, and a dominant-negative phenotype upon induction) strongly suggested that the observed inhibitory effect was specifically due to modification at L198C, rather than to destabilization or degradation of the mutant protein.

Finally, to further confirm that MTSES-modified FtsW(198C) exhibited the expected defective sPG synthesis activity, we performed SMT of FtsW(L198C)-Halo expressed in the presence of a chromosomal FtsW(L198C) copy (AP887, **Supplementary Table 1; Fig. S11).** As shown in **Fig. S11B**, without MTSES, FtsW(L198C)-Halo partitioned into stationary (∼70%), slow-moving (13%), and fast-moving (18%) populations (N = 313 segments). In contrast, MTSES treatment nearly eliminated the active, slow-moving population (2%) and increased the inactive, fast-moving population to 33% (N = 314 segments, **Supplementary Table 5).** Together, these results strongly suggested that chemical blockade of L198 at the donor site inhibits FtsW’s function.

### The FtsWIQLB complex adopts a self-inhibited conformational state

While our MD simulations showed that L2 can access the donor site and approach the acceptor in FtsW’s catalytic groove, and the chemical blockade experiment supported the assigned donor binding site, the pairwise distances between the L2 donor, acceptor, and FtsW D297 did not reach a bond-forming or bond-breaking range (**Fig. 1G**). Across all WT simulations, positively charged residues such as R145 and R243 coordinated the L2 pyrophosphate groups and restricted them to their initial positions along the catalytic groove, keeping them away from D297. These observations suggested that the WT FtsWIQLB complex may exist in a self-inhibited state (**Fig. 2A**, top).

**Figure 2.**
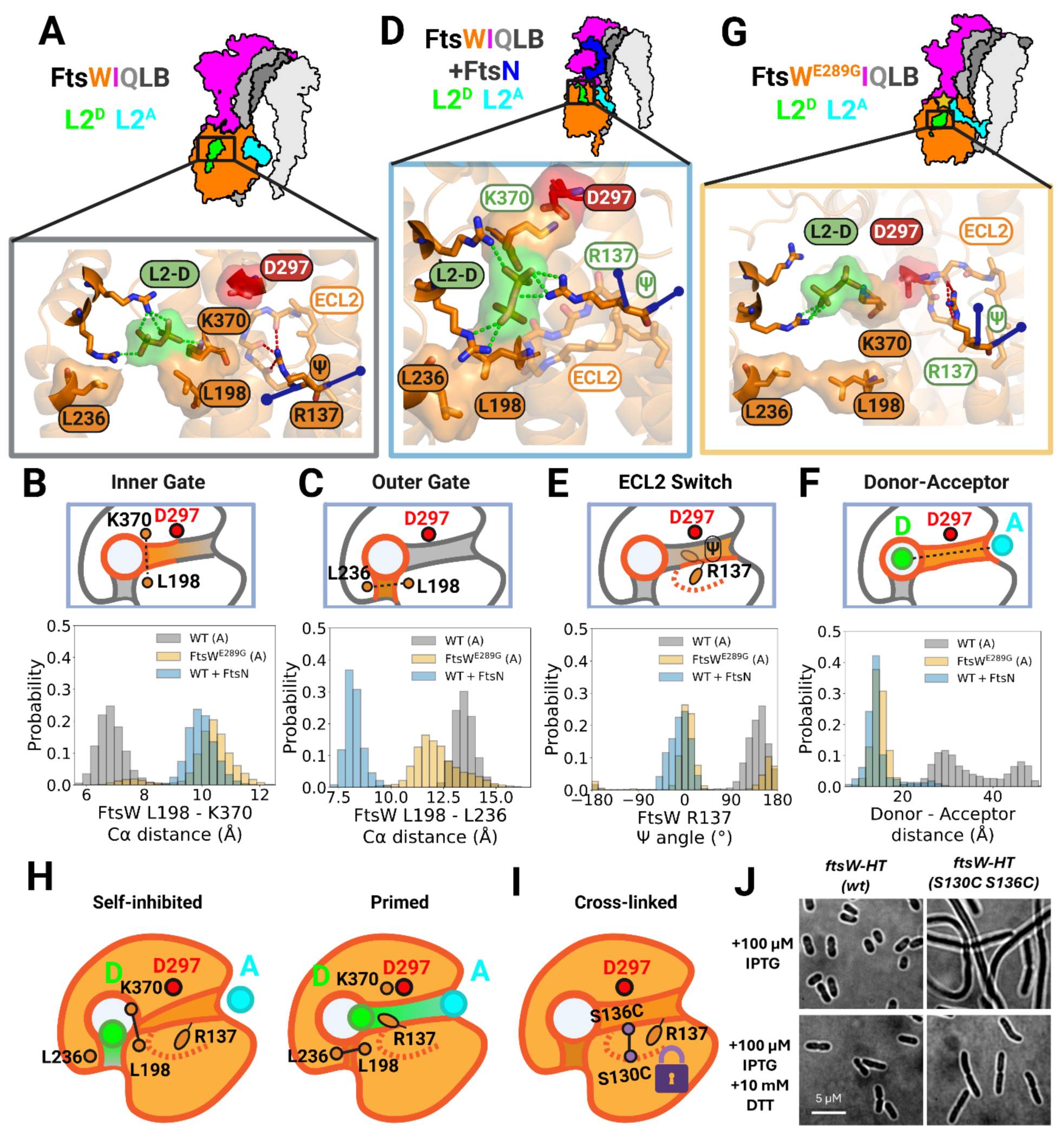
Activation remodels FtsW gating and ECL2 dynamics to prime donor lipid II site for catalysis. **(A, D, G)** Representative structural models illustrating three functional ensembles (top, cartoons): self-inhibited WT FtsWIQLB **(A)**, primed FtsWIQLB–FtsN **(D)**, and uninhibited superfission FtsW^E289G^IQLB **(G)**. Bottom, close-up structural views show donor lipid II (D–L2; green) positioned relative to the catalytic residue D297 (red) and key gating elements (TM6 L198, TM7 L236, ECL5 K370) and the ECL2 residue R137. The FtsW backbone is shown as ribbon; key FtsW residues and the donor lipid II pyrophosphate groups are shown as sticks; steric gating elements near the donor cavity are rendered as surfaces. Elements engaged with the L2^D^ are labelled with white and green labels. **(B)** Inner-gate metric. Top, schematic defining the donor-side inner gate as the separation between L198 (TM6) and K370 (ECL5) adjacent to the catalytic groove. Bottom, probability distributions of the L198–K370 Cα–Cα distance for WT (gray), FtsN-bound (blue), and E289G (yellow) simulations. **(C)** Outer-gate metric. Top, schematic defining the TM6–TM7 outer gate as the separation between L198 (TM6) and L236 (TM7) at the donor-entry interface. Bottom, probability distributions of the L198–L236 Cα–Cα distance for the same ensembles as **(B)**. **(E)** ECL2 switch metric. Top, schematic defining the ECL2 activation readout as the backbone ψ dihedral of R137. Bottom, probability distributions of the unwrapped R137 ψ angle for the same ensembles as **(B)**. **(F)** Donor–acceptor proximity in lipid II–bound simulations. Top, schematic defining the donor– acceptor distance between the donor anomeric carbon (C1) and the acceptor O4 atom. Bottom, distributions of donor–acceptor distances for the same ensembles as **(B)**. **(H)** Summary model for coupled gating and ECL2 switching. In the self-inhibited ensemble (left), the outer gate samples open states, and the inner gate remains closed, with R137 predominantly buried in an inactive ψ configuration. FtsN-bound or E289G systems experience the inner-gate opening and shifts of ECL2 toward an active R137 ψ state, yielding a primed donor-site architecture (right). **(I)** Summary model displaying the design of an ECL2 disulfide crosslink to restrict ECL2 dynamics without mutating R137. Two ECL2 residues (S130 and S136) were mutated to cysteines to permit disulfide formation. **(J)** Brightfield micrographs comparing morphological effects of cells expressing FtsW-Halo (left column, strain AP183), or FtsW(SSCC)-Halo (right column AP760) as sole copies of *ftsW* under control of an IPTG inducible promoter. Cells were imaged from LB plates induced with 100 µM IPTG under non-reducing (−DTT) and reducing (+DTT) conditions. Micrographs are representative of three independent biological replicates. HT: Halo tag.

To identify structural features of the WT complex that might correspond to a self-inhibited conformation, we examined residue conformations in and near the binding sites and catalytic groove. Interestingly, in WT 310K simulations, K370 in ECL5 consistently coordinated the pyrophosphate group of the donor L2 molecule (**Fig. S12A**), but it simultaneously also packed tightly against TM6, forming a physical barrier between the L2 headgroup and the catalytic residue D297 (**Fig. 2A**, bottom). Our previous MD simulation of the WT apo complex also implicated K370 as a residue capable of blocking the catalytic groove^20^. Although K370 has modest sequence conservation (63%), its positive charge and strategic position between R246 and D297 suggest that it may play an important role in regulating donor-site access to D297.

The close packing of K370 against TM6, which we termed the “inner gate”, appeared inhibitory, as it prevented the donor L2 from approaching either D297 or the acceptor L2 molecule (**Fig. 2B**, top). To quantify this packing, we measured the mean Cα–Cα distance between L198, immediately adjacent to TM6, and K370 at 6.86 ± 0.007 Å **(Fig. 2B**, bottom, grey bars), consistent with a tightly closed TM6-ECL5 interface that blocks access of the L2 donor to the catalytic groove. Interestingly, the outer gate between TM6 and TM7 remained largely open, with a mean L198 - L236 Cα–Cα distance at 13.5 ± 0.007 Å (**Fig. 2C**, grey bars). This open outer-gate conformation could facilitate L2 entry into the donor site but would also allow a bound L2 to escape without steric hindrance. Thus, the donor L2 molecule remains poorly positioned for the glycosyltransfer reaction, supporting the view that the WT FtsWIQLB complex adopts a self-inhibited state awaiting activation.

### FtsN^E^ binding induces a primed conformation of FtsW poised for catalysis

In most *γ*-proteobacteria, FtsW requires activation by FtsN^17,43^. Therefore, we investigated how FtsN impacts substrate-bound FtsW conformation. We utilized our previously generated FtsWIQLBN^58-108^ apo simulation^20^, in which the essential domain of FtsN^58-108^ is proposed to bind FtsI and FtsL at the AWI region in the periplasm. Previous studies have shown that FtsN’s essential domain is necessary and sufficient to activate FtsW^19,20,41,42,43^. We docked two L2 molecules into a trajectory snapshot from the FtsWIQLB-N^58-108^ apo simulation and performed a 4 µs all-atom simulation (FtsWIQLB-N^58-108^-L2^D^-L2^A^ at 310K with 10 mM Mg^2+^, **Supplementary Table 2,** Methods). Remarkably, the simulation revealed several critical rearrangements in the interactions between FtsW and bound L2 molecules (**Supplementary Movie 4**, a morph between the final snapshots of the WT-A and FtsN-bound simulations, Sim# 14 and 17 in **Table 2**).

First, K370 in FtsW ECL5 formed a persistent salt bridge with D297^20^ (**Fig. S12B**), widening the gap between K370 and L198 to 10.0 ± 0.007 Å (**Fig. 2B**, bottom, compare grey and blue bars). In the ligand-bound context, this opening effectively cleared a pathway for the donor L2 to enter the catalytic groove (**Fig. 2D**, bottom). Most interestingly, the K370-L2 Donor interaction was mutually exclusive with the K370-D297 interaction throughout the simulation (**Fig. S12C**), suggesting a dynamic switching mechanism. In our previous FtsWIQLB-N^58-108^ apo simulation, we also observed similar K370-D297 salt-bridge contact^20^.

Second, in the presence of FtsN^58-108^, the outer gate formed by TM6 and TM7 remained closed throughout the entire 4 µs-long trajectory (**Fig. 2D**, bottom), with a mean L198-L236 distance of 8.3 ± 0.007 Å (**Fig. 2C**, bottom, blue bars). This stable, closed conformation prevented the donor lipid tail from escaping the cavity, in sharp contrast to the predominantly open outer gate observed in the WT L2-bound FtsWIQLB complex.

Finally, we observed a pronounced backbone rearrangement in FtsW ECL2 (**Fig. 2D**, bottom, **Fig. S13).** ECL2 is part of an extracellular-loop surface conserved among SEDS proteins^6,7^. In particular, R137 in ECL2, a basic residue that is 78% conserved (**Table 1**) and whose R137H substitution leads to a dominant-negative phenotype^33^, flipped into the catalytic groove, where it directly contacted the donor L2 pyrophosphate group (**Fig. 2D**, bottom). In this conformation, R137 was no longer buried or forming polar contacts with the ECL2 backbone carbonyl oxygens, as in the WT complexes (buried surface area BSA ≈ 150 Å^2^ *vs* ≈ 300 Å^2^ in WT, **Fig. S13A-B).** Instead, it consistently adopted a Ψ angle of ∼ 0° compared to ∼ 150° in WT complexes (**Fig. 2E**, bottom, compare blue and gray bars). In this rearranged state, R137 is the sole conserved, positively charged residue positioned to bridge the L2 donor and acceptor molecules, with its Cα within 15 Å of both reactive atoms at the end of the FtsWIQLB-N^58-108^ -L2^D^-L2^A^ simulation.

To confirm that the observed ECL2 rearrangement arises from FtsW activation rather than from bias in the initial L2 docking coordinates, we analyzed previously reported apo MD trajectories initiated from AlphaFold 2 models^20^. We quantified the backbone dynamics over time by defining an “active” ECL2 state as any snapshot in which the Ψ angle of R137 was between -90° and 90° **(Fig. S14A).** We found that R137 began in the active conformation in all predicted structures but relaxed to an inactive state within the first 500 ns across all FtsWIQLB complexes examined (**Fig. S14B**). By contrast, the complex containing FtsN^58-108^ retained the active configuration throughout the 1µs trajectory (**Fig. S14C**). This consistent pattern across simulations supports a self-inhibited ECL2 switch that is relieved upon binding of FtsN.

As a result of these rearrangements, the donor and acceptor L2 molecules moved substantially closer to each other in the presence of FtsN^58-108^, with an average separation of 15.8 Å compared to 35.2 Å and 21.8 Å in the WT 310K-A and 310K-B simulations, respectively (**Fig. 2F, bottom**, blue bars; Table 3). Because these structural changes were absent from simulation trajectories of the WT FtsWIQLB complex with L2 bound at both the putative donor and acceptor sites (FtsWIQLB-L2^D^-L2^A^-310K), they suggested that the binding of the activator protein FtsN^58-108^ to the complex triggers conformational switching in key basic residues, driving the L2 donor deeper into the catalytic groove and positioning the pyrophosphate leaving group for the GT reaction. Collectively, we defined this set of rearrangements as a primed conformation of FtsW poised for catalysis (**Fig. 2H**, right), in contrast to the self-inhibited conformation observed in the WT FtsWIQLB complex (**Fig. 2H**, left).

### FtsW-E289G superfission complex adopts an intermediate activated conformation

To determine whether the FtsN^58-108^-induced conformational changes in FtsW reflect a general activation mechanism, we next examined an FtsW superfission variant that bypasses the requirement for FtsN activation and exhibits hyperactive sPG synthesis^20,22^. We performed a 1 µs atomistic simulation (310K with 10 mM Mg^2+^, **Table 2**) of a complex containing the superfission variant FtsW E289G (FtsW^E289G^IQLB-L2^D^-L2^A^, **Fig. 2G, Supplementary Movie 5)**. E289 resides in ECL4 and is adjacent to the central cavity that harbors the catalytic residue D297. In our prior simulations, it hydrogen-bonded with the periplasmic anchor loop of FtsI and constituted a regulatory interface with FtsI^20^.

In the FtsW^E289G^IQLB-L2^D^-L2^A^ simulation, the donor-side inner gate (L198–K370) widened to 10.2 ± 0.015 Å (**Fig. 2B**, yellow bars), closely resembling the FtsN-bound complex (blue bars) but differing from the WT FtsWIQLB complex (**Fig. 2B**, grey bars). The TM6–TM7 outer gate (L198–L236) also showed reduced separations (12.3 ± 0.017 Å; **Fig. 2C**, yellow bars) relative to WT FtsWIQLB, which are intermediate between those of WT FtsWIQLB and the FtsN-bound complexes (**Fig. 2C**, blue and grey bars). Moreover, we observed extensive conformational rearrangements in FtsW involving ECL2, the ECL4 segment between AH1 and AH2, and the periplasmic region of TM1 (**Fig. S15A**). The R137 backbone Ψ angle displayed a clear two-state distribution (**Fig. 2E**, yellow bars), initially resembling the WT FtsWIQLB angles at ∼150° and later transitioning to the ∼0/360° conformation characteristic of the FtsWIQLB-N^58-108^-L2^D^-L2^A^ complex (**Fig. S15B**, top). This transition allowed R137 to flip out of ECL2 (**Fig. S13C**, R137 – ECL2 BSA ∼= 200 Å^2^) and approach D297, although we did not observe the same direct contact between R137 and the donor L2 pyrophosphate (**Fig. S13D**). Interestingly, FtsW R73, which formed a strong and persistent salt-bridge with E289 in WT FtsWIQLB simulations (**Fig. S15C**), instead engaged more frequently with the acceptor peptide stem in E289G simulations, consistent with the loss of E289 **(Fig. S15D).** As a result, the donor–acceptor distance was, on average, shorter than that in WT FtsWIQLB (A: 14.8 Å; **Fig. 2F**, bottom yellow bars, **Table 3**), and strikingly similar to that of the FtsN-bound complex.

To confirm that these conformational changes were reproducible, we performed a second FtsW^E289G^IQLB-L2^D^-L2^A^ simulation using randomized initial velocities from the same starting coordinates (**Supplementary Movie 6**). This replicate similarly reproduced shifted outer- and inner-gate geometries and consistently shorter donor-acceptor distances compared with WT FtsWIQLB (**Table 3**). However, R137 in this replicate did not undergo a sustained ψ transition and instead showed only transient fluctuations before returning to its initial conformation (**Fig. S15B**, bottom). Despite this difference, both FtsW^E289G^IQLB-L2^D^-L2^A^ simulations showed increased flexibility of FtsW around the donor site and a persistently enlarged gap between the inner gate L198-K370, regardless of whether a strong K370-D297 salt bridge formed (**Fig. S12C**). Notably, in both replicates, the donor L2 disaccharide of L2^D^ flipped directly into the catalytic groove (**Supplementary Movies 5 and 6**), with its nonreducing end occupying the solvent-exposed space between ECL2 and ECL5. Although this orientation is unlikely to support catalysis (the donor can only react at the reducing end), it indicates that E289G leaves the inner gate sufficiently open to accommodate the L2^D^ head group.

Together, these simulations demonstrated that both FtsN and the FtsW^E289G^ superfission variant open the K370 inner gate, thereby removing the principal steric barrier characteristic of the self-inhibited architecture of FtsWIQLB (**Fig. 2H**, left). However, the donor site in the FtsW^E289G^ complex did not attain the fully closed outer gate configuration or the consistently active R137 backbone angle observed in the primed FtsN-bound complex (**Fig. 2H**, right), suggesting that the E289G variant may represent an intermediate, or partially activated, conformational state between the self-inhibited and primed conformations.

### FtsW(K370C) exhibits a strongly dominant-negative phenotype

Given the vital role of K370 as part of the inner gate in controlling the donor’s access to the catalytic groove (**Fig. 2B**), we generated an FtsW(K370C) variant to probe the effect of its modification by MTSES. However, the FtsW(K370C) variant did not support growth in an FtsW depletion background even in the absence of MTSES, and we were unable to culture the depletion strain for imaging, suggesting that the mutation itself is poorly tolerated. We therefore characterized an FtsW(K370C)-Halo fusion in the presence of a *wt* chromosomal *ftsW* copy (AP895, **Supplementary Table 1).** When induced at 100 µM IPTG, FtsW(K370C)-Halo caused severe filamentation despite the presence of WT FtsW and the absence of MTSES (100% of cells elongated in all three replicates, versus 0% for the FtsW-Halo control; 74 ± 5 µm; n = 3 biological replicates, 29 cells; **Fig. S9B,C**), consistent with a strong dominant-negative effect and supporting the functional importance of the K370-containing inner-gate region identified in the simulations.

### Constraining FtsW ECL2 mobility by an engineered disulfide bond blocks cell division

Our results indicate that R137 in ECL2 plays a critical role in FtsW activation. The functional significance of R137 has been demonstrated previously, as a single R137H substitution caused a dominant-negative, lethal phenotype^33^. To examine the contribution of ECL2 dynamics without directly mutating R137, we employed a previously established *in vivo* cysteine crosslinking approach^47^ to restrict ECL2’s mobility and assess its functional impact.

We selected S130 and S136 in ECL2 for cysteine mutagenesis (FtsW(S130C S136C), referred to as FtsW(SSCC) below) based on their low sequence conservation (12% conservation, **Fig. S3)** and spatial proximity in the apo FtsWIQLB complex (**Fig. 2I, Fig. S16A).** In most simulation snapshots, the Cα-Cα distance between the two residues was < 8 Å, which would enable disulfide bond formation in the oxidative periplasmic environment if both were mutated to cysteine^48,49^ (**Fig. S16B**). In the presence of a reducing agent such as DTT (Dithiothreitol) in the growth medium, however, the reduced, non-crosslinked form would be favored over the disulfide-bonded state^42^.

As shown in **Fig. 2J**, the double mutant FtsW(SSCC)-Halo, expressed in a *ftsW* depletion background, was mostly rescued by DTT: 70.0% of cells had normal rod morphology under the reducing condition (**Fig. 2J**, bottom right), against 92.0% for the control expressing WT FtsW-Halo, with the double mutant retaining a 1.36 fold increase in median (95% interval 1.03–1.82; Fig. S7B, Supplementary **Table 6**). However, in the absence of DTT, FtsW(SSCC)-Halo caused a severe filamentous phenotype with 77.2% of cells elongated (range 62.0–94.0%; cell length 12 ± 8 µm; n = 3 biological replicates, 149 cells; **Fig. 2J**, top right, **Fig. S7B),** and cells failed to divide. FtsW(SSCC)-Halo also retained midcell localization under non-reducing conditions in merodiploid cells co-expressing chromosomal WT FtsW (AP893, **Supplementary Table 1; Fig. S8A),** with ensemble JF646 fluorescence at ∼ 100% of the WT FtsW-Halo signal (**Fig. S8B**). When induced with 100 µM IPTG in the same merodiploid background in the absence of DTT, FtsW(SSCC)-Halo produced a modest elongated phenotype (12.9% of cells elongated, range 3.0–27.9%, versus 0% for the FtsW-Halo control; n = 3 biological replicates, 145 cells; **Fig. S9B,C**), consistent with the engineered disulfide forming spontaneously in the oxidizing periplasm and exerting a dominant-negative effect on cell division. In a separate experiment in EZ rich defined medium (EZRDM) at 37 °C without DTT, neither single mutant filamented (1.9% and 13.2% of cells elongated, ranges 1.3–2.4% and 1.9–30.6%, versus 9.9%, range 0–25.9%, for FtsW-Halo), whereas 95.0% of double-mutant cells were elongated (range 89.1–100%; 41 ± 12 µm; n = 3 biological replicates, 217 cells; **Fig. S16D**, Supplementary **Table 7**). Although direct verification of the disulfide bond formation of HaloTag-enriched FtsW(SSCC)-Halo by mass spectrometry was inconclusive (**Fig. S10**, Supplementary **Table 8**), these results showed that the double mutant’s filamentous phenotype was not caused by either substitution alone, or by the simultaneous presence of both cysteine substitutions (**Fig. S16),** but most likely by their disulfide-bond formation. Therefore, we concluded that ECL2 dynamics are essential for FtsW activity.

### FtsN activates an acceptor clamp to stabilize acceptor binding

In FtsWIQLB–FtsN^58–108^ simulations, we observed that FtsN stabilized the binding of not only the donor but also the acceptor relative to WT FtsWIQLB trajectories (**Fig. 3A,B**, **Supplementary Movie 7** for WT-A and **Supplementary Movie 8** for FtsN-bound), with the distribution of donor-acceptor separations forming a distinct population centered on ∼16 Å compared to the ∼20 Å population in the WT-B replicate (**Fig. S17A**, blue bars). Because productive catalysis requires full entry of the incoming acceptor L2 disaccharide into the catalytic groove, we quantified the buried surface area (BSA) between FtsW and the acceptor disaccharide (**Fig. 3C**, **Fig. S17B,C).** In WT simulations, acceptor binding was intrinsically intermittent, primarily maintained by electrostatic contacts between the pyrophosphate group and R84/R145 and by occasional nonspecific packing of the disaccharide against a hydrophobic patch on ECL1 (**Fig. S17B**, residues L74–K83), resulting in either complete disengagement (near zero BSA, WT-A; **Fig. 3C**, gray bars) or only low-to-moderate interfacial sampling (WT-B; **Fig. S17D).** By contrast, in the FtsWIQLB–FtsN^58–108^ complex (**Fig. 3B**), the acceptor entered deeply into the catalytic groove, with the BSA distribution shifted toward substantially higher values (mean at 447 ± 4 Å^2^ with a sustained population near ∼600 Å^2^ (**Fig. 3C**, blue bars; **Fig. S17C, E**).

**Figure 3.**
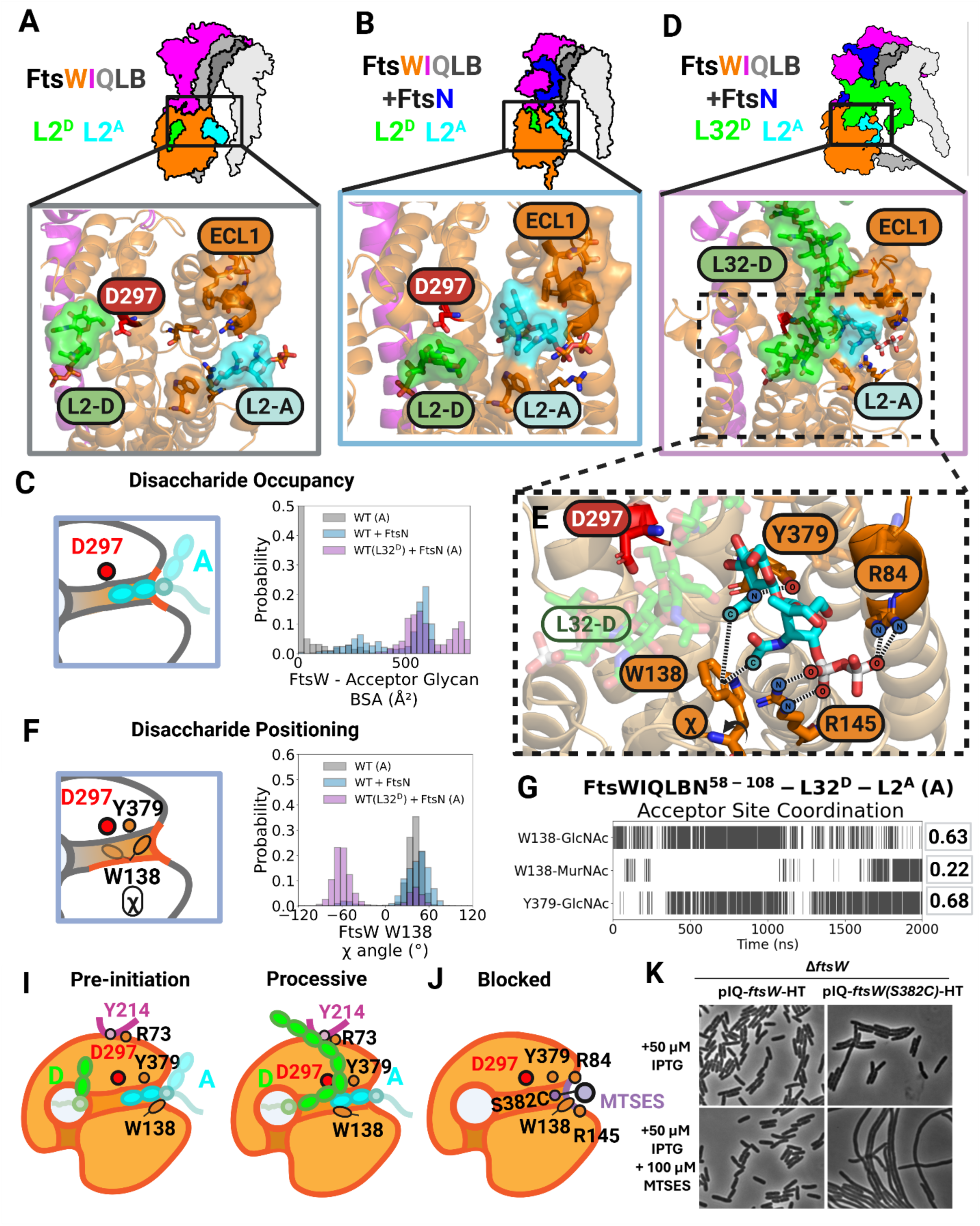
FtsN and donor chain length stabilize acceptor-site binding in FtsW. **(A–B, D)** Representative structural models illustrating three functional ensembles (top, cartoons): **(A)** self-inhibited FtsWIQLB with lipid II in donor and acceptor sites (L2^D^/L2^A^), **(B)** primed FtsWIQLB– FtsN with L2^D^/L2^A^, and **(D)** processive FtsWIQLB–FtsN with an extended donor glycan (L32^D^) and lipid II in the acceptor site (L2^A^). Representative acceptor-site snapshots from MD simulations (bottom). Key acceptor-site features are indicated, including ECL1 and catalytic D297; donor (green) and acceptor (cyan) substrates are labeled where shown. **(C)** Buried surface area (BSA) between FtsW and the acceptor glycan for the indicated trajectories (schematic, left; distributions, right), reporting acceptor-site engagement. **(F)** Acceptor disaccharide positioning metric based on the FtsW W138 side-chain χ angle (schematic, left; distributions, right), which reports formation of a packed acceptor configuration. **(E, G)** Acceptor-site coordination in the processive state. **(E)** Close-up view highlighting a set of contacts that form between the acceptor GlcNAc to FtsW residues (W138, Y379, R84, and R145). **(G)** Binarized contact occupancy over time for the indicated interactions; values at right report the fraction of frames satisfying the contact criterion (distance < 3.5 Å). **(I)** Schematics for acceptor-site engagement during initiation versus processive synthesis: in a pre-initiation state the acceptor glycan is weakly associated with the catalytic groove, whereas an extended donor chain stabilizes a tightly packed acceptor positioned near D297. **(J)** Schematic for chemical occlusion of the acceptor-site cavity: covalent modification at S382C (MTSES) blocks the acceptor interface near W138/Y379 and prevents stable acceptor binding. **(K)** Phase-contrast micrographs of cells expressing FtsW-Halo (AP183) or FtsW(S382C)-Halo (AP766) as the sole copy of *ftsW* from an IPTG-inducible promoter in EZRDM at 37 °C, with or without MTSES as indicated. Images are representative of three independent biological replicates.

Two highly conserved aromatic residues, W138 in ECL2 and Y379 in ECL5 (99% and 75% conserved, respectively, **Fig. S3),** which produced dominant-negative phenotypes when mutated to alanine^33^ and flanked the acceptor disaccharide from opposite sides, engaged the N-acetyl groups on the GlcNAc-MurNAc sugars (**Fig. S18A–C**). These interactions appeared to play a crucial role in stabilizing acceptor binding. Specifically, we observed frequent hydrogen bonding between Y379 and the acceptor GlcNAc acetyl group (37% occupancy), whereas this interaction was essentially absent in WT simulations (**Fig. S18A**). In parallel, W138 engaged in methyl–aromatic packing with the acceptor acetyl group on GlcNAc (**Fig. S18B**) or MurNAc (**Fig. S18C**), consistent with established roles of aromatic residues in carbohydrate recognition, including N-acetylated sugar moieties^50,51^. Consequently, the mean D297–acceptor distance decreased to 14.1 Å compared with WT (29.2 Å in WT-A and 18.6 Å in WT-B; **Table 3**), indicating increased disaccharide occupancy within the catalytic groove. Notably, both FtsW^E289G^IQLB superfission replicates also exhibited moderate-to-strong acceptor engagement (**Fig. S17C**).

Taken together, these results showed that FtsN stabilizes acceptor binding through an asymmetric acceptor clamp: Y379 hydrogen bonding and W138 methyl–aromatic packing engage the acceptor L2 disaccharide from opposite sides of the binding site. These interactions constrain the acceptor disaccharide so that its nonreducing-end GlcNAc is oriented toward D297, a prerequisite for catalysis (**Fig. 3B**).

### Long donor PG chains further stabilize the acceptor site

Single-molecule tracking from our group and others has shown that the activated FtsWIQLB complex is highly processive, capable of polymerizing ∼200-1000 disaccharides in a single synthesis run^22–24,52^. This processive movement requires ongoing transpeptidation by FtsI^22^, in which the nascent PG strand is cross-linked into the existing cell wall while new L2 molecules are repeatedly and rapidly captured at the acceptor site on timescales of hundreds of milliseconds. However, our simulations indicated that acceptor-site binding is highly dynamic and intrinsically unstable even when partially stabilized by FtsN. We therefore asked whether modeling a more physiological state, incorporating a long, membrane-anchored donor PG chain, might reveal additional mechanisms to stabilize the acceptor site for continuous, processive polymerization.

To test this possibility, we built FtsWIQLB-N^58-108^ systems in which we extended the donor L2 to Lipid IV (L4) or Lipid XXXII (L32) by appending additional PG disaccharides using standard glycosidic-bond geometry (**Fig. 3D**, Methods). We then energy-minimized and equilibrated the models to relieve steric clashes while preserving the donor-site pose (Methods). During equilibrium MD, the added PG disaccharides relaxed into a compact ensemble, whereas the peptide stems remained highly dynamic. In the L4 simulation, the short PG chain was largely unconstrained and explored the central cavity, making only transient contacts with ECL2 **(Supplementary Movie 9**). By contrast, the longer L32 chain, docked similarly at the start, formed a stable, extensive interface with FtsWI and the L2 Acceptor (**Supplementary Movie 10).**

Excitingly, the FtsWIQLBN^58-108^-L32^D^–L2^A^ simulation revealed a pronounced stabilization of the acceptor site relative to the corresponding L2^D^ systems. Direct contacts between the extended donor chain and FtsW appeared to drive inward closure of ECL1, which encased the acceptor disaccharide and effectively blocked its dissociation from the binding site (**Fig. 3D**). Accordingly, the L32^D^ system produced the smallest donor-acceptor distances (11.5 Å, **Fig. S17A,** purple bars; **Table 3**), and the largest acceptor-site BSA values under any condition examined (**Fig. 3C**, purple bars; **Fig. S17C).** Interestingly, the BSA distribution exhibited two populations: a ∼ 600 Å² state overlapping the FtsN-bound L2^D^ ensemble, and a higher-BSA population (∼750–800 Å²) that became enriched late in the trajectory (**Fig. S17G**). The L4^D^ simulation populated intermediate BSA values between the short L2^D^ and long L32^D^ systems (**Fig. S17F**), suggesting a length-dependent contribution of the donor PG chain to acceptor-site stabilization during processive synthesis.

Concurrently with ECL1 closure in the L32^D^ simulation, we also observed a marked reorientation of W138 in ECL2. In WT and FtsN-L2^D^ simulations, W138 predominantly adopted an outward-facing position, with only brief excursions into an alternative rotamer conformation. In the L32^D^ simulation, W138 strongly favored an inward-facing rotamer (χ angle ≈ −50°, **Fig. 3E-F**), enabling persistent methyl–aromatic packing against acceptor acetyl groups and allowing MurNAc to approach the base of the catalytic groove toward the end of the trajectory (**Fig. 3G**). As a result of the stabilization of this geometry, the mean D297-acceptor distance further decreased to 9.6 Å (**Table 3**). Together with sustained Y379 hydrogen bonding (**Fig. 3G**), this rotameric switch closes the acceptor clamp around the acetyl groups and enforces a well-defined geometry on both GlcNAc and MurNAc in the acceptor site, thereby helping to position and align the acceptor for catalysis.

### Replicates of long donor PG chain simulations exhibit similar acceptor site stabilization

To ensure that acceptor-site stabilization by extended donor chains did not arise from limited simulation time or a particular starting configuration, we first extended the FtsN^58-108^-L2^D^, L4^D,^ and L32^D^ simulations to 4, 4, and 2 µs, respectively (**Table 2**). With L4^D^, acceptor binding remained highly dynamic, and longer trajectories primarily broadened the range of transient poses rather than revealing new binding modes (**Supplementary Movie 9**); In contrast, the L32^D^ simulation generated a more complex interaction landscape, as the extended glycan chain and multiple peptide stems formed numerous nonspecific contacts and increased the likelihood of trapped metastable states. Therefore, we next constructed three independent 2 µs replicates of the L32^D^ system using randomized initial velocities (Methods).

Across all three L32 replicates, acceptor-site stabilization was reproducibly enhanced, although the persistence of specific acceptor clamp interactions varied. In replicate A, inward-facing W138 packing occurred for ∼60–80% of the trajectory and Y379– acetyl hydrogen bonding for ∼50–70%, whereas replicates B and C sampled these contacts more intermittently (W138 packing at ∼25–50%; Y379 hydrogen bonding at ∼10–50%) (**Fig. S18A,B,D**). Deep acceptor insertion marked by W138-MurNAc contacts was likewise enriched in replicate A (∼20–25%) but rare in replicates B and C (<5%) (**Fig. S18C**). Despite these quantitative differences, all L32^D^ simulations showed a clear shift toward inward W138 orientations relative to shorter-donor systems (**Fig. S18D**), indicating that long donor chains consistently bias the ensemble toward a clamp-stabilized acceptor geometry. Together, these analyses supported a model in which a processive sPG synthase complex exploits long donor-chain engagement to promote stable, specific acceptor-site binding and to position the acceptor glycans closer to D297, even as the nascent donor chain samples a broad conformational landscape.

### Long PG chain-FtsI anchor loop interactions reinforce acceptor binding

In all our simulations containing FtsN^58-108^, we observed that the donor peptide stems frequently engaged with the FtsI anchor loop (**Fig. S19A**). The FtsI anchor-loop is a short periplasmic loop that sits directly over FtsW ECL4 in the catalytic groove, which we previously proposed to activate FtsW by modulating its interactions with FtsW ECL4^20^. To test whether and how the donor PG length modulates these contacts, we quantified interactions between donor peptide-stem amino acids and a conserved arginine pair (R213 and R216) on the FtsI anchor-loop (**Fig. S19A**). We counted a trajectory snapshot as “tethered” when any donor stem amino acid contacted either arginine (N---O < 3.5 Å). Under this binary definition, donor-anchor-loop tethering occurred in ∼43% of the FtsWIQLBN– L2^D^–L2^A^ simulation, compared to ∼1% in the corresponding WT simulations lacking FtsN (**Fig. S19B**). Extending the donor to L4^D^ modestly increased tether occupancy (∼58%), consistent with additional stem acids becoming available to engage the anchor-loop (**Fig. S19A**). With an L32^D^ donor, tethering frequencies spanned a much broader range across replicates (29%, 51%, 92%), reflecting multiple modes by which the extended polymer can engage the anchor loop and form more widely distributed, transient interactions along the PG chain **(Fig. S19B).** Notably, in one L32^D^ trajectory, a glycan acetyl group from the fourth donor disaccharide formed a persistent contact with the anchor-loop aromatic residue Y214. This acetyl proximity (scored as when any acetyl methyl carbon approached any Y214 ring atom within 4.0 Å) persisted for >80% of the simulation after the first ∼250 ns **(Fig. S19D).** We interpret this acetyl–Y214 interaction as one example of the broader PG chain–anchor-loop engagement that a long donor polymer can access while remaining bound in an activated donor site. Although the precise interacting residue will likely vary between species (Y214 is only 11% conserved), any combination of salt-bridge and sugar contacts that constrain the donor chain near the FtsW/FtsI interface should favor the engagement between the donor chain and the FtsI anchor loop.

Importantly, FtsI anchor-loop engagement provides a direct physical mechanism for acceptor-site stabilization: by biasing the donor polymer to remain positioned over the catalytic cavity, tethering increases the likelihood that the extended chain also presses against FtsW ECL1, promoting inward ECL1 closure around the acceptor site (**Fig. 3D**) and thereby reinforcing the acceptor clamp and suppressing acceptor escape (**Fig. 3E–G**). In this way, a long donor chain reinforces the geometry-defining features that convert transient acceptor engagement into a more reliable, processive configuration. In this regime, donor elongation and acceptor-site stabilization become mutually reinforcing, progressing from a pre-initiation configuration (**Fig. 3I**, left) to a processive synthase after only a few rounds of polymerization (approximately L8, **Fig. 3I**, right).

### Chemically blocking the acceptor-site cavity using S382C causes loss of FtsW activity

Our simulations indicate that donor-site activation alone is not sufficient for productive catalysis unless the acceptor disaccharide is also stabilized in a defined packing pose against the periplasmic rim of the catalytic groove. In activated complexes, this acceptor interface is characterized by high glycan burial in FtsW and a preferred disaccharide orientation shaped by contacts with W138 and neighboring basic residues (**Fig. 3B–D**). We therefore asked whether chemically blocking this acceptor-side cavity would compromise FtsW function *in vivo*.

To test this idea, we mutated S382 to cysteine (**Fig. 3J, Fig. S3).** S382 occupies a highly conserved and solvent-accessible position near residues implicated in acceptor stabilization (W138, R145, R84). We reasoned that covalent modification at this position should inhibit FtsW by disrupting acceptor packing. Consistent with this expectation, an FtsW(S382C)-Halo fusion supported normal growth with few elongated cells in the absence of MTSES when WT FtsW was depleted (10.3% of cells elongated, range 2.4–15.9%, versus 3.4%, range 0–6.8%, for WT FtsW, n = 2 replicates; **Fig. 3K**, top right, **Fig. S7C**, strain AP766, **Supplementary Table 1),** whereas addition of MTSES produced a strong filamentation phenotype and abolished constriction (**Fig. 3K**, bottom right, 89.1% of cells elongated, range 73.1–98.3%, versus 10.3% without MTSES; cell length 24 ± 7 µm; n = 3 biological replicates, 161 cells; **Fig. S7C**, **Supplementary Table 9**). This MTSES-dependent loss of division paralleled chemically blocking the donor side of the groove (**Fig. 1H**) or constraining ECL2 dynamics (**Fig. 2J**). The whole-cell fluorescence intensity for the FtsW(S382C)-Halo fusion was ∼80% of FtsW-Halo in the merodiploid background (AP894, **Supplementary Table 1)** when treated with MTSES (**Fig. S8A, B**), and FtsW(S382C)-Halo retained robust septal localization, with midcell JF646 fluorescence comparable to that of the WT FtsW-Halo signal (**Fig. S8A**, **C**). Under these merodiploid conditions with 100 µM IPTG induction, FtsW(S382C)-Halo produced only a barely measurable dominant-negative effect, with or without MTSES (5.7 ± 0.2 and 5.8 ± 0.3 µm; 1.9% and 3.7% of cells elongated without and with MTSES, ranges 0–3.8% and 0–9.5%, versus 0% and 3.6% for the FtsW-Halo control; **Fig. S9A,C**, Supplementary **Table 10**). Mass spectrometry to identify direct modification of S382C was similarly unsuccessful (**Fig. S10, Supplementary Table 8**).

To further confirm the above observations, we performed SMT of FtsW(S382C)-Halo (AP766, **Supplementary Table 1; Fig. S11C,D).** In the absence of MTSES, FtsW(S382C)-Halo molecules displayed slow, processive trajectories characteristic of active sPG synthesis, with 25% of trajectory segments showing the active slow-moving population, 12% fast-moving, and 63% stationary (N = 109 segments). With MTSES, the slow-moving population fell to 3% while the fast-moving population increased to 29% (N = 181 segments, **Supplementary Table 5),** indicating that chemical blockade of the acceptor site shifted FtsW(S382C)-Halo from the active synthesis population to the inactive, Z-track-associated population. Together, these results supported a model in which the periplasmic acceptor-site cavity must remain intact and accessible for FtsW to function, and that occluding this interface is sufficient to disrupt cell division.

### MD simulations of chemically modified FtsW cysteine variants support inhibited FtsW function

To further verify whether the engineered cysteine modifications produce their inhibitory effects through the mechanisms predicted by the simulations, we explicitly modeled each variant with the corresponding cysteine-dependent bond in the FtsN-bound complex of FtsWIQLB-L2^D^-L2^A^ (**Fig. 4**; **Table 2**). Importantly, each variant was built from the same FtsN-bound starting structure used as the representative activated trajectory, so that these minimal modifications directly test whether disrupting donor-site gating (**Fig. 4A**), ECL2 switching (**Fig. 4B**), or acceptor-site packing (**Fig. 4C**) is sufficient to prevent the complex from reaching a tightly bound, active state.

**Figure 4.**
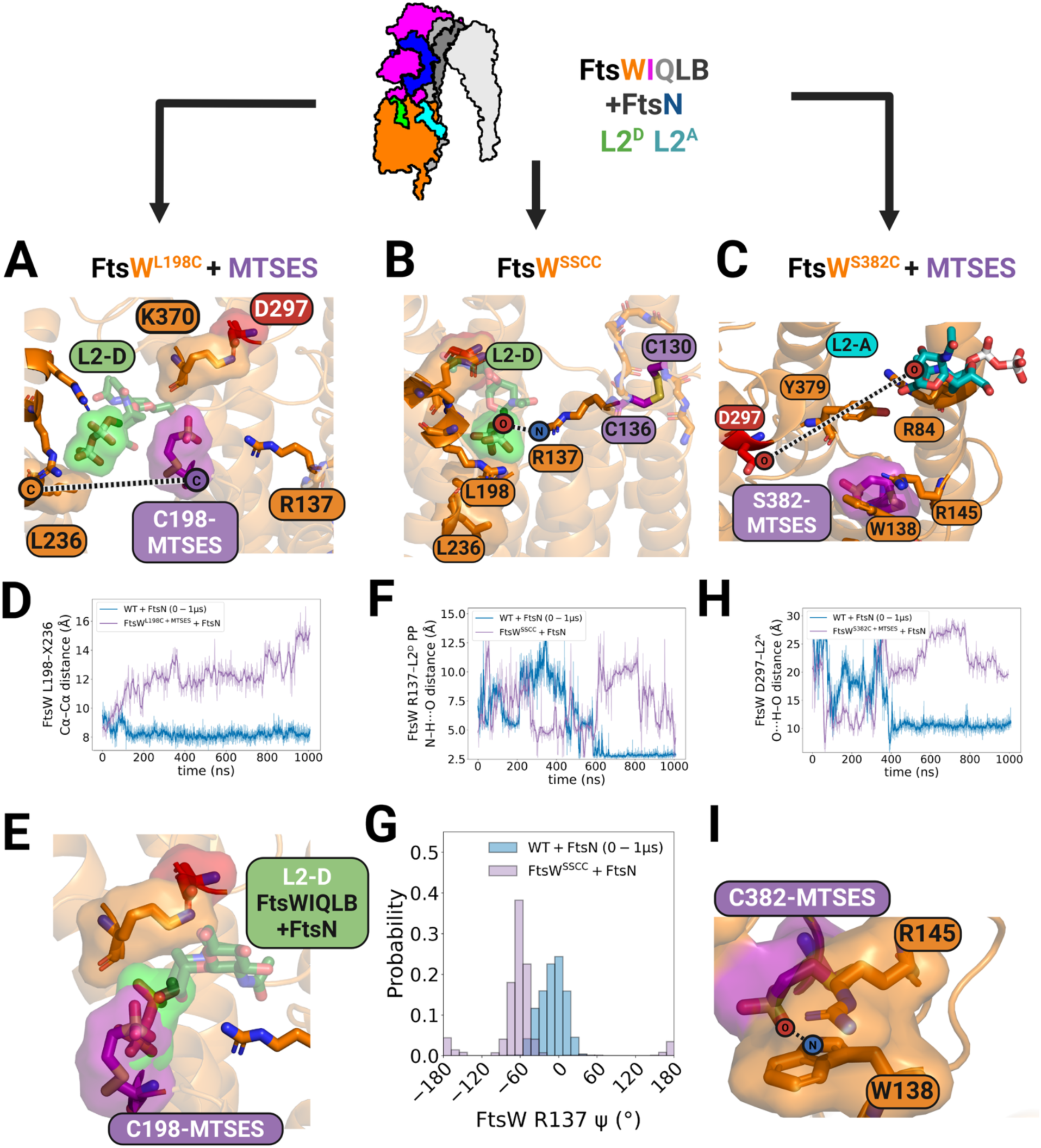
MD modeling of engineered FtsW cysteine mutants reveal compromised functions. **(A–C)** Representative structural views of each engineered perturbation modeled in the FtsN-bound FtsWIQLB–L2ᴰ–L2ᴬ complex: the donor site of FtsW(L198C)–MTSES (**A**), ECL2 of FtsW(SSCC) (**B**), and the acceptor site of FtsW(S382C)–MTSES (**C**). Dashed lines mark the distances used to define donor-site gating (L198–L236), R137–donor contact (N–H···O), and acceptor engagement (D297–acceptor), quantified in the panels below. **(D)** Outer-gate dynamics. Time traces of the FtsW L198–L236 Cα–Cα distance across the 1-µs trajectories for the unmodified FtsN-bound reference (blue) and FtsW(L198C)+MTSES (purple) showed significantly increased distances between L198 and L236. **(E)** Steric occlusion of the donor site: the C198–MTSES adduct overlaps the volume occupied by donor L2 in the bound pose; panel (A) is aligned to the last frame of the activated reference simulation, with reference L2 shown as transparent sticks. **(F)** R137–donor contact. Time traces of the R137–L2ᴰ pyrophosphate N–H···O distance for the FtsN-bound reference (blue) and FtsW(SSCC) (purple) show that the disulfide-constrained ECL2 fails to coordinate the donor substrate. **(G)** R137 ψ-angle probability distributions for the same trajectories as (**F**) show that it is outside of the active switch state (defined as −90° < ψ < 90°). **(H)** Acceptor engagement. Time traces of the D297–acceptor distance for the FtsN-bound reference (blue) and FtsW(S382C)+MTSES (purple) show the loss of stable acceptor-site packing. **(I)** Acceptor-clamp diversion. In the FtsW(S382C)+MTSES trajectory, the C382–MTSES adduct packs against W138, whose indole ring stacks with the acceptor-coordinating R145 guanidinium; corresponding time-resolved contact occupancies are shown in **Fig. S18E**.

As shown in **Fig. 4A**, MTSES-modified L198C positioned the MTSES group in the donor-site cavity, between the donor L2 and the catalytic groove, and shifted the TM6–TM7 outer gate toward more open conformations (**Fig. 4D**, purple curve), different from the tightly closed donor-site geometry observed in the FtsN-bound reference complex (**Fig. 4D**, blue curve). Notably, the MTSES adduct occupied the same region sampled by one of the donor L2 phosphate groups in the unmodified complex (**Fig. 4E**, light green), providing a direct steric and electrostatic mechanism for impaired donor engagement.

By contrast, introducing the SSCC disulfide bond into the FtsWIQLB-L2^D^-L2^A^-FtsN complex (**Fig. 4B**) led to large distance separations between R137 and the donor pyrophosphate, with only ∼ 3% of the trajectory dropping to the level observed in the FtsWIQLB-L2^D^-L2^A^-FtsN complex at the end of the simulation (**Fig. 4F**, compare purple and blue curves), despite FtsW retaining an active-like R137 ψ angle (−90° < ψ < 90°) throughout the trajectory (**Fig. 4G**). Thus, the engineered disulfide does not simply lock ECL2 into an inactive backbone conformation. Instead, it appears to restrict the conformational flexibility R137 needs to engage the donor substrate productively.

Finally, we examined the effect of MTSES modification of S382C in the same FtsWIQLB–L2^D^–L2^A^-FtsN complex and performed a 1-µs equilibrium MD simulation (**Fig. 4C**; **Table 2**). Consistent with the imaging results in **Fig. 3K**, MTSES modification did not cause a large-scale collapse of the acceptor pocket; instead, the negatively charged sulfate group produced pronounced local reorganization of residues implicated in acceptor recognition, leading to deteriorated acceptor engagement (**Fig. 4H**). The S382C-MTSES trajectory showed substantially larger separations (∼20 – 30 Å) between the catalytic residue D297 and acceptor over the latter half compared to that in the original complex (**Fig. 4H**, purple curve), more closely resembling the weak acceptor engagement observed in simulations lacking FtsN (**Fig. 3A**). This disengagement was accompanied by a local reorganization of W138, which formed persistent interactions with the MTSES sulfate, while the acceptor-coordinating R145 had its guanidinium group stacked against the W138 indole ring (**Fig. 4I**). This W138–R145 stacking interaction was strongly enriched in the S382C trajectory, occurring in approximately 74 % of snapshots compared with ∼5 % in the unmodified FtsN-bound reference simulation (**Fig. S18E**). Thus, the S382C-MTSES adduct diverted two residues normally engaged in positioning the incoming acceptor L2 molecule.

Taken together, the three perturbation simulations converged on the same FtsW-inhibited outcome by reducing access of key FtsW residues to the primed substrate configurations observed in the activated FtsWIQLB–L2^D^–L2^A^-FtsN complex, consistent with our proposed activation mechanism. Thus, these results established a direct connection between the structural model of FtsW substrate binding and the functional state of FtsW in living cells.

### Conserved and divergent L2 binding features in diderm and monoderm FtsW homologs

To test whether the L2-binding features identified in *E. coli* FtsW are conserved beyond the genus *Escherichia*, we generated AlphaFold ^53^ models of FtsWIQLB homologs from representative diderm (Gram-negative) bacteria (*Klebsiella pneumoniae*, *Caulobacter crescentus,* and *Pseudomonas aeruginosa*) and the monoderm (Gram-positive) bacterium *Bacillus subtilis* (Methods). The predicted complexes showed substantial structural similarity, consistent with a conserved structure-function relationship analogous to that observed in *E. coli*. We then docked species-specific L2 molecules into the donor and acceptor sites of each FtsW homolog (**Table 2**) and performed short, unbiased MD trajectories (92–100 ns) for each complex to assess the conservation of key binding features.

Across the three diderm homologs, the overall electrostatic organization of the donor-site binding pocket in FtsW was preserved (Fig. S20A–C). Specifically, the *Klebsiella pneumoniae* model showed nearly identical coordination of the donor L2 pyrophosphate group by positively charged arginine residues *kp*R231 and *kp*R228 in AH1 (corresponding to *E. coli* FtsW *ec*R246 and *ec*R243 respectively) and *kp*K355 in ECL5 (the *ec*K370 inner gate equivalent, **Fig. S20A).** In the *Caulobacter crescentus* model, there is no FtsI anchor loop interfacing with FtsW ECL4 (**Fig. S20B**, top), and FtsW lacks arginines at the positions corresponding to *ec*R145 and *ec*R243. However, *cc*R65, *cc*R225, and *cc*K342 formed sustained interactions with L2 that replicated the electrostatic logic of the donor site, acceptor site, and inner gate, respectively (**Fig. S20B**, bottom). This suggests that the core binding and regulatory motifs of *E. coli* FtsW are also conserved in α-proteobacteria despite global divergence in FtsWIQLB structure. Interestingly, in the *Pseudomonas aeruginosa* FtsWIQLB complex, *pa*K348 in ECL5 (*ec*K370 equivalent) had negligible electrostatic contact with donor L2, analogous to an opened inner gate; its *pa*R115 in ECL2 consistently oriented towards the donor L2, corresponding to the flipped *ec*R137 position in the activated, FtsN-bound *E. coli* complex (**Fig. S20C**) and differing from the more buried conformations of *kp*R122 and *cc*R118 (**Fig. S20A,B**). These features may help explain why the paFtsWIQLB complex has a less stringent requirement for activation by FtsN *in vivo* and *in vitro*^7,54^.

The *Bacillus subtilis* homolog complex showed similar donor-site coordination by conserved arginine residues *bs*R223 and *bs*R220 (*ec*R246 and *ec*R243 equivalent) and acceptor-site engagement by *bs*R50 (*ec*R84). However, it diverged at positions that participated in donor-site activation in diderm FtsW complexes (Fig. S20D): the *ec*K370 and *ec*R137 equivalents are *bs*T344 and *bs*S105, respectively, implying that the donor-site architecture and its electrostatic gating are organized differently in this genus. Furthermore, in a short *B. subtilis* trajectory, we observed that a Mg^2+^ ion rapidly localized near the donor pyrophosphate and remained coordinated by *bs*T344 and a conserved glutamic acid residue (*bs*E270) together with the donor pyrophosphate for a substantial fraction of the simulation (**Fig. S21A, C**), whereas comparable Mg^2+^ localization was not observed in any *E. coli* trajectories under identical ionic conditions (**Fig. S21B**).

To test whether these electrostatic differences reflected broader evolutionary patterns, we analyzed FtsW sequence alignment from the *Pseudomonadota* (primarily diderm) and *Bacillota* (primarily monoderm) phyla separately. In *Pseudomonadota,* the ECL2 *ec*R137-W138 motif was 100% conserved (**Fig. S22A**, **top**), whereas in *Bacillota* there was no dominant conserved residue at the X105-W106 position (equivalent of ecR137-W138), with R or S occurring at similar frequencies at position 105 (**Fig. S22A**, **bottom**). In ECL5 the acceptor-site SYGGSS motif was conserved in both phyla, but the inner gate position was universally K in *Pseudomonadota* and predominately T in *Bacillota* (Fig. S22B). Together, these observations likely suggest that monoderm homologs may rely more heavily on metal coordination to stabilize a catalytic donor-site geometry, whereas diderm homologs achieve comparable stabilization primarily through protein side-chain interactions.

### A protein language model task head identifies binding-site residues across 6,859 FtsW orthologs

Finally, we asked whether the FtsW L2 binding sites identified by our structural and functional analyses exhibit a ligand-binding signature detectable from sequence alone. We applied AiPP-LigBind^55^, a task head built on the protein language model (pLM) ESMC^56^, to rank residues by their intrinsic ligand-binding propensity across sequences from 6,859 FtsW orthologs. From the sequence alignment used in **Fig. S3,** we generated a 414-residue consensus FtsW sequence, retaining *E. coli*-referenced residue numbering, and mapped the conservation (**Fig. S23A**) and AiPP-LigBind score (**Fig. S23B**) onto an ESMFold2^56^ model of the consensus FtsW sequence. The two scores overlapped most strongly around the donor and catalytic regions, including R137, R243, R246, and D297, whereas the acceptor side scored lower and less uniformly (**Fig. S23B**; Supplementary **Table 11**).

We then applied AiPP-LigBind to each of the 6,859 FtsW sequences and mapped the predictions back onto the *E. coli*-referenced scaffold. At the exact aligned positions, R137, R243, R246, and D297 ranked among the 20 highest-scoring residues in 99.2%, 83.1%, 99.7%, and 93.2% of orthologs, respectively (**Fig. S23C** and **Supplementary Table 12).** Other high-ranking residues within ±2 sequence positions of L198, K370, Y379, and S382 were also ranked within the Top20 and in 99.9%, 94.4%, 99.9%, and 99.1% of orthologs, respectively (**Fig. S23C**). In contrast, R84, R145, and L236 showed little ligand-binding propensity at either the exact position or the immediate sequence neighborhood (**Fig. S23C**).

Thus, the pLM task head identified the donor and catalytic residues in nearly all FtsW orthologs but localized the acceptor and gating sites only approximately, flagging positions adjacent to the true residues. This asymmetry is consistent with our MD simulations: a gated, electrostatically defined donor pocket versus an acceptor site that becomes competent only upon FtsN-induced clamping and PG chain engagement.

## DISCUSSION

In this work, we defined the putative donor and acceptor L2 binding sites in FtsW (**Fig. 1**) and identified inhibitory and activating conformations of the *E. coli* FtsWIQLB complex using all-atom MD simulations combined with targeted mutagenesis (**Fig. 2-3**). We propose that an L2 molecule can access the inner donor binding pocket formed by FtsW’s AH1, TM6, and TM7 through an outer gate between TM6 and TM7 (Fig. S6). The acceptor binding site is located at FtsW’s TM2 and opens to the outside. FtsW’s ECL2 and ECL5 encompass both donor and acceptor binding sites and the catalytic groove. Positively charged residues, R246 and R243 at the donor site, and R84 and R145 at the acceptor site, coordinate L2’s pyrophosphate group and stabilize the binding of L2 at each site (**Fig. 1F**). In the absence of any activating factor, the WT FtsWIQLB complex exists in a self-inhibited state, with a closed inner gate formed by the packing between K370 and TM6 that prevents the donor L2 from approaching the acceptor L2, an open outer gate that allows the exit of the donor L2, and unstable acceptor L2 binding (**Fig. 5A**).

**Figure 5.**
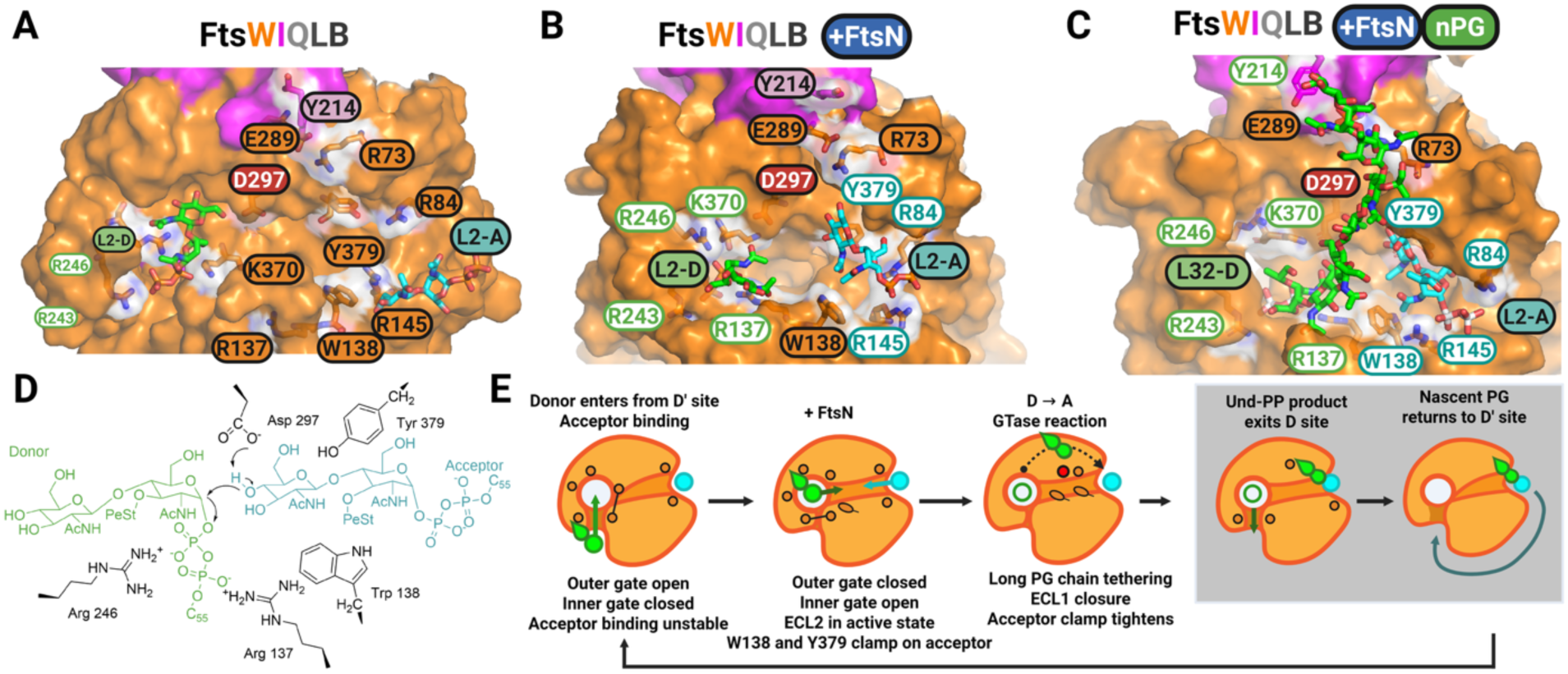
Model for FtsN-dependent activation and long donor PG chain stabilization in FtsWIQLB. **(A–C)** Representative periplasmic views of FtsW’s donor and acceptor sites showing progressive opening of the central catalytic groove from a self-inhibited FtsWIQLB-L2^D^-L2^A^ complex (**A**), to a primed FtsWIQLB–FtsN-L2^D^-L2^A^ complex (**B**), and finally to a processive FtsWIQLB–FtsN-L32^D^-L2^A^ complex (**C**). Key residues lining the donor and acceptor cavities are labeled, with those actively engaged with the substrate molecules labeled with white backgrounds and font color corresponding to the donor (green) and acceptor (cyan) molecules. **(D)** Proposed processive GTase cycle for donor/acceptor binding, catalysis and exchange. A new donor substrate enters through the outer gate to occupy the donor site, while acceptor binding is initially unstable in the self-inhibited state (outer gate open; inner gate closed). FtsN binding promotes closure of the outer gate and opening of the inner gate, stabilizing acceptor engagement through W138 and Y379 clamping on the acceptor, and enabling a pre-catalytic donor–acceptor geometry through ECL2 dynamics. Long donor chain tethers to FtsI’s anchor loop and further stabilizes acceptor binding by closing ECL1 and tightening the acceptor clamp. Glycosyltransfer (GTase) chemistry transfers the glycan from donor to acceptor to form nascent peptidoglycan (NPG). After catalysis, the Und-PP product exits the donor site, and the newly elongated glycan repositions to reoccupy the donor site for the next polymerization cycle. The last two steps (gray box) are subject to further investigation. **(E)** Schematic of the glycosyl transfer step showing formation of the β-1,4 glycosidic bond by nucleophilic attack of the acceptor O4 on the donor anomeric carbon (C1), with Asp297 positioned as the catalytic base and Arg137 contacting the donor pyrophosphate.

In the presence of the essential domain of the activator protein FtsN or the superfission variant FtsW(E289G), we observed dramatic conformational changes in FtsW: K370 formed a salt bridge with D297, opening the inner gate; the outer gate closed, preventing the escape of the donor L2 molecule; the acceptor binding was also stabilized due to the acceptor clamp formed by W138 and Y379. Most importantly, ECL2 re-oriented, and R137 flips to coordinate donor pyrophosphate and bring it deeper into the catalytic groove to the acceptor L2 (**Fig. 5B**).

When the donor site was occupied by a long nascent PG chain (L32), we observed a pronounced shift toward stable, high-contact acceptor binding, driven by inward closure of ECL1 that encases the acceptor headgroup and suppresses dissociation, together with an inward-facing W138 rotamer that sustains acceptor clamp contacts (**Fig. 5C**). In this context, tethering of the extended PG chain to the FtsI anchor loop further biases the donor polymer toward the FtsW/FtsI interface, promoting ECL1 closure and thereby reinforcing the acceptor clamp.

Taken together, these results supported a model in which FtsN primes a catalytically competent donor site and enhances acceptor binding, while subsequent polymer growth further stabilizes acceptor engagement to sustain processivity (**Fig. 5D**). This model proceeds as a stepwise transition from an initial electrostatically guided encounter, dominated by pyrophosphate coordination and favorable lipid–protein packing, to an activated state in which key gating elements and loop conformations remodel to retain the donor substrate, clamp the acceptor in a productive pose, and bring the two headgroups into a configuration for chemistry as depicted in the *S_N_2* reaction (**Fig. 5E**). Below we discuss in detail key considerations and limitations of the model.

### FtsWIQLB as the minimal, catalytically competent complex

We chose to use the FtsWIQLB complex, rather than FtsW or FtsWI alone, as the minimal, catalytically competent assembly for evaluating substrate binding and activation because of several considerations. First, FtsW alone displays minimal GTase activity *in vitro* and becomes substantially more active only upon FtsI binding^4^. Addition of the FtsQLB subcomplex further enhances FtsW’s GTase activity in the reconstituted *Pseudomonas aeruginosa* system^54^. Whether *E. coli* FtsQLB enhances or restrains FtsWI in the absence of FtsN has not been determined. Second, previous immunoprecipitation^57^ and single-molecule imaging studies^20^ have established that FtsWIQLB forms a stable complex that supports processive septal PG synthesis both *in vitro* and *in vivo*. Third, genetic evidence suggests that FtsN’s essential domain binds FtsI and FtsL at the AWI region in the periplasm, triggering conformational changes at the interface between FtsI’s anchor loop and FtsW’s ECL4, which are critical for FtsW’s activation^19,20^. Therefore, only in the context of the full FtsWIQLB complex can we investigate the structural remodeling and activation of FtsW by activating factors in a physiologically meaningful context.

### L2-bound FtsWIQLB model provides a structural explanation for prior dominant-negative mutants

Our search for donor and acceptor binding sites in FtsW was guided by sequence conservation (**Fig. S3**), structural homology^8,31^, and an extensive body of genetic studies that identified critical residues in the FtsWIQLB complex^13,14,22,33,34,58^. In turn, the resulting L2-bound model provides a coherent structural explanation for previously observed defects of several dominant negative mutations of FtsW that map to either the donor site, the acceptor site, or the ECL2 switch region (Li et al^33^, **Table 4**). These dominant-negative variants are not simply misfolded or nonfunctional; they assemble into stable complexes with FtsI and FtsQLB and compete with WT FtsW, effectively “poisoning” an otherwise active complex upon overexpression. Many of these residues are predicted to be critical in the FtsWIQLB-L2^D^-L2^A^ model. For example, R243 and R246 are both critical electrostatic contacts for the donor ligand in our computational FtsW model, and mutation of either residue to histidine leads to a dominant negative phenotype. These observations are consistent with the donor site requiring strong guanidinium-mediated coordination of the donor pyrophosphate, stabilizing it as both a bound substrate and a leaving group in the GTase reaction^59^. Our simulations further implicate R137 in pyrophosphate coordination when ECL2 is in the active state, which is consistent with the dominant-negative R137H. Finally, we propose that the W138A and Y379A mutations would directly destabilize glycan packing at the acceptor site by removing the aromatic side chains that position the acceptor by packing against the methyl groups in the acetyl moieties in GlcNAc and MurNAc.

**Table 4.** Mechanistic Mapping of known FtsW Dominant Negatives.

| **Mutation** | **Dominant-negative effect**<sup>33</sup> | **FtsW Location** | **Group** | **Why dominant negative?** |
|---|---|---|---|---|
| **K133I** | ++ | ECL2 | ECL2 | Unclear |
| **A135T** | ++++ | ECL2 | ECL2 | (disrupts ECL2 - ECL3 packing and by extension activation switch) |
| **A135V** | ++++ | ECL2 | ECL2 | (disrupts ECL2 - ECL3 packing and by extension activation switch) |
| **R137H** | ++ | ECL2 | ECL2 | Disrupts activation switch |
| **Q147E** | ++++ | ECL2 | Asite | Unclear |
| **Q147K** | ++ | ECL2 | Asite | Unclear |
| **P196T** | ++ | ECL3 | Dsite | (Disrupts outer and inner gate regulation ) |
| **L198I** | + | TM6 | Dsite | Disrupts Dsite binding / outer and inner gate regulation |
| **G199A** | +++ | ECL3 | Dsite | (Disrupts outer and inner gate regulation ) |
| **Y242H** | ++++ | ECL4 | Dsite | (proximity to the donor-site glycan) |
| **R243H** | +++ | ECL4 | Dsite | Disrupts electrostatic coordination |
| **R243L** | ++++ | ECL4 | Dsite | Disrupts electrostatic coordination |
| **R243P** | ++ | ECL4 | Dsite | Disrupts electrostatic coordination |
| **R243S** | +++ | ECL4 | Dsite | Disrupts electrostatic coordination |
| **R246H** | ++ | ECL4 | Dsite | Disrupts electrostatic coordination |
| **G371S** | +++ | ECL5 | Dsite | (ECL5 flexibility; proximity to K370) |
| **G381V** | ++ | ECL5 | Asite | (ECL5 flexibility; proximity to Y379) |
| **G381D** | ++ | ECL5 | Asite | (ECL5 flexibility; proximity to Y379) |
| **W138A** | ++++ | ECL2 | Asite | Disrupts glycan binding and positioning |
| **D297A** | ++ | ECL4 | Catalytic | Disrupts catalysis |
| **Y379A** | ++ | ECL5 | Asite | Disrupts glycan binding and positioning |
| **G380A** | ++ | ECL5 | Asite | (ECL5 flexibility; proximity to Y379) |

We also assessed the dominant-negative behavior of the engineered cysteine variants directly. FtsW(SSCC) and FtsW(L198C) each produced a mild dominant-negative effect (12.9% of cells elongated for SSCC; 35.7% without MTSES and 60.1% with MTSES for L198C; versus 0% for the control), whereas FtsW(S382C) produced only a barely measurable one, with or without MTSES (1.9% and 3.7%, versus 0% and 3.6% for the control; mean of three replicate percentages in each case). This was unexpected, because substitutions at neighboring ECL5 positions — G371S, Y379A, G380A, and G381V/D — are dominant negative in the study. However, the assays differed in the expression level: Li et al. scored dominant negativity from pDSW208, a plasmid whose basal expression is sufficient to complement an FtsW depletion, and titrated IPTG induction to ≥125 µM for these mutants. By contrast, FtsW in our merodiploid strains expressed from pIQ required ∼ 50 µM IPTG, and we did not exceed 100 µM IPTG in the dominant-negative assay. Furthermore, S382C showed the lowest ensemble fluorescence of the three variants, indicating slightly lower expression. In contrast, FtsW(K370C) was strongly dominant-negative, underscoring the donor site’s sensitivity to steric and electrostatic disruption. It therefore remains possible that stronger induction would reveal a dominant-negative phenotype for all three variants.

Several other dominant-negative mutations, including Y242H, Q147E, and a cluster of glycine substitutions surrounding Y379, do not yet admit a clear mechanistic explanation within our current framework. These unresolved cases highlight key targets for future work aimed at understanding how pocket geometry, loop flexibility, transient interactions, and longer timescale conformational dynamics can collectively shape FtsW’s activation and processivity.

### Donor and acceptor binding sites align with experimental structures and reveal unresolved dynamic conformations

Currently, no experimentally determined structure of FtsWIQLB in complex with L2 is available. We constructed the FtsWIQLB-L2^D^-L2^A^ complex based on our previous MD model of the apo *E. coli* ecFtsWIQLB assembly. The resulting model aligns well with the apo cryoEM structures of the *Pseudomonas pa*FtsWIQLB homolog complex^7^, and the *E.coli* RodA-PBP2 paralog complex^8^, while additionally capturing conformations of protein segments and lipids that were not resolved in those structures. In particular, FtsW’s TM7 and AH1 region, key elements of the donor cavity identified in our simulations, were unresolved in the *pa*FtsWIQLB map. Our simulations indicate that this region is intrinsically dynamic **(Fig. S6),** likely undergoing repeated opening and closing during each catalytic cycle to regulate donor entry, site closure, and product release. Thus, the AH1-TM7 may function as an active gating element in FtsW rather than a static structural scaffold.

The analogous TM7-AH1 region was resolved in the cryoEM structure of the elongasome RodA-PBP2 complex, but with lower resolutions^8^. Notably, TM7 is markedly tilted toward the exterior of RodA and into the membrane, forming a cavity with TM6 and TM9 that closely matched the donor site identified in our MD simulations. An elongated lipid density in this cavity was assigned to the cleaved product UndPP. A second cavity near TM2 in RodA, proposed as a potential lipid-binding site by coarse-grained MD simulations, resemble the acceptor site we described here for FtsW. Critically, key electrostatic residues in these two cavities map directly onto the arginine residues flanking the FtsW catalytic groove (RodA R210, R48, and R109 map onto FtsW R246, R84, and R145, respectively; **Fig. S2B**) and coordinating the L2 pyrophosphate groups. These comparisons demonstrate that the donor–acceptor architecture and core electrostatic network identified in our FtsWIQLB– L2^D^–L2^A^ model are evolutionarily conserved across SEDS glycosyltransferases, supporting a shared mechanistic framework for lipid II polymerization.

### Donor site activation by conformational gating and R137-switching

In FtsWIQLB-L2^D^-L2^A^ simulations, entry of the donor L2 glycan into the FtsW catalytic groove is blocked by the packing of K370 against TM6, forming an inner gate that prevents the reactive atoms of the donor and acceptor L2 molecules from sampling the ∼4 Å separation expected immediately prior to an *S_N_*2 glycosyltransfer reaction. Furthermore, the TM6-TM7 outer gate remained open, which may favor the dissociation of the bound L2 substrate. This configuration (**Fig. 5A**) suggests that the WT complex adopts a self-inhibited state, in which these inhibitory interactions must be relieved by an activating factor before the substrates can attain a productive orientation for subsequent catalysis. A self-inhibited conformation of FtsWIQLB is consistent with a highly regulated sPG synthesis process essential for proper spatiotemporal septum morphogenesis and predicts that an activating factor must release the wild-type complex for enzymatic activity.

In *E. coli*, FtsN and specific superfission variants are known activators of FtsW^13,19-22,39^. To elucidate conformational changes associated with this activation, we modeled two representative activated complexes: FtsWIQLB bound to FtsN^58-108^ and FtsWIQLB harboring the superfission variant^13,20^ FtsW^E289G^. We focused on FtsN’s essential domain (FtsN^58-108^) because it is both necessary and sufficient for viability^21^, whereas the rest of FtsN, including its cytoplasmic tails, the long disordered periplasmic linker and the SPOR domain are dispensable for FtsW’s activation^45,46^. We previously proposed that FtsN^58-108^ binds to the FtsI and FtsL in the AWI region to modulate the interface between FtsI’s anchor loop and FtsW’s ECL4. The superfission allele FtsW^E289G^ maps to the same ECL4 and bypasses the requirement of FtsN (albeit yielding slightly longer cells), indicating that it constitutively mimics the FtsN-activated state of FtsW^22^.

In the FtsWIQLB-FtsN^58-108^-L2^D^-L2^A^ simulation, the K370-TM6 inner gate shifted toward an open configuration, the TM6-TM7 outer gate closed to prevent L2 escape, and ECL2 more consistently sampled a R137 Ψ conformation in which R137 became solvent accessible and positioned to placing positive charges at the pyrophosphate leaving group (**Fig. 5B**). Previous mutagenesis showed that R137H is dominant negative^33^ and we found that restricting ECL2 dynamics by an SSCC disulfide bond blocked division in most cells (77.2% elongated, **Fig. 2J**). Consistent with this loss of function, FtsW(SSCC) in the absence of DTT failed to support division in most FtsW-depleted cells, had a mild dominant negative effect when overexpressed in merodiploid cells, and retained robust midcell localization. The simulations added nuance to this picture: imposing the disulfide on an otherwise catalytically competent complex (with FtsN bound and ECL2 in the active Ψ state) did not force ECL2 into an inactive Ψ conformation, but the conformational restriction markedly diminished the R137–donor electrostatic contact that we propose is required for sPG synthesis. Notably, in the RodA-PBP2 cryoEM structure, the homologous ECL2 loop containing R101 (*ec*R137 equivalent) was not resolved, and the R101A mutation abolished RodA’s GTase activity^8^.

FtsW^E289G^ appeared to destabilize inhibitory interactions by disrupting the FtsW ECL4-FtsI anchor domain interface, which led to an open K370–TM6 inner gate and increased flexibility of ECL2 (compare **Supplementary Movies 5** and **6**). However, although the donor L2 headgroup rapidly enters the FtsW^E289G^ catalytic groove, we observed only transient, not stable, flipping of R137 in ECL2 to coordinate donor pyrophosphate, as seen in the FtsN^58-108^ complex (**Supplementary Movie 8**). These observations suggest that FtsW^E289G^ broadens the dynamic ensemble to include active configurations but fails to consistently sustain them like FtsN. In other words, release of inhibitory interactions is necessary but insufficient: R137 flipping in ECL2 is still required to act as an essential switch to initiate productive synthesis. This property hence could account for the elongated phenotype of *ΔftsN* ftsW^E289^ cells: growth is possible because the inner gate can open without upstream FtsN signaling, but cells are elongated due to poor regulation of substrate binding and inconsistent initiation of sPG synthesis by the ECL2 switch.

### Conservation of donor site activation and its variation in *Pseudomonadota*

Although some mechanistic details described above may be specific to *E. coli*, core features of our gate-switch model appear broadly conserved within the *Pseudomonadota* phylum (primarily diderm bacteria), as key residues such as ecK370 and ecR137 are nearly invariant (>99.7% conserved, **Fig. S22).** This conservation suggests that the K-ECL5 and R-ECL2 donor site architecture is effectively universal among species that require FtsN to activate FtsW. Indeed, in ∼100 ns simulations of *Klebsiella pneumoniae* and *Caulobacter crescentus* FtsWIQLB, the corresponding residues adopted inactive-like configurations, consistent with a conserved, self-inhibited conformation that must be released by FtsN-dependent activation. In contrast, in our simulations of *Pseudomonas aeruginosa* FtsWIQLB and the corresponding cryo-EM structure^7^, ECL2 is mostly a beta-sheet rather than a flexible loop, and the *ec*R137 equivalent residue *pa*R115 was locked in an active ψ state by the surrounding secondary structure (**Fig. S20C**). This “pre-flipped” ECL2 switch may contribute to the observations that FtsN is only conditionally essential in this species, and not required for FtsW activation *in vitro*^54,60^. Longer simulations will be needed to fully test this hypothesis, as our analysis of *E. coli* ECL2 dynamics indicates that the R137 ψ angles require ∼500 ns to stabilize.

### Activation of the Acceptor Site by long PG chains

MD simulations with two bound L2 molecules provide a practical model for the first GTase reaction, in which identical donor and acceptor substrates are simultaneously present in the FtsW cavity. However, sustained septal synthesis requires repeated recruitment and turnover of a new L2 acceptor each cycle^61^, making acceptor-site occupancy a central determinant of processivity. In our L2/L2 trajectories, acceptor binding is intermittent, with weak glycan packing and pyrophosphate coordination as the dominant persistent contact. Thus, donor-site activation alone may be insufficient to maintain a persistently engaged acceptor during the earliest synthesis cycles, consistent with single-molecule tracking in which the slow, synthesis-coupled FtsW population speeds up when precursor level is increased^22^.

To better approximate a processive regime, we simulated an FtsN-activated complex bearing a long donor polymer (L32). In the presence of L32, sugars 3–4 of the donor chain (the second disaccharide counting from the donor site) frequently contact the acceptor disaccharide and promote tighter acceptor-site binding, including enhanced packing of W138 against acceptor acetyl group **(Fig. 5C)** and generally within the acceptor clamp (**Fig. 3, Fig. S17).** Simultaneously, the extended polymer engages the FtsI anchor loop near Y214, creating a tether that links the catalytic groove to FtsI through the nascent PG. The acceptor-stabilizing effect is driven largely by interactions made by the first ∼8 sugars. Crucially, L4 also contains sugars 3–4 but lacks anchor-loop tethering via glycans and does not reliably engage with the acceptor site, suggesting a length threshold at which polymer growth becomes self-reinforcing **(Fig. S19D, Fig. S17F)**.

Together, these observations support a model in which the first ∼1–3 GTase cycles may be intrinsically inefficient because the nascent polymer is not yet long enough to engage the FtsI anchor domain^20^. Once the donor chain reaches a minimal length to sustain glycan–glycan contacts and anchor-loop tethering, acceptor retention becomes more reliable, and each completed round of synthesis reinforces the next by maintaining polymer engagement with FtsW and FtsI (**Fig. 5C**). In cells, this framework implies that additional divisome factors may be needed to bias early cycles toward productive acceptor capture and help the synthase enter a fully processive state. Such a length-dependent bias would also favor elongation over initiation, concentrating the L2 pool into fewer, longer chains rather than many short ones.

### Cysteine modifications to probe residue-specific mechanisms *in vivo*

Cysteine substitution combined with chemical modification has been widely used to inactivate bacterial periplasmic proteins at specific residues when no inhibitors are available^62–65^. This strategy allowed us to test, residue by residue, the mechanistic predictions of our MD simulations. In particular, we exploited disulfide bond formation between S130C and S136C of FtsW(SSCC) under non-reducing conditions to assess the importance of ECL conformational flexibility. While this approach does not substitute for NMR or hydrogen-deuterium exchange, or other structural methods, it complements these approaches by offering a route to residue-level mechanistic information in live cells. We also attempted to identify modified cysteines by HaloTag-enriched mass spectrometry, but the results were inconclusive because of the well-known difficulty of recovering peptides from polytopic membrane proteins and because the cysteines were positioned on either short or very long tryptic fragments (**Fig. S10**). Nevertheless, our wild-type controls in the absence of chemical modification, together with cell morphology, localization, and SMT analyses, showed that the observed phenotypes arise from modification at the intended residues rather than protein loss, validating this strategy for probing residue-specific mechanisms in vivo.

### Remaining mechanistic gaps and future directions

One limitation of our MD simulations is that the donor-acceptor distance never reached a true pre-catalytic geometry of < 4 Å, even in L32^D^ simulations, as would be expected immediately before an *S_N_2*-like glycosyltransfer reaction. Defining such near-reaction states will be particularly important for developing transition-state-like inhibitors. Nonetheless, the FtsN-activated L32^D^ complex likely represents our best current approximation to this configuration. This limitation leaves open the possibility that additional conformational adjustments by FtsW or the substrates, such as Y379 reorientation that steers the acceptor toward D297, occur on timescales not captured here. Using the ∼10 nm/s velocity of FtsW on the sPG track and the ∼1 nm contour-length per disaccharide^66^, we estimate that a single reaction cycle likely occurs on the order of ∼100 ms under favorable conditions. This time scale is currently beyond the reach of all-atom MD simulations for our system. Addressing this limitation will require enhanced sampling simulations together with experimental validations using high-resolution crosslinking or structural studies.

A second unresolved aspect is that our simulations did not capture product exit or the rebinding of the newly elongated PG chain back at the donor site (**Fig. 5D**, gray box), the two steps that pose the greatest challenge to sustained processivity. A model in which the acceptor L2 relocates through the catalytic groove to occupy the donor site after each catalytic cycle is an appealing hypothesis for achieving resetting and processivity, and an analogous model was proposed for RodA-PBP2 by Nygaard et al^8^. The donor and acceptor sites are separated by the protein core, and internal translocation of the Und-PP lipid tail from the acceptor to the donor cavity would require the tail to traverse the transmembrane region of FtsW. This mechanism would require substantial unfolding or rearrangement of the protein core or the substrate hydrophobic tail to leave the membrane’s hydrophobic environment, which we did not observe in any of our trajectories, including simulations with L2 bound only at the acceptor site.

Without such a translocation pathway, the growing polymer would need to disengage from the acceptor site and reengage at the donor site. Without a tethering interaction, repositioning of the new reducing end of the donor would be diffusion-limited, which is difficult to reconcile with the steady, processive synthesis observed *in vivo*. Our long-chain simulations suggest that upstream contacts with FtsI can keep nascent PG engaged while allowing the reducing end to sample the membrane, thereby biasing its return to the donor site. Because the extended chain is held largely at the FtsI anchor loop/FtsW ECL4 interface, it is conceivable that, after covalent attachment to the acceptor, the Und-PP-linked reducing end could reposition toward the donor site while the polymer stays anchored, allowing the acceptor to serve as the next donor without the chain fully dissociating. However, we have not yet tested whether this coupling alone is sufficient on its own or how it emerges during the earliest cycles before engaging with FtsI’s TPase domain. Defining how product exit and polymer re-routing are coordinated to preserve processivity thus represents the next mechanistic question and a direct bridge from the activation principles defined in this work to a complete cycle of PG synthesis.

A third limitation concerns the ECL2 activation switch. Although the engineered disulfide experiments support the proposed rearrangement, ECL2 dynamics have not been observed directly; experimental approaches such as site-directed crosslinking or spectroscopic probes would provide an important test of the model.

A final limitation concerns the membrane model. All simulations reported here used a pure POPE bilayer, chosen so substrate-bound trajectories could be compared directly with our previously published apo FtsWIQLB simulations, which used the same composition^20^; holding the membrane composition fixed isolates the effects of substrate binding from those of lipid composition. The *E. coli* inner membrane is a mixture of phosphatidylethanolamine, phosphatidylglycerol and cardiolipin, and the charged headgroups of the latter two could influence the dynamics of a protein whose substrate is itself an anionic lipid, particularly around the membrane-facing donor and acceptor cavities. Examining the effects of mixed PE/PG/cardiolipin bilayers on gating and substrate binding is an important direction for future work.

## METHODS

### System Preparation and Homology Modeling

The structural models of the *E. coli* FtsWIQLB and FtsWIQLBN complexes were prepared using coordinates from our previous 1µs Anton2 simulations^20^. The structural models for the divisome complexes for other organisms or FtsWIQLB(N 30-108) were generated using sequences acquired from UniprotKB and AlphaFold3 (DeepMind, 2024) multimer prediction via the AlphaFold 3 server^53^. AlphaFold3 was run with default settings, and the top-ranked models were selected based on predicted Local Distance Difference Test (pLDDT) scores and Predicted Aligned Error (PAE) metrics. Cytoplasmic regions of the proteins were truncated prior to preparation for MD simulations (residues 1–19 of FtsQ, 1– 45 of FtsW and 1–18 of FtsI were removed, whereas FtsL and FtsB were kept at full length). Where appropriate, predicted protein–protein interfaces were cross-validated against available cryoEM structures or homology-based models. Systems were constructed in a (180 Å)^3^ or (200Å)^3^ orthorhombic simulation cell. For the L32 simulations, the 200 Å cubic box was built with CHARMM-GUI Membrane Builder and contained a POPE bilayer (642 lipids in the upper leaflet and 645 in the lower leaflet), 191,134 TIP3P water molecules, 0.15 M NaCl and 0.01 M MgCl_2_ (554 Na^+^, 35 Mg^2+^ and 604 Cl^-^ ions).

### Protein Sequence Analysis

Protein sequences were retrieved from UniProtKB using keyword queries and processed with custom Python scripts. Entries were filtered to retain only sequences annotated with the target gene name (GN annotation) and to include a single representative sequence per organism (deduplicated by organism/species), removing redundant isoforms/duplicate records using the Organism (OS) annotation. Multiple sequence alignments (MSA) were generated with ClustalW^32^ via the Galaxy Pasteur ClustalW server. Alignments were analyzed with custom Python pipelines, and sequence-logo visualizations were generated using the Logomaker python library^67^. Residue conservation in FtsW was quantified across the 6,859-sequence alignment as the per-position amino-acid frequency among sequences containing an unambiguous residue at that position. Alignment gaps were retained in the sequence-logo analysis in **Fig. S3,** such that total amino-acid counts at each position also reflect column occupancy. For numerical residue-conservation calculations, gaps and ambiguous residues were excluded from the denominator, and amino-acid frequencies were calculated among sequences containing an unambiguous residue at the corresponding alignment position

To generate a consensus FtsW sequence, the same 6,859-sequence multiple-sequence alignment used for **Fig. S3** was analyzed using E. coli FtsW as a positional scaffold. For each alignment column corresponding to an E. coli residue, the most frequent standard amino acid was selected after excluding gaps and ambiguous residues, yielding a 414-residue consensus sequence. Conservation of each consensus position was defined as the fraction of non-gap, non-X sequences containing the modal amino acid at that position. A structural model of the consensus sequence was generated using ESMFold2^56^, and residue conservation values were mapped onto the corresponding structural positions. **Protein–L2 Complex Docking**

L2 molecules were docked manually into proposed donor and acceptor grooves using PyMOL (The PyMOL Molecular Graphics System, Schrödinger, LLC). Docking was guided by four criteria: (1) proximity of candidate arginine residues to D297; (2) presence of solvent-accessible basic residues for pyrophosphate coordination; (3) cavity geometry supporting substrate orientation; and (4) presence of nearby hydrophobic grooves to accommodate the C55 lipid tail. Initial placements were checked for steric clashes between protein and substrate and then equilibrated by short, restrained MD simulations prior to production runs using CHARMM-GUI protocols^68,69^. For dual-substrate simulations, both donor and acceptor L2 molecules were simultaneously docked into their respective binding sites.

### L2 and Nascent Peptidoglycan Parameterization

*E. coli* lipid II (L2) parameters were derived using CGENFF^69,70^ via the CHARMM-GUI Ligand Reader^69^. For simulations containing extended donor substrates (L4 and L32), nascent peptidoglycan (NPG) chains were constructed in CHARMM using custom scripts.

In this framework, nascent PG was represented as a polymer composed of seven residue types (UNDP, ANAM, BNAG, BNAM, ADGG, MDAP, and DALA), corresponding to undecaprenyl pyrophosphate, α-MurNAc, β-GlcNAc, β-MurNAc, L-alanine–D-isoglutamate, meso-diaminopimelate, and D-alanine, respectively. UNDP, ANAM, BNAG, BNAM, and DALA and their associated linkage patches have direct equivalents in standard CHARMM force fields and were renamed as needed to avoid naming collisions (from UNDPP, AMU2AC, BGLCNA, and BMU2AC).

The ADGG residue was adapted from the CHARMM 3BMDP patch to model the L-alanine–D-isoglutamate dipeptide at the base of the stem and supports extension via isopeptide bonds from the CH2–COO⁻ terminus, by analogy to GLY. The MDAP residue was adapted from published DAP parameters^71^ by converting CHARMM22 atom types to CHARMM36m, using CHARMM LYS and 6CL as analogies.

NPG-derived lipid II and extended chains were validated in short membrane-only simulations to confirm retention of stereochemistry at all glycosidic linkages and planarity of peptide bonds, with no unphysical behavior observed. NPG chains were initialized in a pre-docked configuration using equilibrated FtsWIQLBN–L2–L2 structures to define the coordinates of the undecaprenyl pyrophosphate and the first two glycans; remaining units were built from internal coordinate tables in CHARMM. No NPG-specific restraints were applied beyond standard equilibration, and chains relaxed freely during production runs. In simulations with extended donor substrates (L4 and L32), both donor and acceptor substrates were derived from the NPG parameter set.

### Equilibrium MD simulations of FtsWIQLB systems

Prepared FtsWIQLB system structures were embedded in a 1-palmitoyl-2-oleoyl-sn-glycero-3-phosphoethanolamine (POPE) membrane bilayer and solvated with 150 mM NaCl and, in applicable simulations, 10 mM MgCl2, using the CHARMM-GUI Membrane Builder^68^ or Multicomponent Assembler^72^. Equilibration was performed according to the CHARMM-GUI equilibration protocol involving a series of gradually relaxing sidechain and backbone restraints.

Atomistic MD (MD) simulations were carried out in two stages: system preparation and equilibration with NAMD version 2.14^36^ (Table 2, Sim# 1-37) or GROMACS 2023.3^73^ (Table 2, Sim# 42-43c, 53, 54, 58), with all preparation and equilibration run on the Rockfish cluster at Johns Hopkins University, followed by production runs, which used Anton’s native MD software on Anton 2 (Table 2, Sim# 1-37) and Anton 3^74^ (Table 2, Sim# 42-43c, 53, 54, 58); neither NAMD nor GROMACS was run on Anton. The CHARMM36m force field^75^ was used for protein and lipids, with TIP3P for water molecules (parameters for L2 and its polymeric derivatives described above). The systems were equilibrated in the NPT ensemble (1 atm, 300K or 310K) with positional restraints applied to backbone atoms, followed by 8-52 ns of unrestrained equilibration. CHARMM-GUI protocols were used for both NAMD and GROMACS, which therefore differ mainly in the engine. Each system was energy-minimized and then equilibrated in six stages with progressively weaker position restraints on the protein (1 fs time step for the first three stages and 2 fs for the last three), followed by production with a 2 fs time step. In both engines, nonbonded interactions used a 12 Å cutoff (switching from 10 Å), long-range electrostatics were computed by particle-mesh Ewald, and bonds involving hydrogen atoms were constrained. In NAMD, minimization ran for 10,000 steps, temperature was controlled by Langevin dynamics (damping coefficient 1 ps^- 1^), and pressure by the Langevin piston method (period 50 fs, decay 25 fs) from the third stage onward. In GROMACS, minimization used steepest descent (up to 50,000 steps; force tolerance 1000 kJ mol^-1^ nm^-1^), temperature was controlled by the v-rescale thermostat, pressure by semi-isotropic C-rescale coupling, and hydrogen-bond constraints by LINCS.

Independent simulation replicates were generated by initiating production runs from the same equilibrated coordinates with randomized initial velocities (i.e., distinct Maxwell– Boltzmann velocity assignments). For Anton 2 trajectories (e.g., Sim# 14b and 15b), velocities were randomized at the start of the NAMD preproduction stage (minimization/equilibration/8 ns unrestrained MD), followed by 1 µs production on Anton 2. For Anton 3 trajectories (e.g., Sim# 43b and 43c), preproduction was done in GROMACS, velocities were then randomized immediately prior to Anton 3 submission, and production runs were performed on Anton 3.

For FtsWIQLBN–L2–L2, a 1 µs production trajectory was generated on Anton 2 and subsequently extended by 3 µs on Anton 3 under identical force-field parameters and thermodynamic conditions. Owing to different default output intervals (0.24 ns per frame on Anton 2; 1.2 ns per frame on Anton 3), the combined trajectory contains nonuniform frame densities. Time-resolved binary state analyses were constructed from contiguous state intervals and therefore preserve the temporal behavior of the full 4 µs trajectory without subsampling.

Production simulations on Anton, using Anton’s native MD software, were performed with a timestep of 2.5 fs (A2) or 2.0 fs (A3). Bond lengths for hydrogen atoms were constrained using the M-SHAKE algorithm. An r-RESPA integrator was used. Long-range electrostatics were computed every 3 steps using the k-space Gaussian split Ewald method. Atomic coordinates were written at an output frequency of 0.24ns/snapshot (A2) or 1.2ns/snapshot (A3). Production MD simulations on Anton were carried out for 1-4 µs. Each trajectory was unwrapped using the PBCTools plugin of VMD and aligned to the backbone atoms of FtsW to facilitate comparison across complexes. Code for analyzing simulations was written using the MDAnalysis python package (2.8.0).

### Metadynamics Simulations

To carry out enhanced sampling of L2 insertion into the donor site, well-tempered metadynamics simulations were performed at 300K using GPU-accelerated NAMD. Two collective variables (CVs) were defined: (1) distance between the center of mass (COM) of TM6 and TM7, and (2) distance between the COM of the C55 tail of L2 and the COM of FtsW TM8–FtsI TM interface. Gaussian biasing potentials were deposited every 2 ps with an initial height of 0.2 kcal/mol, a width of 0.1 Å, and bias factor of 10. Simulations were run until at least one full insertion event was observed and the C55-protein interface distance fell below 25 Å. The resulting biased trajectory (2-mtd3) was 188 ns long.

Metadynamics output PMFs were analyzed using custom Python scripts. For visualization, the surface was discretized into stepped contour levels and regions outside the sampled/accessible domain were masked (defined as grid points with finite PMF values within the plotted energy range). To estimate the lowest-energy path (LEP) connecting two configurations on the PMF surface, we treated the 2D PMF grid as an 8-neighbor graph and computed a minimum-cost path using Dijkstra’s algorithm. Endpoints were specified in CV space and mapped to the nearest grid points. The per-step cost was defined as a weighted sum of geometric step length in CV space and the PMF value at the destination node (cost =*w*_dist_Δ*s* + *w*_*E*_*F*), enabling a tunable tradeoff between path smoothness and energetic preference. A one-dimensional free-energy profile was extracted by plotting *F* along the path as a function of cumulative arc-length (path progress).

### Analysis of Structural Dynamics

Trajectory analyses were performed using MDAnalysis^62^ Python library and custom Python scripts. Root-mean-square deviation (RMSD), distance between catalytic residues and L2 atoms, and dihedral angle distributions (e.g., Ψ angle of R137) were computed over the course of the simulation. The number and persistence of FtsWI-PG salt bridges, and electrostatic coordination of PG pyrophosphates were calculated with a 3.5 Å distance cutoff between acidic and basic atoms. Surface contact areas were calculated using a custom script based on the get_area command in PyMOL, with parameters dot_solvent, dot_density, and solvent_radius set to 1, 4, and 1.4, respectively. The surface contact area between FtsW and the PG sugar groups was determined by calculating and adding the solvent exposed surface area (SASA) for each component individually, then subtracting the SASA of the whole complex.

### Plasmid and Strain Construction

Plasmids used in this study (Supplementary Table 2) were assembled by Polymerase Chain Reactions (PCR) amplifying insert and vector DNA fragments followed by In-Fusion cloning (Takara Biosciences, In-Fusion HD Cloning Kit). Oligonucleotides (Integrated DNA Technologies) used in PCR amplifications are described in Supplementary Table 3. All plasmids were verified by DNA sequencing. After construction, electroporation was used to transform plasmids to create strains of interest (Supplementary Table 1) under appropriate antibiotic selection.

### In vivo functionality of FtsW(SSCC) and treatment of cells with DTT

Strains AP183 and AP760 were struck from frozen glycerol stocks onto Luria Bertani Agar (LB) plates supplemented with carbenicillin (Carb; 60 µg/mL), chloramphenicol (Chlor; 25 µg/mL), and 0.2% arabinose, and incubated approximately 16–18 h at 37°C. Single colonies were streaked onto LB+Carb plates containing 100 µM IPTG and grown approximately 16–18 h at 37°C.

The following morning, colonies were inoculated into 3 mL EZRDM (EZ Rich Defined Media) +Carb+100 µM IPTG and grown at 30°C with 240 rpm shaking to OD_600_ ≈ 0.5. Aliquots (100 µL) were spotted onto filter disks on LB+carb plates containing 100 µM IPTG ± 10 mM DTT, then incubated approximately 16–18 h at room temperature. Cells were spotted onto plates rather than grown in liquid culture because DTT-containing liquid medium did not support growth. Cells were resuspended in 1× PBS (phosphate-buffered saline), fixed in 4% paraformaldehyde (PFA), washed twice in 1× PBS, concentrated by centrifugation, and imaged by brightfield microscopy on 3% agarose pads in PBS.

### In vivo functionality of FtsW(S382C) and inhibition of mutant with MTSES

Strains AP183 and AP766 were struck from frozen glycerol stocks onto LB plates supplemented with Carb, Chlor, and 0.2% arabinose, and incubated approximately 16–18 h at 37°C. Single colonies were streaked onto LB+Carb plates containing 100 µM IPTG and grown approximately 16–18 h at 37°C.

The following morning, colonies were inoculated into 500 µL LB+Carb+100 µM IPTG and grown at 37°C with 240 rpm shaking for 2 h to OD_600_ = 0.5–1.0. Cultures were diluted 200-fold into EZRDM+Carb ± 50 µM IPTG, ± 100 µM MTSES (Sodium (2-sulfonatoethyl) methanethiosulfonate) and grown at 37°C with shaking to OD_600_ ≈ 0.4. Cells were pelleted by centrifugation, concentrated and resuspended in EZRDM, and immediately imaged by phase-contrast microscopy on 3% agarose pads in EZRDM.

### In vivo functionality of the FtsW(S130C) and FtsW(S136C) single mutants

Strains AP30 (BW25113, no plasmid, grown without antibiotics), AP183 (FtsW-Halo), AP756 (FtsW(S130C)-Halo), AP758 (FtsW(S136C)-Halo) and AP760 (FtsW(SSCC)-Halo) were grown and imaged as described above for FtsW(S382C), except that MTSES was omitted and cultures were diluted into EZRDM+Carb with 50 µM IPTG and no DTT, grown at 37°C to OD600 ≈ 0.4, and imaged by phase-contrast microscopy. Cell lengths were measured as described in “Cell-length quantification and statistics”.

### In vivo functionality of FtsW(L198C) and inhibition of mutant with MTSES

Strains XY396 and AP852 were struck from frozen glycerol stocks onto LB plates supplemented with Carb, Chlor, and 0.2% arabinose for the induction of the WT FtsW from a plasmid, and incubated approximately 16–18 h at 37°C. Single colonies were inoculated into 3 mL LB+Carb+50 µM IPTG for the induction of the FtsW variant from the complementation plasmid and grown to saturation (∼8 h) at 37°C with 240 rpm shaking.

Saturated cultures (2 µL) were inoculated into 3 mL M9 (minimal media)+Carb ± 100 µM IPTG, ± 100 µM MTSES and grown at 25°C to OD_600_ = 0.15–0.4 (∼20 h) with shaking. 500 µL of cells were pelleted by centrifugation, gently resuspended in 50 µL M9, spotted onto 3% agarose pads in M9, and immediately imaged by phase-contrast microscopy.

### Dominant Negative Characterization of FtsW-Halo mutant strains

AP892, AP893, AP894, AP895, and AP896 were streaked from frozen glycerol stocks onto LB agar plates supplemented with carbenicillin (Carb; 60 µg/mL) and incubated for 16– 18 h at 37°C. The following morning, individual colonies were inoculated into 1 mL EZ Rich Defined Medium (EZRDM) supplemented with Carb and 100 µM IPTG and grown at 37°C with shaking at 240 rpm until reaching an OD600 of 0.2–0.5. Cultures were subsequently diluted into EZRDM to an initial OD600 of 0.002 and grown in the presence or absence of 100 µM MTSES for 3.5 h, or until reaching an OD600 of approximately 0.4. Cells were then collected, concentrated, and spotted directly onto 3% agarose pads for immediate phase-contrast microscopy.

For fluorescence microscopy, concentrated cells of AP892, AP893, AP894 + MTSES, or AP896 + MTSES were labeled with 1 µM JF646-HaloTag ligand (Promega) for 15 min at 37°C with shaking. Cells were subsequently washed four times with 500 µL EZRDM to remove unbound ligand and concentrated prior to imaging. Cells were spotted onto 3% EZRDM agarose pads and immediately imaged by phase contrast and fluorescence microscopy.

Quantification of fluorescence intensity from individual cells expressing fluorescently tagged proteins was performed using MicrobeJ^76^.

### Cell-length quantification and statistics

Cell lengths were measured manually in Fiji (ImageJ) using the segmented-line tool and converted to micrometres using the pixel size recorded for each imaging day. Each experiment comprises three independent biological replicates, except where noted in the figure captions; *n* refers to biological replicates throughout, and cell numbers are given in the Source Data.

Cell lengths are displayed as SuperPlots^77^: individual cells are coloured by biological replicate, replicate medians are superimposed, and the bar marks the geometric mean of the replicate medians ± s.e.m. Cells were scored as elongated when their length exceeded a threshold set for each panel at twice the median length of that panel’s control group, with imaging days pooled for the control (dashed line in the figures); this is an operational criterion based on a cell that has failed one round of division being about twice the normal division length. The percentage of elongated cells was calculated separately for each biological replicate and is reported as the mean of the replicate percentages with the range across replicates (n = 3 biological replicates unless stated; n = 2 for the WT control groups in the S382C experiment). As a complementary descriptive summary, differences between conditions are also expressed as the ratio of medians in the Supplementary Tables. Intervals are the 2.5th and 97.5th percentiles of 40,000 bootstrap ratios (test group / reference group). In each iteration, cells were resampled with replacement within every replicate, replicates present in both groups were then resampled with replacement, and the ratio was taken of the geometric means of the resulting replicate medians (random seed 11). A ratio of 1 means the medians are the same. Only replicates present in both groups were used, so a replicate that one group lacks was dropped from that comparison. A replicate drawn twice in one iteration carries the same cell resample, which makes the interval conservative. These intervals are descriptive and are not used as significance tests. Analyses were performed in Python 3.11 using NumPy and SciPy.

### Single-molecule tracking and data analysis

Single-molecule tracking (SMT) was performed using a similar optical setup as previously described, with an Olympus IX-81 inverted microscope equipped with a 1.45 NA ×100 objective and an Andor DU888 camera in epifluorescence-illumination mode. Imaging was performed using a 646-nm laser. The focal plane was positioned approximately 0.3 µm above the position of maximum cell contrast to image molecules localized near the bottom half of the cylindrical portion of the cell body. Cells expressing proteins from the uninduced low-expression pIQ-HT plasmid series were labeled with the JF650X HaloTag ligand (Promega). JF650X was used because of its increased brightness and photostability relative to JF646.

Before imaging, cells were grown overnight in LB medium containing the appropriate antibiotic and subsequently inoculated into M9 medium. Cells were harvested during exponential growth at an OD600 of 0.1–0.5. Cells were concentrated by centrifugation at 5,000 × g for 2 min and resuspended in 100 µl M9 medium. Cells were labeled with 0.4 µl of 16 µM JF650X dye at final concentration of 64 nM and incubated for 10 min with gentle rocking. Following labeling, cells were washed four times with M9 medium by centrifugation at 5,000 × g for 2 min and gentle resuspension. After the second wash, cells were transferred to a fresh microcentrifuge tube to minimize carryover of dye adsorbed to the tube. Following the third wash, the OD600 was measured and used to determine the volume for the final resuspension. After the fourth wash, cells were concentrated to an OD600 of approximately 1-2 and loaded onto 30 µL agarose pad prepared with M9 medium within a Geneframe imaging chamber (1.5 x 1.6 cm). Approximately 0.8 µl of concentrated cells was applied to the agarose pad and allowed to partially dry before imaging.

Single molecules were continuously imaged using a 500-ms exposure time with a 1:10 neutral-density filter and 7 mW output power from the 646-nm laser. These illumination conditions were reduced from those used previously for JF646 because JF650X provided increased brightness and photostability. The relatively slow imaging rate was used to reduce the contribution of rapidly diffusing molecules and to facilitate tracking of processive membrane-associated molecules.

### Mass spectrometry of HaloTag-enriched FtsW

Cell growth and harvest. Strains AP892, AP893, AP894 and AP896 (Supplementary Table 1) were inoculated from streaked single colonies into LB containing 50 µg/mL carbenicillin and 500 µM IPTG and grown at 37 °C for 1–2.5 h. Starter cultures were used to inoculate 100 mL LB containing 50 µg/mL carbenicillin and 500 µM IPTG to a calculated starting OD600 of 0.003. Where indicated, MTSES was added at the time of inoculation to a final concentration of 100 µM and was present throughout growth. Cultures were grown at 37 °C with shaking and harvested at OD600 ≈ 0.25–0.40. Cells were held on ice for 20 min, pelleted at 4,100 × g for 20 min at 4 °C, resuspended in 1 mL ice-cold PBS, pelleted again and frozen on dry ice.

### HaloTag Affinity Enrichment and Differential Cysteine Alkylation of FtsW

Cell pellets (AP892, AP893, AP894, AP896) were resuspended on ice in 1 mL chilled lysis buffer containing 50 mM Tris-HCl (pH 7.6), 150 mM NaCl, and 0.5% n-dodecyl-β-D-maltoside (DDM) (Thermo fisher). Cells were disrupted by sonication on ice at 55% amplitude using alternating 5-s on and 5-s off pulses for 5 min. Lysates were clarified by centrifugation at 15,000 × g for 15 min at 4 °C. Protein concentrations in the supernatants were determined using a bicinchoninic acid (BCA) assay and adjusted to 0.1 mg/mL with lysis buffer.

For affinity enrichment, 50 µL of HaloLink resin suspension (Promega) per sample was pelleted at 800 × g for 1 min and washed three times with 400 µL of binding buffer containing 50 mM Tris (pH 7.6), 150 mM NaCl, and 0.2% DDM. The resin was resuspended in 50–100 µL of binding buffer and incubated with 1 mL of normalized lysate for 30–60 min at room temperature with rotation. Following centrifugation at 800 × g for 2 min, the resin was washed three times with 1 mL of binding buffer.

Resin-bound proteins were incubated in 250 µL of resuspension buffer (8 M urea, 100 mM ammonium bicarbonate, 0.2% DDM). Free thiols were alkylated with 4-vinylpyridine at a final concentration of 45 mM for 30 min at 37 °C with mixing at 700 rpm. The resin was pelleted at 800 × g and washed three times with 500 µL of binding buffer. The resin was then resuspended in 250 µL of resuspension buffer, and disulfide bonds were reduced by adding 4.5 µL of 700 mM dithiothreitol (DTT) and incubating for 30 min at 37 °C. The resin was pelleted at 800 × g and washed with 500 µL of binding buffer. Newly exposed thiols were alkylated by adding 9 µL of 700 mM iodoacetamide (IAA; final concentration 40 mM) and incubating for 45 min in the dark with rotation.

### Proteolytic Digestion and Peptide Cleanup

Before digestion, the urea concentration was adjusted to 2 M with 100 mM ammonium bicarbonate. Each sample received 1 µg of trypsin and 0.5 µg of chymotrypsin and was incubated for 8 h with rotation at 37 °C. Digests were acidified with trifluoroacetic acid (TFA) to a nominal final concentration of 1% and desalted using Sep-Pak Vac 1cc (50mg) C18 cartridges (Waters). Cartridges were conditioned twice with 1 mL of elution buffer (80% acetonitrile and 0.1% TFA in water) and equilibrated four times with 1 mL of aqueous 0.1% formic acid. Samples were loaded slowly, and cartridges were washed four times with 1 mL of aqueous 0.1% formic acid. Peptides were eluted with 1 mL of elution buffer, with eluate collection assisted by centrifugation at 300 rpm for 5 min in an Eppendorf 5910R centrifuge. Peptides were dried by speed vac and stored at -80°C until injection.

### Mass Spectrometry Data Acquisition

Peptide samples were resuspended in 0.1% formic acid (Optima, Fisher) in LC grade water (Optima, Fisher) to a final concentration of 1 μg/μL. Approximately 1μg of peptides were injected into a Thermo UltiMate 3000 UHPLC system for chromatographic separation. The column temperature was maintained at 40 °C, and the flow rate was 0.300 μL/min for the duration of the run. Solvent A (0.1% formic acid (FA)) and Solvent B (0.1% FA in ACN) were used as chromatography solvents. Peptides were allowed to accumulate onto the trap column (Acclaim PepMap 100, C18, 75μm x 2cm, 3μm, 100Å column) for 10 min (during which the column was held at 2% Solvent B). The peptides were resolved by switching the trap column to be in-line with the separating column (Acclaim PepMap RSLC, C18, 75μm x 2cm, 2μm, 100Å column), increasing the gradient to 5% B over 15 min and then applying a 47 min linear gradient from 5% B to 25% B. Then the gradient was increased linearly to 40% B over 13 minutes. Next, the gradient increased again from 40% B to 90% B over 5 min. The gradient was held at 90% B for 5 min. The column was then cleaned with a sawtooth gradient to purge residual peptides between runs in a sequence. A one-hour blank cleaning sequence was also run between every 5 samples to ensure a clean column. A Thermo Orbitrap Ascend mass spectrometer was used to analyze protein digests. A full MS scan in positive ion mode was followed by 20 data-dependent MS scans. The full MS scan was collected using a resolution of 120000 (m/z 200), an AGC target of 6E5, a maximum injection time of 64 ms, and a scan range from 350 to 1500 m/z. The data-dependent scans were collected with a resolution of 30000 (m/z 200), an AGC target of 1E5, and an isolation window of 8 m/z units. To dissociate precursors prior to their reanalysis by MS2, peptides were subjected to an HCD of 30% or 32% normalized collision energies. Fragments with charges of 8 or higher, and unassigned were excluded from analysis, and a dynamic exclusion window of 30.0s was used for the data-dependent scans.

### FragPipe Analysis

For analysis of the mass spectrometry data, FragPipe (version 24.0), MSfragger (version 4.4.1), IonQuant (version 1.11.20), Philosopher (version 5.1.3), and diaTracer (version 2.2.1) were used. The LFQ-MBR template was applied, and the search database included ftsW-halo, or its mutants (SSCC, S382C or L198C), with decoys and contaminants generated by FragPipe. Methionine oxidation, N-terminal clipping, and N-terminal acetylation, cysteine pyridylethylation, cysteine carbamidomethylation and cysteine +139.96 Da (modified by (2-Sulfonatoethyl)methanethiosulfonate) were specified as dynamic modifications. The false discover rate is set to 0.05 for IonQuant.

Across WT and mutant samples with and without MTSES, peptides and the four native cysteines were identified at various levels with modifications (purple), but none of the engineered cysteine positions or their native positions (except one) were recovered. In FtsW(L198C)-Halo, 109 peptides were identified covering 63.1% of the fusion, comparable to wild-type FtsW-Halo (107 peptides, 64.9%). FtsW(SSCC)-Halo recovered 28 peptides covering 24.7% of the fusion. A peptide spanning position 382 was identified in wild-type FtsW-Halo and in FtsW(L198C)-Halo, where that position is a serine, but not in the FtsW(S382C)-Halo sample. Peptide recovery across FtsW was lower in the three treatment samples, falling from 105–109 peptides in the untreated variants to 7–28 in the treated and double-cysteine samples. The preparative cultures required for mass spectrometry were grown under harsher conditions than this study’s imaging and phenotypic assays: cells were induced at 500 µM IPTG, five-fold above the level used throughout this study, and MTSES was present from inoculation through the entire growth rather than for a defined treatment window (SI Methods). FtsW(L198C) with MTSES and FtsW(SSCC) are already dominant negative at 100 µM IPTG, so prolonged growth under stronger induction could have compromised cell physiology and reduced the yield of intact FtsW-Halo. Modified or disulfide-locked FtsW may also partition into poorly soluble complexes lost during lysis and affinity capture.

### AiPP-LigBind analysis

Residue-level ligand-binding propensity was predicted using AiPP-LigBind as described previously^55^. The configuration used here employed per-residue representations from hidden layer 76 of ESM-C 6B (esmc-6b-2024-12)^56^ and the previously described 200-member LigBind ensemble. The final LigBind score for each residue was calculated as the arithmetic mean of the sigmoid-transformed outputs across the ensemble and was interpreted as a relative ligand-binding propensity rather than a calibrated binding probability.

LigBind was first applied to the 414-residue consensus FtsW sequence, and scores were mapped onto the same consensus structural model used for conservation visualization. The predefined E. coli-referenced donor/catalytic/activation residues R137, R243, R246, and D297; gating residues L198, L236, and K370; and acceptor-pocket/clamp residues R84, W138, R145, Y379, and S382 were annotated. Among the 20 highest-scoring consensus residues, non-mechanistic residues within ±2 positions in primary sequence of these sites were additionally identified.

For the representative-ortholog analysis, LigBind was applied independently to FtsW from E. coli (P0ABG4), P. aeruginosa (Q9HW00), B. subtilis (O07639), C. crescentus (B8H092), and K. pneumoniae (A0A378ACG3). Residue correspondences followed the ortholog-specific mappings used in the manuscript and Supplementary Information, supplemented by local sequence correspondence where necessary.

For the family-wide analysis, LigBind was applied independently to each of the 6,859 ungapped FtsW sequences and the predictions were mapped back to the multiple-sequence alignment and the E. coli-referenced scaffold. Exact-position recurrence was calculated as the fraction of orthologs containing an unambiguous residue at an aligned position in which that residue ranked among the 20 highest-scoring LigBind predictions. As a length-normalized robustness comparison, recurrence was also calculated using the highest-ranking ⌈0.05L⌉ residues in each sequence, where L is the ungapped sequence length. Local sequence-neighborhood recurrence was evaluated within ±2 positions of each predefined mechanistic site in the ungapped primary sequence; the center residue and all other predefined mechanistic residues were excluded from the adjacent-only analysis. Sequence-neighborhood criteria refer only to primary-sequence adjacency and do not imply three-dimensional proximity.

## Code and data availability

### Code availability

AiPP-LigBind is described previously^55^. Custom scripts for generating the FtsW consensus sequence, mapping LigBind predictions to the multiple-sequence alignment, calculating family-wide recurrence, and generating the associated analyses are available at Zenodo: https://doi.org/10.5281/zenodo.22116961. The MD analysis code (FtsW Dynamics) is available at Zenodo: https://doi.org/10.5281/zenodo.23989670, and the nascent PG parameterization code (Nascent PG Builder) at https://doi.org/10.5281/zenodo.24014488. The cell-length and fluorescence analysis code is deposited with the corresponding measurements at https://doi.org/10.5281/zenodo.24014699.

### Data availability

Source data underlying the consensus conservation and LigBind analysis, representative-ortholog analysis, and family-wide recurrence calculations are available at Zenodo: https://doi.org/10.5281/zenodo.22116961. The MD trajectories (FtsW Dynamics) are available at Zenodo: https://doi.org/10.5281/zenodo.23989670, and the cell-length and fluorescence measurements at https://doi.org/10.5281/zenodo.24014699. The deposited trajectories have water, ions and the membrane removed, so analyses that depend on solvent or lipid coordinates cannot be reproduced from them directly.

## Notes

### Competing Interest Statement

The authors have declared no competing interest.

### Summary of Updates

Revised in response to peer review. Author list and affiliations updated. Added molecular dynamics simulations of the engineered cysteine mutants (new Figure 4). Added single-molecule tracking of FtsW(L198C and S382C), mass spectrometry of HaloTag-enriched FtsW variants, and cell-length quantification with statistics. Added ligand-binding propensity (AiPP-LigBind) analyses across FtsW orthologs. Expanded the Discussion. Text clarified throughout; Methods, figures, Supplementary Information and data/code availability updated.

### Funding

- National Institute of Health (NIH): 2R35GM136436, F32GM143895, F32GM150262, F31GM146442
- Pittsburgh Supercomputing Center (PSC): MCB230000P, MCB230000P

