## Supplementary for "Substrate binding and activation mechanism of the essential bacterial septal cell wall synthase FtsW"

**Supplementary Information**

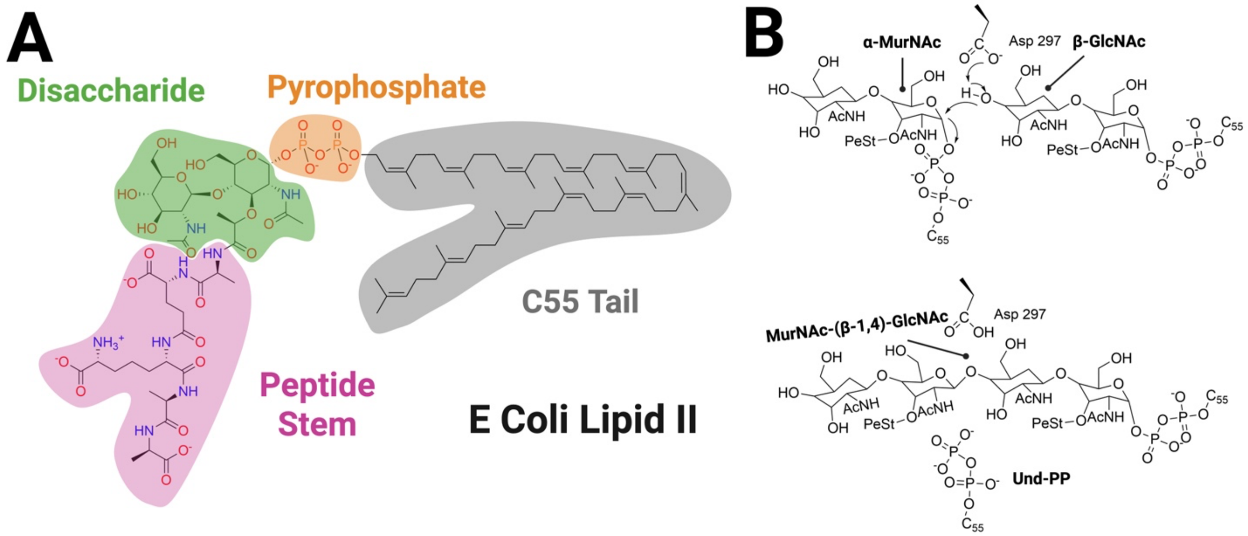


**Figure S1. Lipid II architecture and proposed SEDS glycosyltransferase reaction.**

**(A)** Chemical structure of *E. coli* lipid II highlighting the disaccharide headgroup (MurNAc–GlcNAc; green), pyrophosphate linker (orange), pentapeptide stem (magenta), and the C55 undecaprenyl tail (grey).

**(B)** Schematic representation of the FtsW glycosyltransferase (GTase) reaction, shown using the SN2-like inverting mechanism proposed for the SEDS enzymes. The putative catalytic aspartate (Asp297 in *E. coli* FtsW; Asp262 in RodA) deprotonates the acceptor GlcNAc 4'-OH, enabling nucleophilic attack on the donor MurNAc anomeric carbon (C1) to form the β-1,4 glycosidic bond and release the donor undecaprenyl-pyrophosphate carrier (Und-PP).

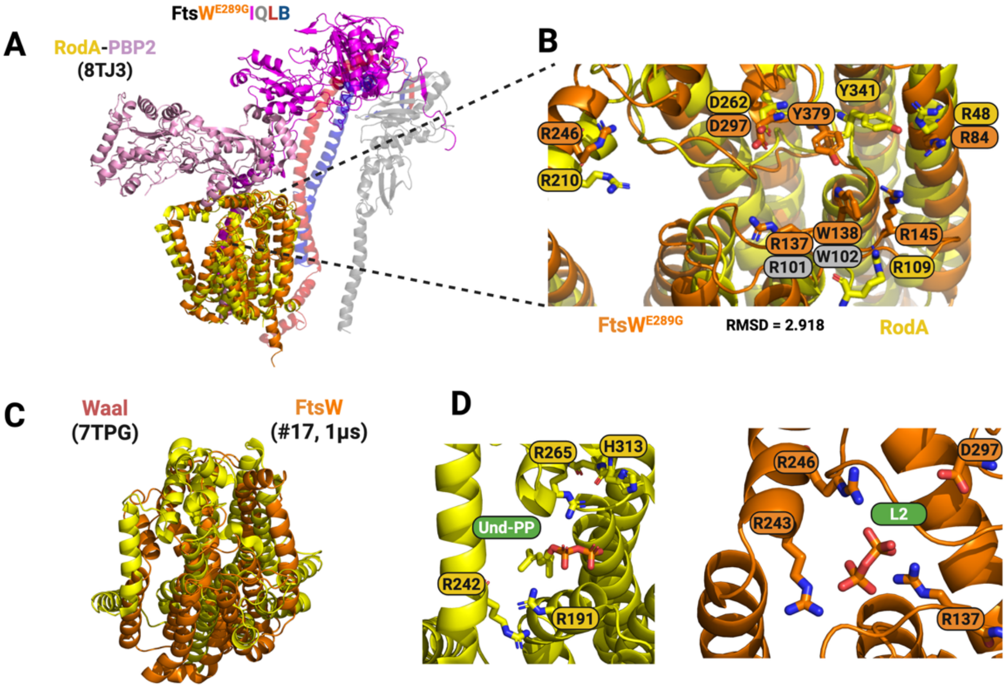


**Figure S2. Structural Comparison of *E. coli* FtsW, *E. coli* RodA, and *C. metallidurans* WaaL.**

(**A**) Structural alignment of FtsW to RodA, using the final snapshot of a 1μs apo FtsW$^{E289G}$IQLB trajectory[1] and the cryo-EM structure of RodA-PBP2[2] as the source of atomic coordinates and the GTases as the alignment selections. Structural alignment of FtsW$^{E289G}$ with RodA (PDB: 8TJ3) yields an RMSD of 2.918 Å. (**B**) Superimposed structural view of the FtsW and RodA catalytic grooves, with key residues shown as sticks with atomic colors. RodA residues not resolved in the cryo-EM structure are marked with gray labels. (**C**) Structural alignment of FtsW to WaaL, using the final snapshot of a 1μs FtsWIQLB + FtsN$^{58-108}$ trajectory (sim 17) and the cryo-EM structure of WaaL as the source of atomic coordinates, and the donor site ligands (Und-PP and L2) as the alignment selections. Because this alignment is defined by the ligands rather than by the proteins, no protein RMSD is reported for this comparison. (**D**) Structural views of the WaaL (red) and FtsW (orange) donor sites, with key residues involved in ligand coordination or catalysis shown as sticks.

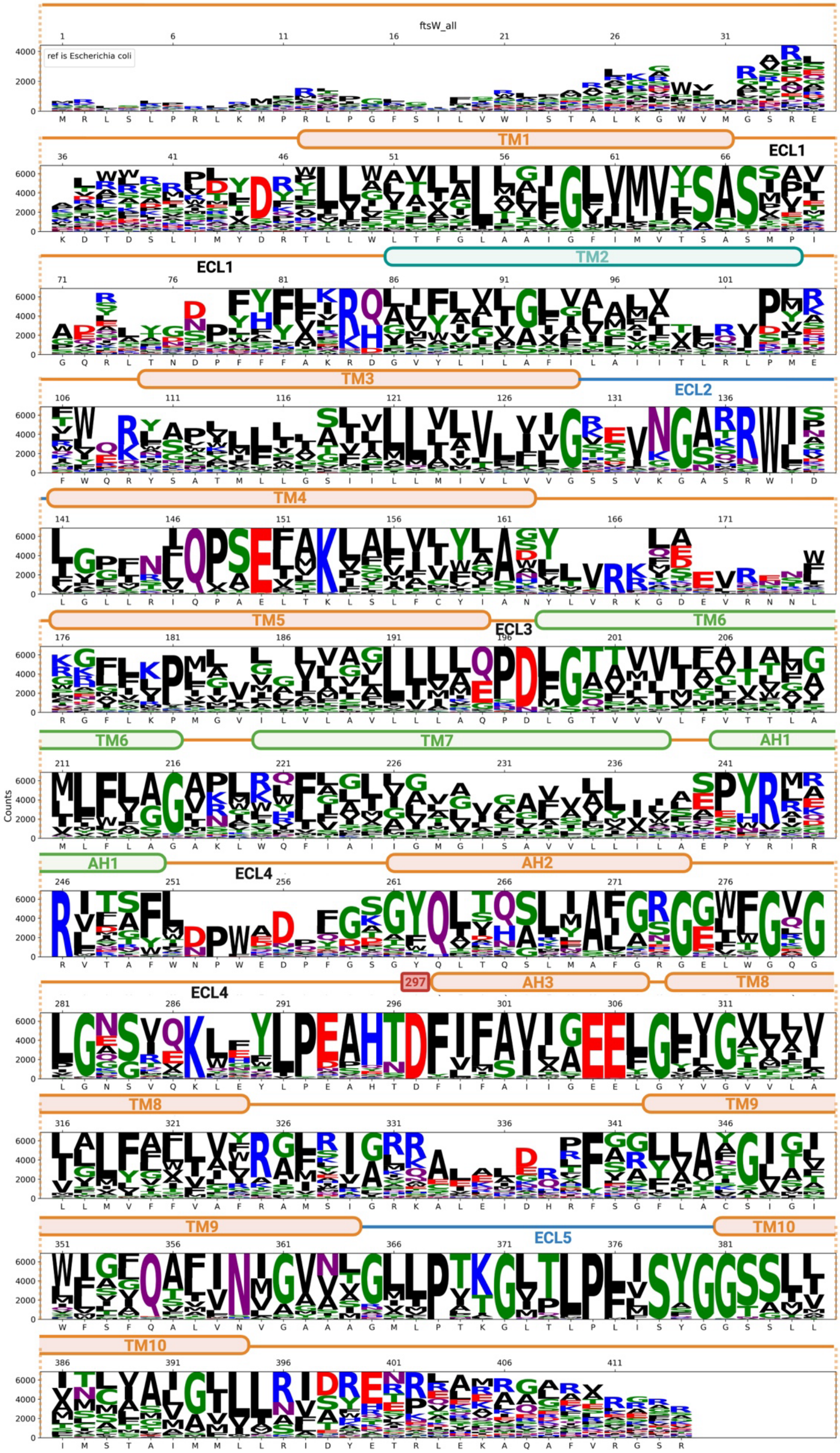

ftsW_all
ref is Escherichia coli
Counts
TM1
ECL1
TM2
TM3
ECL2
TM4
TM5
ECL3
TM6
TM7
AH1
ECL4
AH2
297
AH3
TM8
TM9
ECL5
TM10

**Figure S3. Sequence conservation of FtsW across 6,859 bacterial genomes.**

Sequence logo representation of a multiple sequence alignment of 6,859 FtsW orthologs, with *Escherichia coli* FtsW used as the reference. Position numbers correspond to the E. coli sequence and the E. coli residue identity is shown beneath each position. Letter height reflects residue conservation (counts), with the most conserved residues shown in larger font. Secondary-structure elements are annotated above the alignment based on the FtsW topology shown in Fig. 1B, including transmembrane helices TM1–TM10, extracellular loops ECL1–ECL5, and amphipathic helices AH1–AH3 within ECL4. Structural features and functionally implicated regions are color-coded as in Fig. 1B: donor-site residues in green, acceptor-site residues in cyan, residues contributing to both sites in blue, and the putative catalytic residue D297 in red. Colors indicate residue class: basic (blue), acidic (red), polar uncharged (green), and hydrophobic (black). Helix boundaries are assigned from the apo simulation ensemble and are approximate: the ends of several transmembrane helices fray over the course of the trajectories, so a residue at the extreme of a helix — for example, L198 at the N-terminal end of TM6, or S382 at the N-terminal end of TM10 adjacent to ECL5 — may be assigned to the helix or to the flanking loop depending on the snapshot.

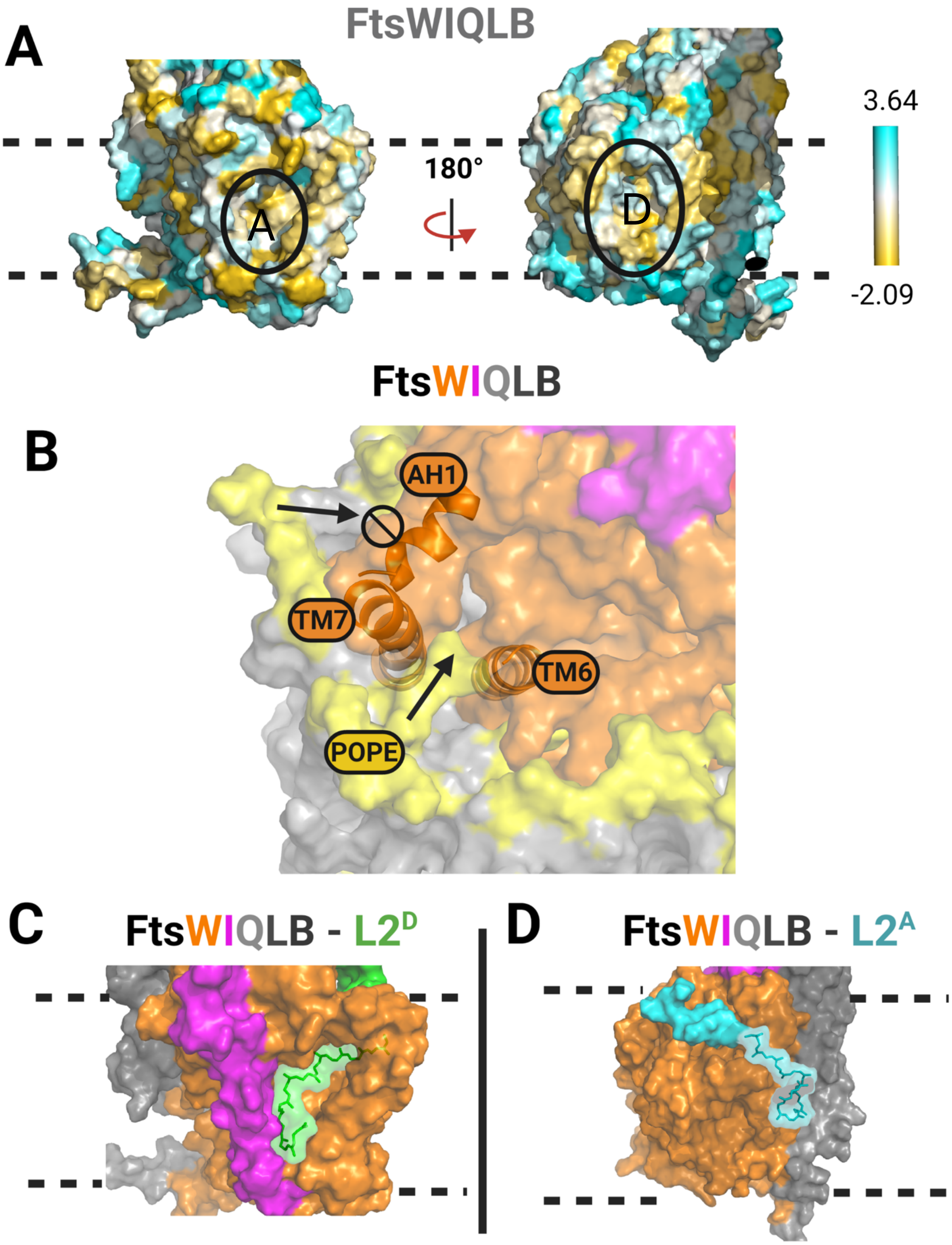


**Figure S4. Hydrophobic cavities and substrate-access pathways in FtsW.**

**(A)** Surface representation of FtsWIQLB colored by residue hydrophobicity on the Wimley–White scale (kcal/mol; gold, most hydrophobic; cyan, most hydrophilic). Two orientations related by a 180° rotation are shown. Dashed lines indicate the approximate membrane boundaries. Membrane-accessible periplasmic cavities are outlined (black ovals, with A and D for acceptor and donor sites, respectively).

**(B)** Amphipathic helix 1 (AH1) forms a barrier on the periplasmic surface that disfavors passage (arrow and circle-backslash symbol) of polar lipid headgroups (yellow). A representative POPE molecule is shown able to access the interior donor cavity via the interface between transmembrane helices 6 and 7 (TM6 and TM7); AH1, TM6, TM7 and POPE are labeled (arrows).

**(C–D)** Docked lipid II (L2) poses in the donor (**C**) and acceptor (**D**) sites, shown with the same viewing orientation and membrane boundaries as in (A). FtsI is colored magenta, FtsW orange, and FtsQLB gray. The L2 sugar headgroup is shown as an opaque surface and the lipid tail as sticks with a transparent surface overlay.

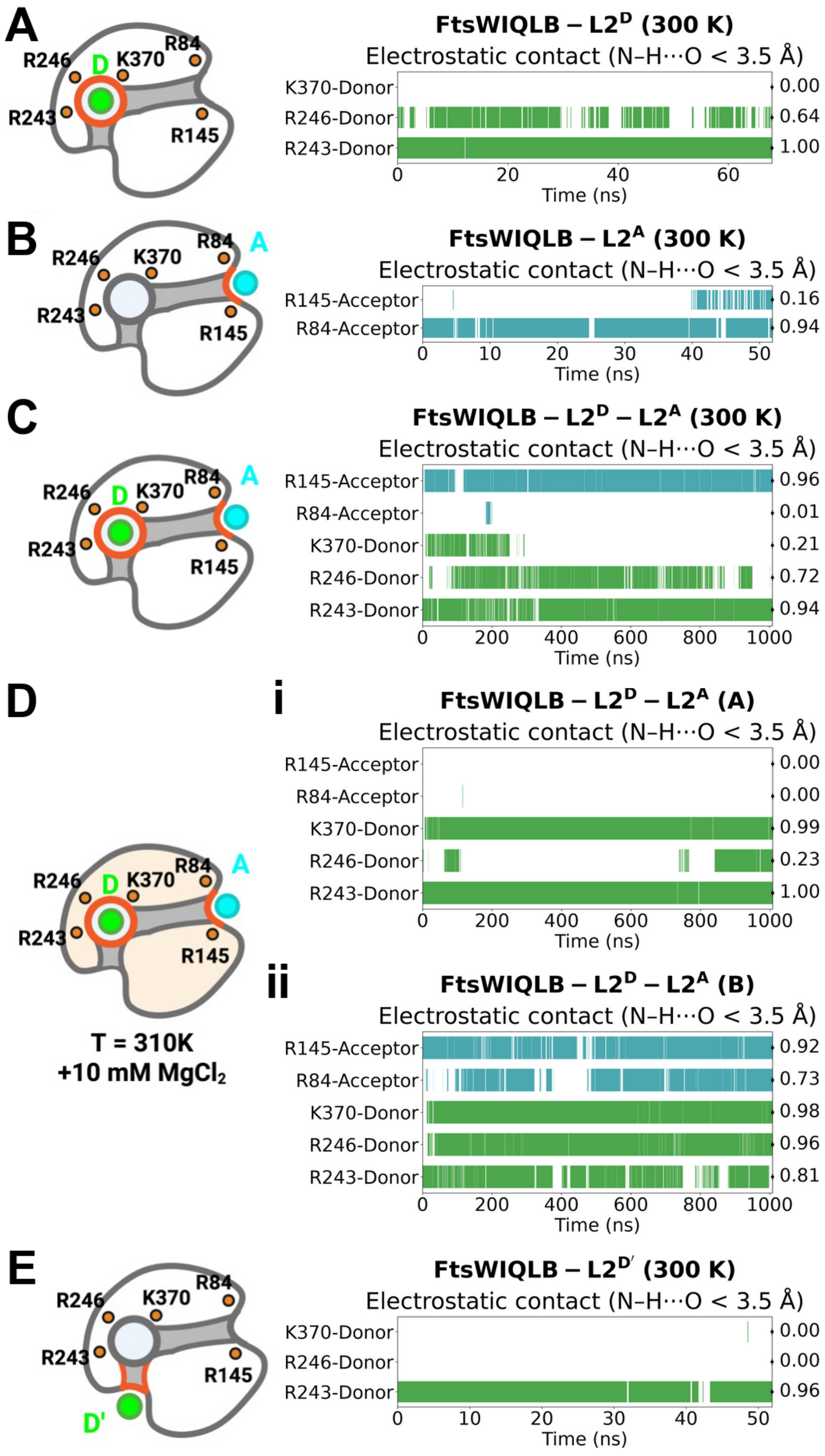
A
R246
D
K370
R84
R243
R145
FtsWIQLB – L2^D (300 K)
Electrostatic contact (N–H⋯O < 3.5 Å)
K370-Donor
R246-Donor
R243-Donor
0.00
0.64
1.00
Time (ns)
B
A
FtsWIQLB – L2^A (300 K)
R145-Acceptor
R84-Acceptor
0.16
0.94
C
FtsWIQLB – L2^D – L2^A (300 K)
0.96
0.01
0.21
0.72
0.94
D
i
FtsWIQLB – L2^D – L2^A (A)
0.00
0.00
0.99
0.23
1.00
ii
FtsWIQLB – L2^D – L2^A (B)
0.92
0.73
0.98
0.96
0.81
T = 310K
+10 mM MgCl₂
E
D'
FtsWIQLB – L2^D' (300 K)
0.00
0.00
0.96

**Figure S5. Electrostatic coordination of lipid II by conserved basic residues in FtsW.**

Binary contact traces report electrostatic interactions between FtsW basic side chains and lipid II pyrophosphate oxygens across equilibrium simulations. Contacts are scored as present when the heavy-atom distance criterion for an N–H···O interaction is met (< 3.5 Å). For each panel, the left schematic indicates the substrate position within the catalytic groove (donor, D; acceptor, A; donor-entry, D′) relative to conserved basic residues, and the right plot shows contact presence over time; values at right report the overall fraction of frames satisfying the contact criterion.

**(A)** Donor-bound state ($L2^{D}$): contacts between the donor pyrophosphate and K370, R246, and R243.

**(B)** Acceptor-bound state ($L2^{A}$): contacts between the acceptor pyrophosphate and R84 and R145.

**(C)** Dual-occupancy state ($L2^{D}$–$L2^{A}$): donor (K370, R246, R243) and acceptor (R84, R145) pyrophosphate contacts over the full trajectory.

**(D)** Dual-occupancy state ($L2^{D}$–$L2^{A}$) at 310K and in the presence of 10 mM MgCl2; two independent replicates.

**(E)** Donor-entry state ($L2^{D'}$): contacts between the donor pyrophosphate and conserved basic residues at the external entry position.

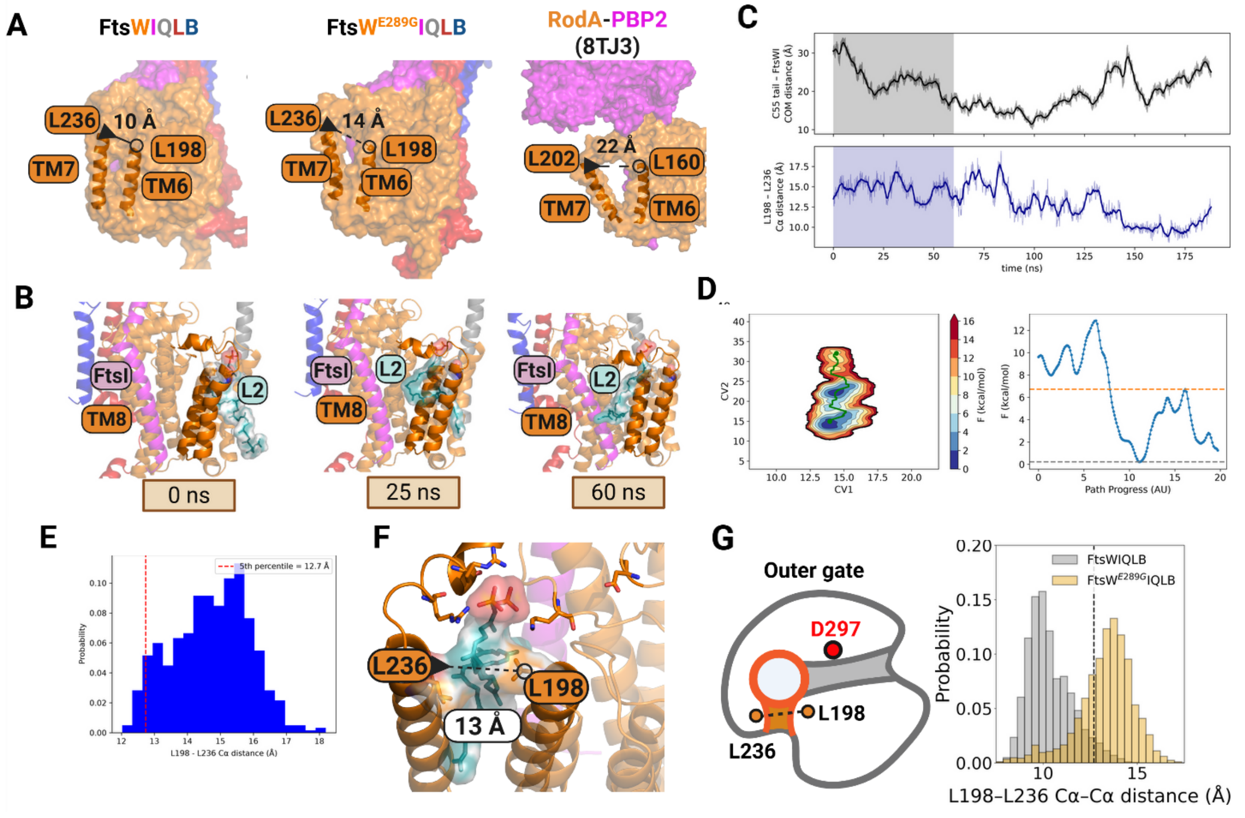


**Figure S6. Metadynamics simulations showing that TM6–TM7 outer-gate opening promotes lipid II access to the FtsW donor cavity.**

**(A)** Structural comparison of the TM6–TM7 outer-gate width, measured as the Cα–Cα distance between FtsW L198 (TM6) and L236 (TM7), shown for WT apo FtsW (left), apo FtsW$^{E289G}$ (middle), and RodA (right) from the RodA–PBP2 cryo-EM structure (PDB 8TJ3; RodA L160 on TM6 and L202 on TM7). Distances are annotated.

**(B)** Representative trajectory snapshots from well-tempered metadynamics illustrating L2 entry through the TM6–TM7 interface, progressing from the external donor-entry position (D') into the donor site (D) at the indicated times.

**(C)** Metadynamics time series showing the lipid II C55 tail position collective variable (top) and the corresponding TM6–TM7 gate distance (L198–L236 Cα–Cα; bottom). The shaded region marks the D'→D transition window.

**(D)** Two-dimensional free-energy surface (PMF) as a function of lipid II tail position and TM6–TM7 separation (left), and the free-energy profile along the lowest-energy path connecting the D' and D basins (right). Dashed lines indicate the lowest energy point during the transition (grey) to the highest energy point (yellow) immediately before full transition into D.

**(E)** Distribution of TM6–TM7 separations sampled during the D'→D transition (0–60 ns). The dashed line denotes the 5th-percentile lower bound (12.7 Å), used as an entry-derived criterion for an "open" outer gate.

**(F)** Transition snapshot illustrating an outer-gate opening of ~13 Å during lipid II entry, with TM6/TM7 and the substrate highlighted.

**(G)** Distributions of TM6–TM7 gate distances from equilibrium simulations of apo FtsWIQLB, comparing WT (gray) and E289G (yellow), showing a shift toward larger openings in E289G. The dashed line marks the entry-derived “open” threshold.

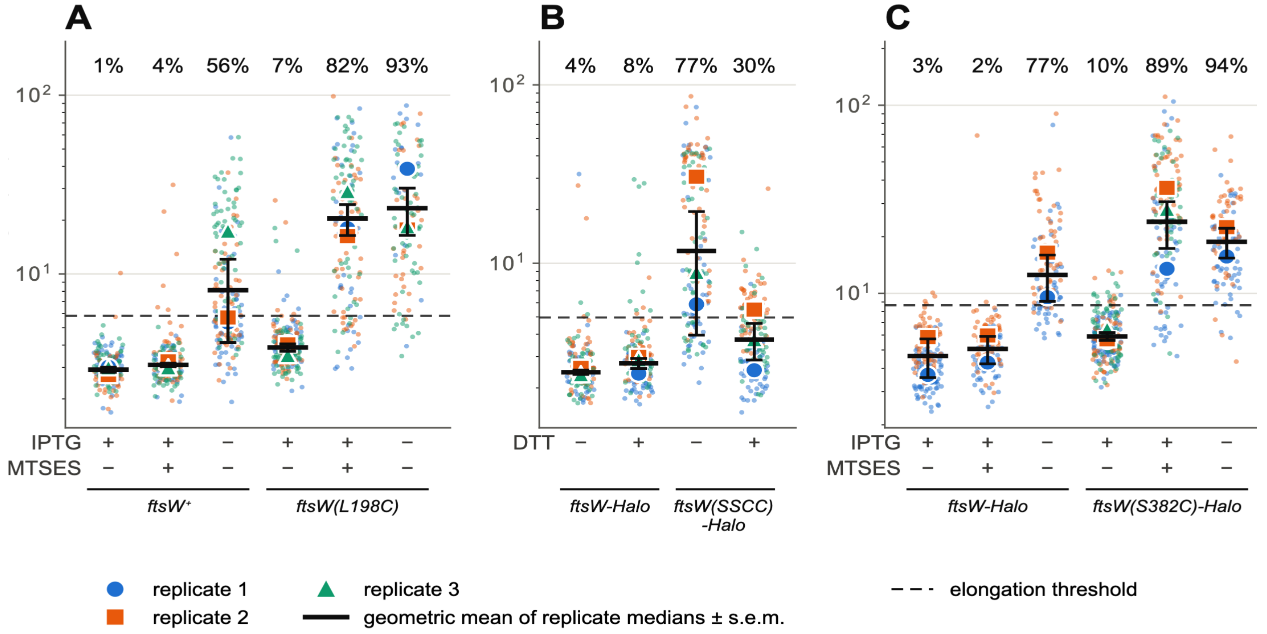


**Figure S7. Cell length measurements of FtsW mutants.**

(**A**) FtsW-depletion strains complemented with FtsW wild-type (FtsW$^+$) or FtsW(L198C), both untagged, grown in M9-glucose at 25 °C, ± 100 μM IPTG, ± 100 μM MTSES. (**B**) Strains expressing FtsW-Halo or FtsW(SSCC)-Halo, grown in EZRDM at 30 °C and then spotted on LB plates containing 100 μM IPTG ± 10 mM DTT and incubated 16–18 h at room temperature before imaging. (**C**) Strains expressing FtsW-Halo or FtsW(S382C)-Halo, grown in EZRDM at 37 °C, ± 50 μM IPTG, ± 100 μM MTSES. Small circles are individual cell data points, colored by biological replicate; circles, squares and triangles are the medians of replicates 1, 2 and 3; the black bar is the geometric mean of the replicate medians ± s.e.m. Note the logarithmic axis. The dashed line marks the elongation threshold, defined for each panel as twice the median length of its control group (the first group in the panel, imaging days pooled), and the percentage above each group is the mean of the replicate percentages of cells longer than this threshold. Measurements were from three independent biological replicates, except the three FtsW-Halo groups and FtsW(S382C) −IPTG in (C), which comprised two. Group summaries, pooled medians, per-replicate percentages and the bootstrap comparisons are in Supplementary Table 4 (A), Supplementary Table 6 (B) and Supplementary Table 9 (C).

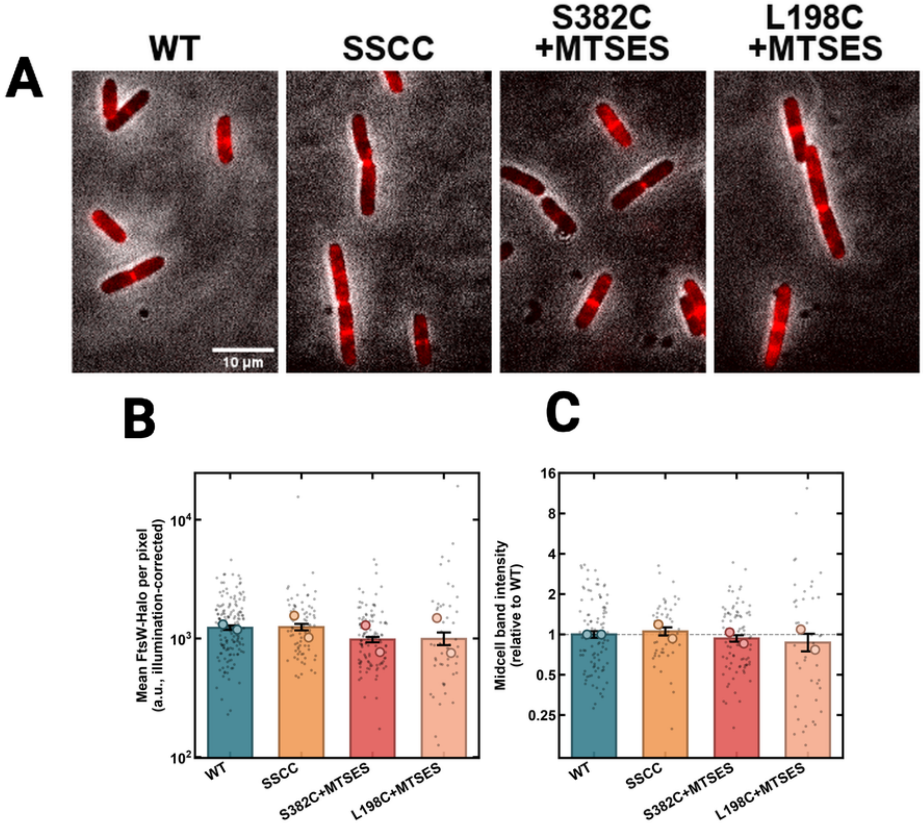


**Figure S8. FtsW-Halo variants localize to the midcell in BW25113 merodiploid cells.**

Cells grown in EZRDM (rich-defined medium) at 37°C were induced with 100 μM IPTG for 5.5 hours and incubated with or without 100 μM MTSES for 3.5 hours prior to preparation for imaging. Cells were labeled with a saturating concentration of HT-JF646 ligand (1 μM, 15 min at 37 °C) for FtsW localization. Data were from two independent biological replicates. (**A**) Representative micrographs of ectopically induced FtsW-Halo, FtsW(SSCC)-Halo (in the absence of DTT), FtsW(S382C)-Halo with MTSES, or FtsW(L198C)-Halo with MTSES. (**B**) Average expression levels of Halo-tagged FtsW and variants measured by mean JF646 fluorescence signal per pixel (illumination-corrected; log scale). Bars show geometric means ± geometric s.e.m.; small grey points are single-cell data points and large circles are replicate geometric means. Whole-cell signal was 101% (SSCC), 79% (S382C + MTSES) and 81% (L198C + MTSES) of WT (n = 159, 79, 113 and 55 cells for WT, SSCC, S382C + MTSES and L198C + MTSES); only S382C + MTSES differed significantly from WT (Welch's t-test on log-transformed values, $p = 5 \times 10^{-4}$). (**C**) Divisome localization efficiency of Halo-tagged FtsW and variants measured by the midcell band intensity relative to WT (n = 105, 52, 94 and 46 cells for WT, SSCC, S382C + MTSES and L198C + MTSES, respectively). Bars show geometric means ± geometric s.e.m. across single cells pooled from the two biological replicates; small grey points are single-cell data points and large circles are replicate geometric means. Geometric means were 1.05 (SSCC), 0.93 (S382C + MTSES) and 0.87 (L198C + MTSES) of WT; none differed significantly from WT (Welch's t-test on log-transformed values, $p \geq 0.38$).

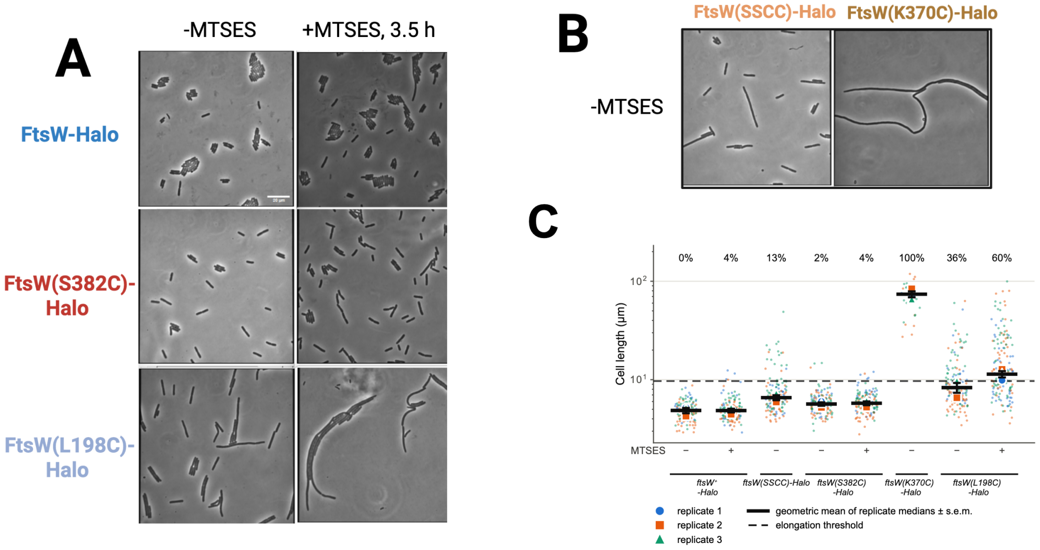


**Figure S9. Characterizations of dominant negative phenotypes of key FtsW active site mutants.**

**(A,B)** Phase-contrast images of *E. coli* FtsW merodiploid strains grown in EZRDM at 37°C co-expressing chromosomal WT *ftsW* and *ftsW-Halo* mutants from a pIQ plasmid induced with 100 μM IPTG. Note that 50 μM IPTG induction of *ftsW* in these plasmids is sufficient for complementation in an *ftsW* depletion background.

(**A**) Without MTSES treatment (left column), the *ftsW(L198C)-Halo* merodiploid strain (bottom) has a slightly elongated morphology relative to *ftsW-Halo* (top) and *ftsW(S382C)-Halo* (middle) cells. With 100 μM MTSES (right column), *ftsW(L198C)-Halo* cells showed a more dominant-negative, filamentous phenotype.

(**B**) In the absence of MTSES, cells expressing *ftsW(SSCC)-Halo* (left) and *ftsW(K370C)-Halo* (right) in a *wt ftsW* background exhibited dominant-negative, elongated, and filamentous phenotypes, respectively.

(**C**) Quantification of cell length for selected strains. SuperPlots summarize geometric means of cell lengths (μm) for *ftsW/ftsW-halo* merodiploid strains under growth conditions that lead to co-expression without MTSES (+IPTG, -MTSES), or with MTSES inhibition (+IPTG, +MTSES). Measurements were performed across three independent biological replicates. The dashed line marks the elongation threshold (twice the median length of the FtsW-Halo control), and the percentage above each group is the mean of the replicate percentages of cells longer than this threshold, as described for Fig. S7. *ftsW(K370C)-Halo* measurement was from 29 cells across three replicates, because its filaments were extremely long and few. Group summaries, pooled medians, per-replicate percentages and the bootstrap comparisons are in Supplementary Table 10.

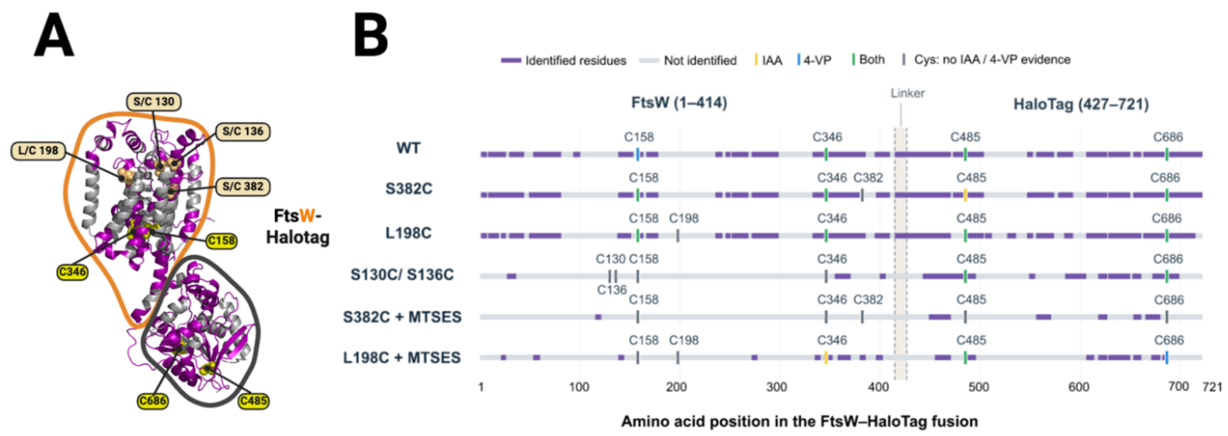


**Figure S10. Mass spectrometry of HaloTag-enriched FtsW-Halo variants.**

(**A**) AlphaFold model of the FtsW-Halo fusion, with FtsW outlined in orange and HaloTag in dark grey. Native cysteines (C158 and C346 in FtsW; C485 and C686 in HaloTag) are shown as yellow spheres; engineered positions (S130C, S136C, L198C and S382C) are shown as orange spheres.

(**B**) Sequence coverage and cysteine modifications in FtsW-Halo variants. Each horizontal line represents the FtsW-Halo fusion under the indicated condition. Purple segments indicate residues covered by identified peptides, whereas light-gray segments indicate regions without peptide coverage. Vertical ticks mark cysteine positions: yellow, iodoacetamide-derived carbamidomethylation (IAA); blue, 4-vinylpyridine modification (4-VP); green, both modifications observed in separate peptide identifications; dark gray, neither modification detected. Residue numbers refer to the full fusion sequence. Dashed lines delimit the linker (residues 415–426), separating FtsW (1–414) from HaloTag (427–721). Gray cysteine markers indicate insufficient peptide evidence to determine modification status and should not be interpreted as unmodified cysteines. Across WT and mutant samples with and without MTSES, peptides and the four native cysteines were identified at various levels with modifications (purple segments), demonstrating an oxidative periplasmic environment, but none of the engineered cysteines or their native residues (except S382 in wild-type FtsW-Halo and in FtsW(L198C)-Halo) were recovered, treated, or untreated. In FtsW(L198C)-Halo, 109 peptides were identified, covering 63.1% of the fusion, comparable to wild-type FtsW-Halo (107 peptides, 64.9%). FtsW(SSCC)-Halo recovered 28 peptides, covering 24.7% of the fusion. Peptide recovery across FtsW was lower in the three treatment samples, falling from 105–109 peptides in the untreated variants to 7–28 in the treated and double-cysteine samples, likely due to cell losses under inhibited conditions.

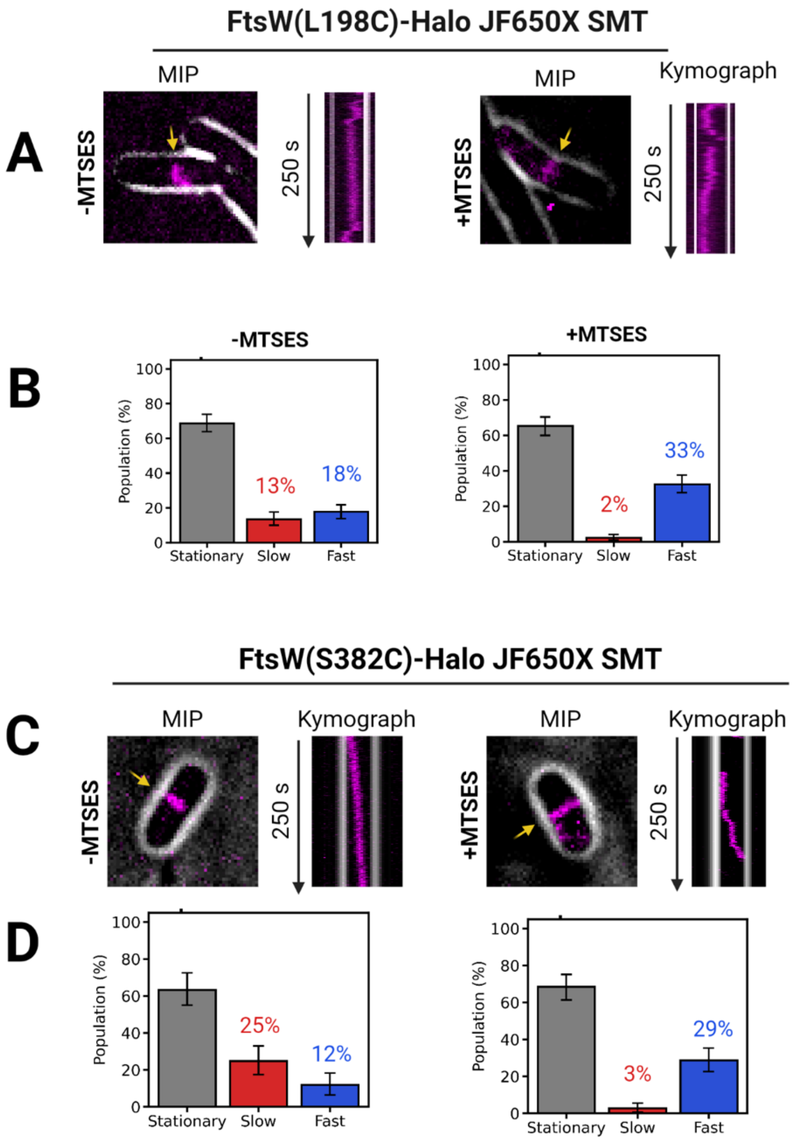


**Figure S11. Single-molecule tracking (SMT) reveals the inhibition of FtsW(S382C)-Halo and FtsW(L198C)**

(**A**) Representative FtsW(L198C) cells (black body outlined by light grey) overlaid with Maximum Intensity Projection (MIP) and corresponding single-molecule tracking trajectory (arrow) displayed as kymographs (magenta) over the course of 250s across the midcell. (**B**) Three classes of FtsW(L198C)-Halo single-molecule dynamics. Error bars represent 95% bootstrap confidence intervals from 313 (−MTSES) and 314 (+MTSES) trajectory segments pooled from two biological replicates. (**C**) Representative FtsW (S382C) cells (black body outlined by light grey) overlaid with Maximum Intensity Projection (MIP) and corresponding single-molecule tracking trajectory (arrow) displayed as kymographs (magenta) over the course of 250s across the midcell. (**D**) Three classes of FtsW(S382C) single-molecule dynamics. Error bars represent 95% bootstrap confidence intervals from 109 (-MTSES) and 181 (+MTSES) trajectory segments pooled from two biological replicates.

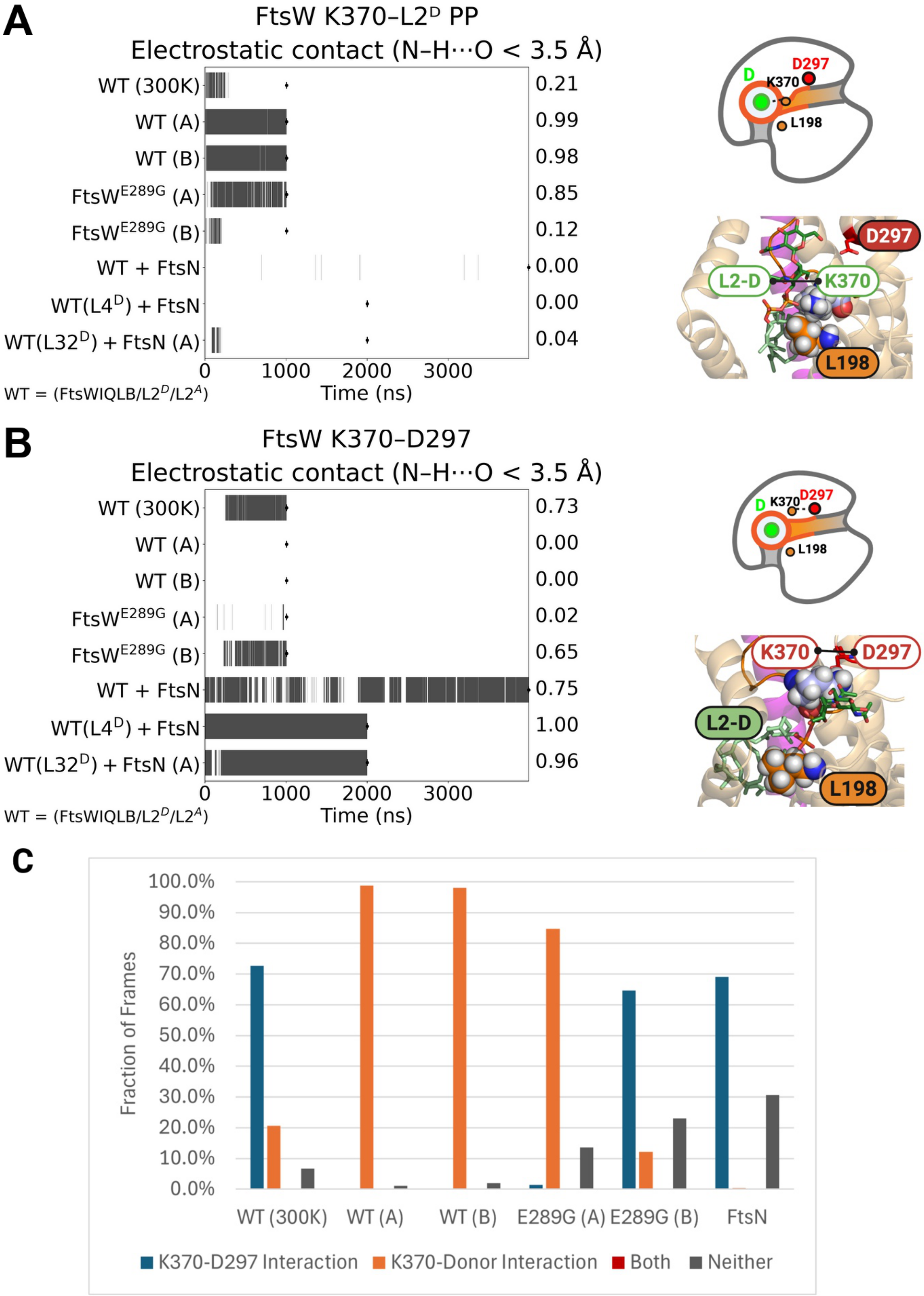
A
FtsW K370–L2ᴰ PP
Electrostatic contact (N–H···O < 3.5 Å)
WT (300K)
WT (A)
WT (B)
FtsWE289G (A)
FtsWE289G (B)
WT + FtsN
WT(L4D) + FtsN
WT(L32D) + FtsN (A)
0.21
0.99
0.98
0.85
0.12
0.00
0.00
0.04
0
1000
2000
3000
Time (ns)
WT = (FtsWIQLB/L2D/L2A)
D
D297
K370
L198
L2-D
B
FtsW K370–D297
Electrostatic contact (N–H···O < 3.5 Å)
WT (300K)
WT (A)
WT (B)
FtsWE289G (A)
FtsWE289G (B)
WT + FtsN
WT(L4D) + FtsN
WT(L32D) + FtsN (A)
0.73
0.00
0.00
0.02
0.65
0.75
1.00
0.96
0
1000
2000
3000
Time (ns)
WT = (FtsWIQLB/L2D/L2A)
C
Fraction of Frames
100.0%
90.0%
80.0%
70.0%
60.0%
50.0%
40.0%
30.0%
20.0%
10.0%
0.0%
WT (300K)
WT (A)
WT (B)
E289G (A)
E289G (B)
FtsN
K370-D297 Interaction
K370-Donor Interaction
Both
Neither

**Figure S12. K370 alternates between coordinating the donor pyrophosphate and contacting catalytic D297.**

**(A)** Binary contact traces for the interaction between FtsW K370 and the donor lipid II pyrophosphate (donorPP), scored using an N–H⋯O distance criterion (< 3.5 Å). Each row corresponds to one trajectory (listed at left); tick marks denote frames satisfying the contact criterion, and values at right report the overall contact fraction. Right, representative snapshot showing the K370–donorPP geometry ($L2^{D}$) in the TM6–TM7 outer-gate region relative to L198 and catalytic D297.

**(B)** Binary contact traces for the K370–D297 N–H⋯O interaction (distance < 3.5 Å) across the same set of trajectories, displayed as in (A). Right, representative snapshot illustrating the K370–D297 contact geometry.

**(C)** Summary of contact-state occupancy for each condition, classifying frames as K370–D297 only, K370–donorPP only, both, or neither using the criteria in **(A–B)**.

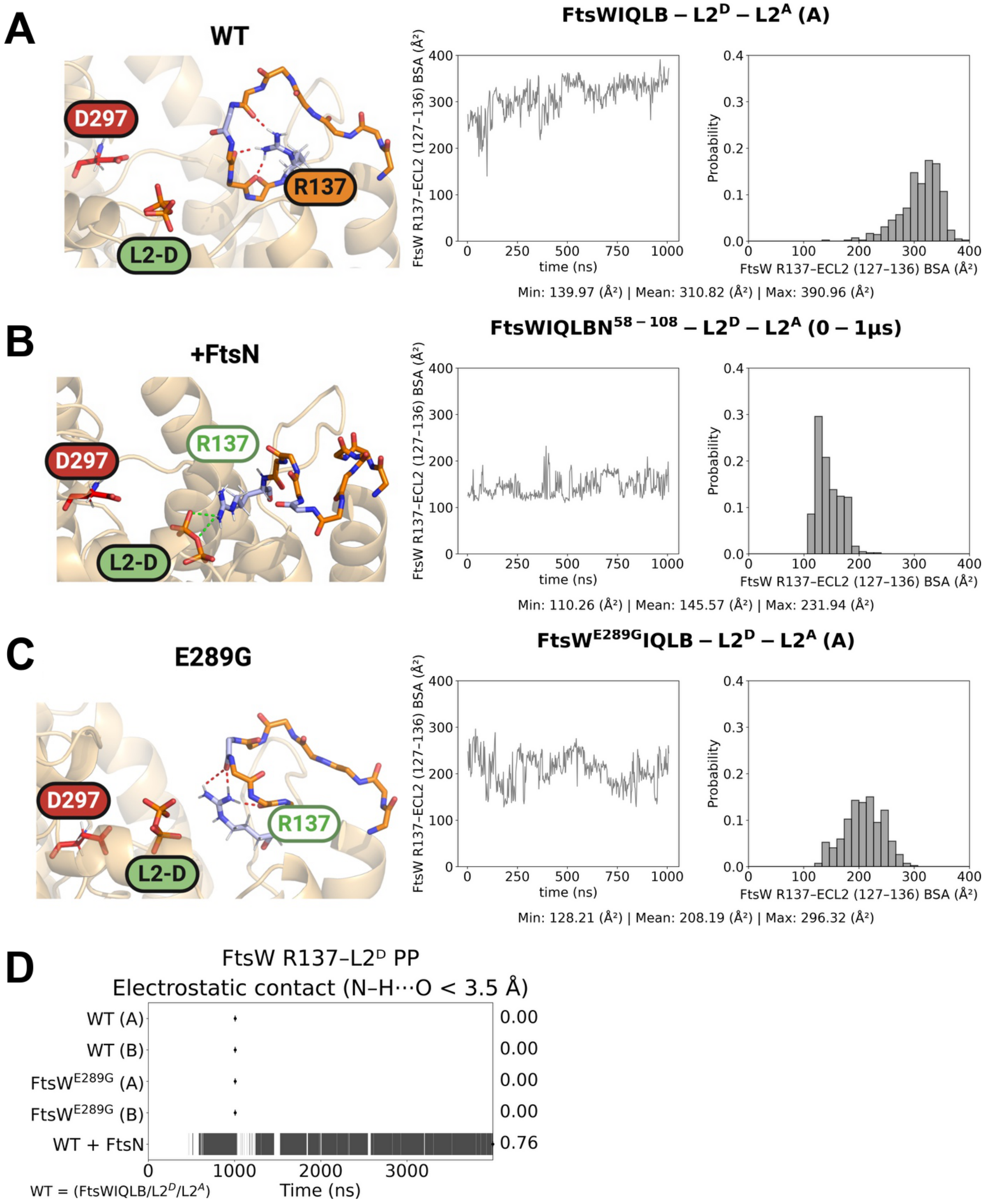


**Figure S13. FtsN remodels the ECL2 conformation and promotes R137 coordination of the donor pyrophosphate.**

**(A–C)** Representative snapshots (left) showing the position of FtsW R137 relative to the donor lipid II substrate ($L2^D$) and catalytic D297 in WT (A), +FtsN (B), and FtsW(E289G) (**C**). Plots at right report the buried surface area

(BSA) between R137 and ECL2 (FtsW residues 127–136) over time (left) and as a distribution (right); minimum, mean, and maximum BSA values are listed below each histogram.

**(D)** Binary contact traces for the R137–donor pyrophosphate interaction, scored using an N–H⋯O distance criterion (< 3.5 Å). Each row corresponds to one trajectory (listed at left); tick marks denote frames satisfying the contact criterion, and values at right report the overall contact fraction.

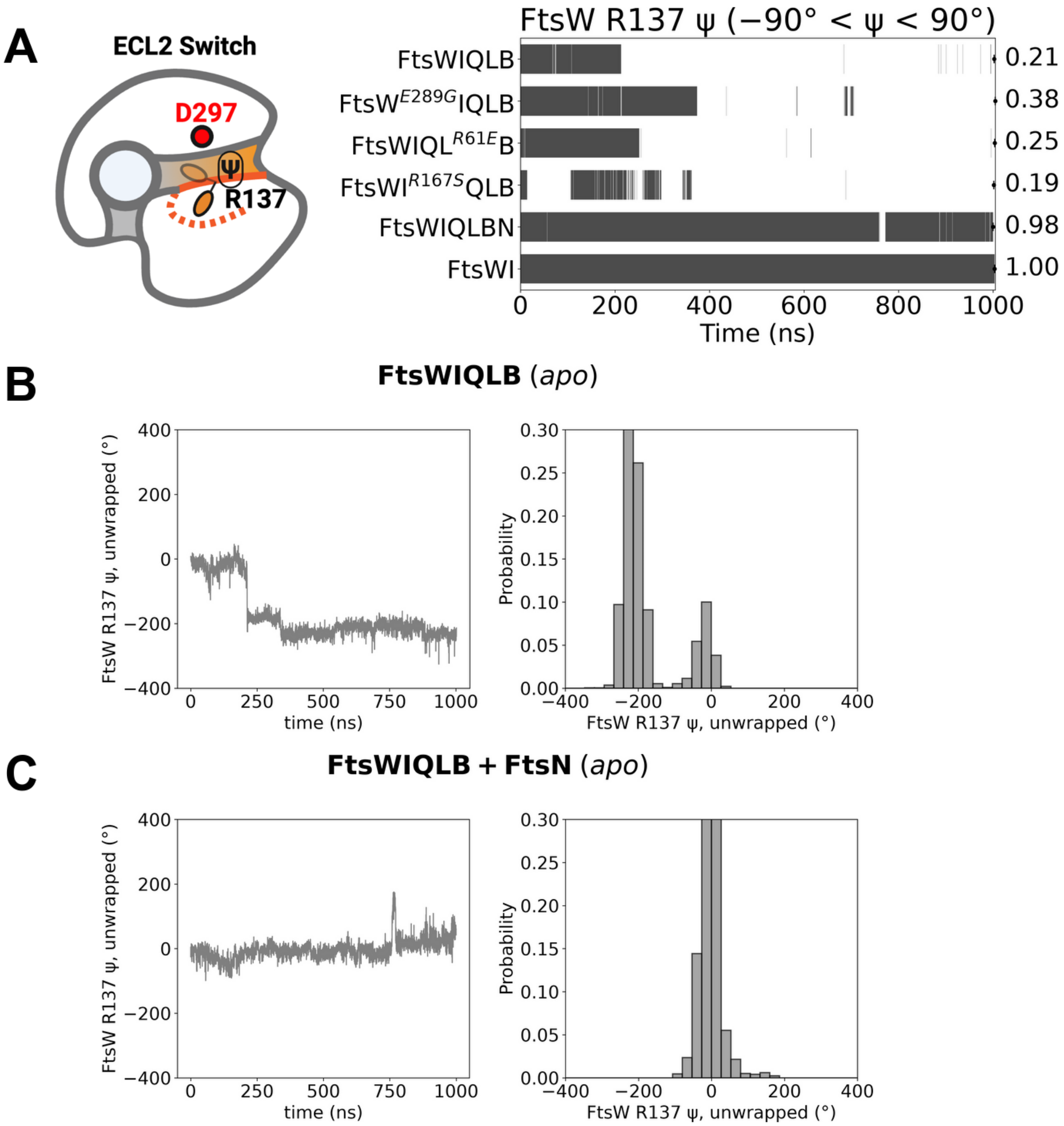


**Figure S14. ECL2 dynamics and R137 ψ-angle switching across apo divisome assemblies.**

**(A)** Schematic illustration of the ECL2 switch, highlighting the backbone ψ dihedral of R137 and position relative to the catalytic residue D297. Binary occupancy plots (right) highlight the fraction of each trajectory classified as active (−90° < ψ < 90°) for each apo trajectory analyzed. The bar plot reports the active-state occupancy for each apo complex analyzed: FtsWIQLB (0.21), FtsW(E289G)IQLB (0.38), FtsWIQLB carrying FtsL(R61E) (0.25), FtsWI(R167S)QLB (0.19), FtsWIQLB + FtsN (0.98), and FtsWI in the absence of the FtsQLB subcomplex (1.00).

**(B)** FtsWIQLB ECL2 dynamics. Left, time series of the unwrapped R137 ψ angle over the 1-μs trajectory. Right, corresponding probability distributions.

**(C)** FtsWIQLB + FtsN$^E$ ECL2 dynamics. Left, time series of the unwrapped R137 ψ angle. Right, corresponding probability distribution.

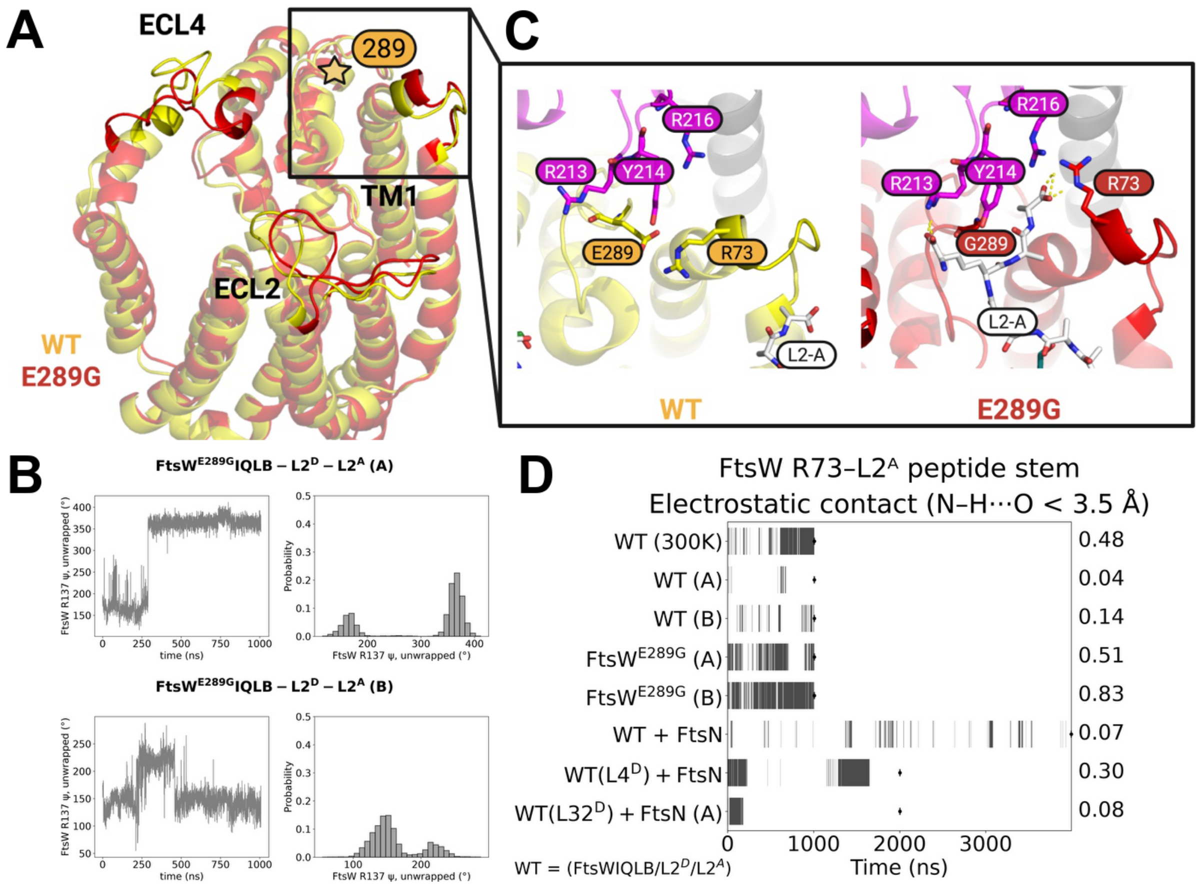


**Figure S15. E289G remodels the FtsW backbone and increases R73 contacts with the acceptor peptide stem.**

**(A)** Structural overlay of WT and FtsW$^{E289G}$ highlighting differences in ECL2, ECL4, and TM1 (WT, yellow; E289G, red).

**(B)** Time traces (Left) and histograms (Right) tracking the unwrapped Psi Angle of FtsW R137 in E289G-A (Top) and E289G-B (Bottom)

**(C)** Expanded view of the acceptor-site region comparing WT (E289) and E289G (G289) and showing the positioning of R73 relative to nearby residues and the acceptor lipid-II (L2-A).

**(D)** Binary contact occupancy over time for FtsW R73 contact with acceptor peptide stem acids (X < 3.5 Å) across trajectories listed at left. Black segments indicate snapshots meeting the contact criterion; values at right report the time-weighted fraction of snapshots in contact.

contact.

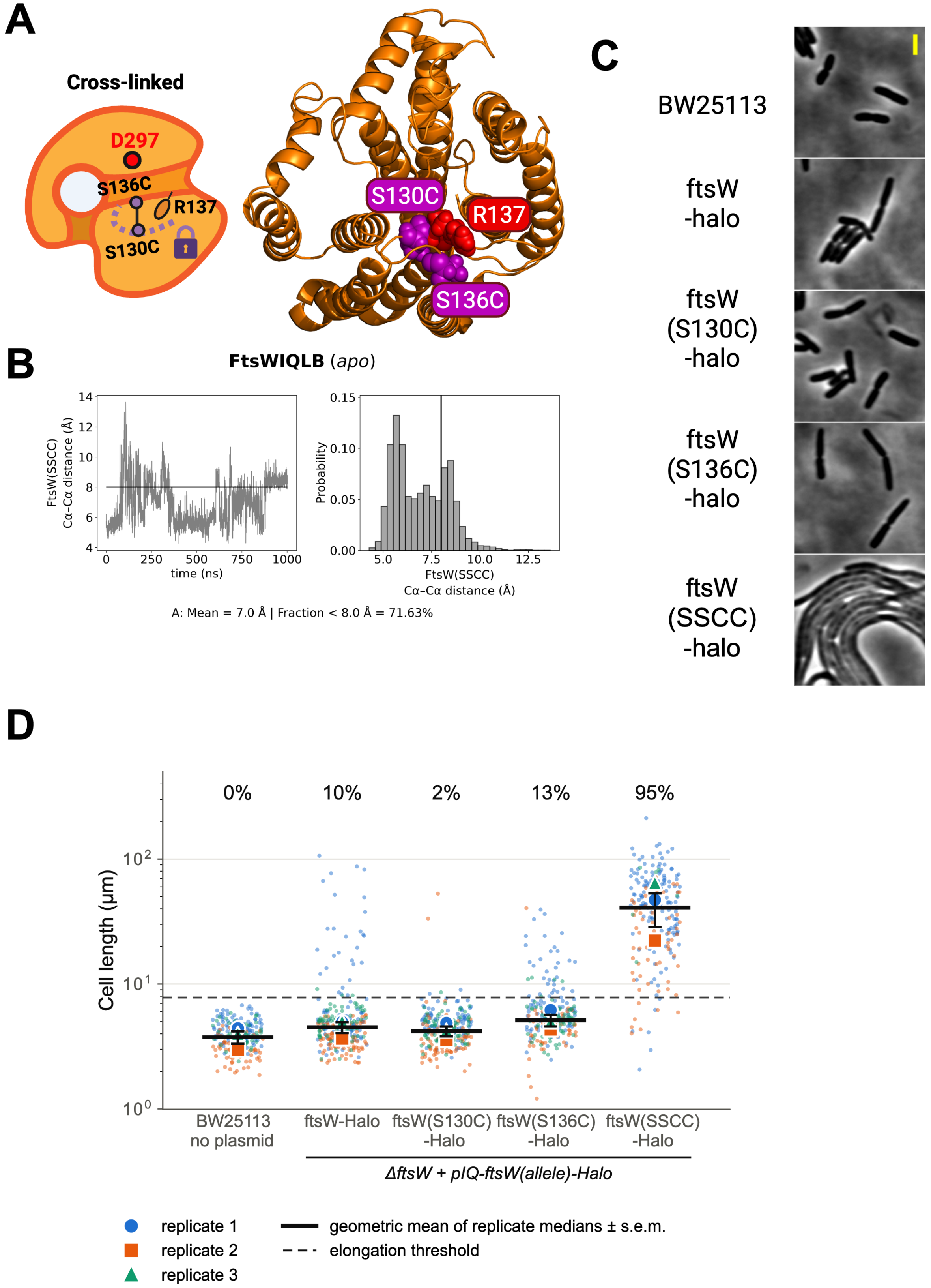
A
Cross-linked
D297
S136C
R137
S130C
S130C
R137
S136C
B
FtsWIQLB (apo)
FtsW(SSCC) Cα-Cα distance (Å)
time (ns)
Probability
FtsW(SSCC) Cα-Cα distance (Å)
A: Mean = 7.0 Å | Fraction < 8.0 Å = 71.63%
C
BW25113
ftsW -halo
ftsW (S130C) -halo
ftsW (S136C) -halo
ftsW (SSCC) -halo
D
0%
10%
2%
13%
95%
Cell length (µm)
BW25113 no plasmid
ftsW-Halo
ftsW(S130C) -Halo
ftsW(S136C) -Halo
ftsW(SSCC) -Halo
ΔftsW + pIQ-ftsW(allele)-Halo
replicate 1
replicate 2
replicate 3
geometric mean of replicate medians ± s.e.m.
elongation threshold

**Figure S16. Disulfide crosslinking of ECL2 restricts its dynamics and inhibits cell division.**

(**A**) Schematic and structural view of the engineered ECL2 disulfide crosslink. ECL2 residues S130 and S136 (purple) were mutated to cysteines to permit disulfide formation in the oxidative periplasm, thereby restricting ECL2 mobility without mutating R137 (red). The position of R137 and the catalytic residue D297 are indicated. (**B**) In WT apo simulations, S130 and S136 are frequently positioned within disulfide-forming distance. Left, time series of the FtsW S130–S136 Cα–Cα distance; right, corresponding distribution (vertical line at 8 Å). (**C**) Phase-contrast microscopy of *E. coli ftsW* depletion strains complemented with the indicated *ftsW* alleles in the oxidative periplasmic environment (- DTT). Single mutants FtsW(S130C)-halo and FtsW(S136C)-halo maintain WT-like morphology, whereas the double mutant FtsW(SSCC)-Halo exhibits pronounced filamentation under non-reducing conditions, consistent with inhibition of division upon ECL2 crosslinking. (**D**) Cell length measurements for strains in (**C**), grown in EZRDM at 37 °C with 50 μM IPTG and no DTT. BW25113 carries no plasmid; the remaining strains are *ftsW*-depletion strains carrying the indicated pIQ-*ftsW(allele)-halo*. Neither single mutant filamented (1.9% and 13.2% of cells elongated for S130C and S136C, versus 9.9% for ftsW-halo), whereas 95.0% of double-mutant cells were elongated (41 ± 12 μm; n = 3 biological replicates, 217 cells). The dashed line marks the elongation threshold, twice the median length of the BW25113 control, and the percentage above each group is the mean of the replicate percentages of cells longer than this threshold (plot conventions as in Fig. S7). Group summaries, pooled medians, per-replicate percentages and the bootstrap comparisons are in Supplementary Table 7.

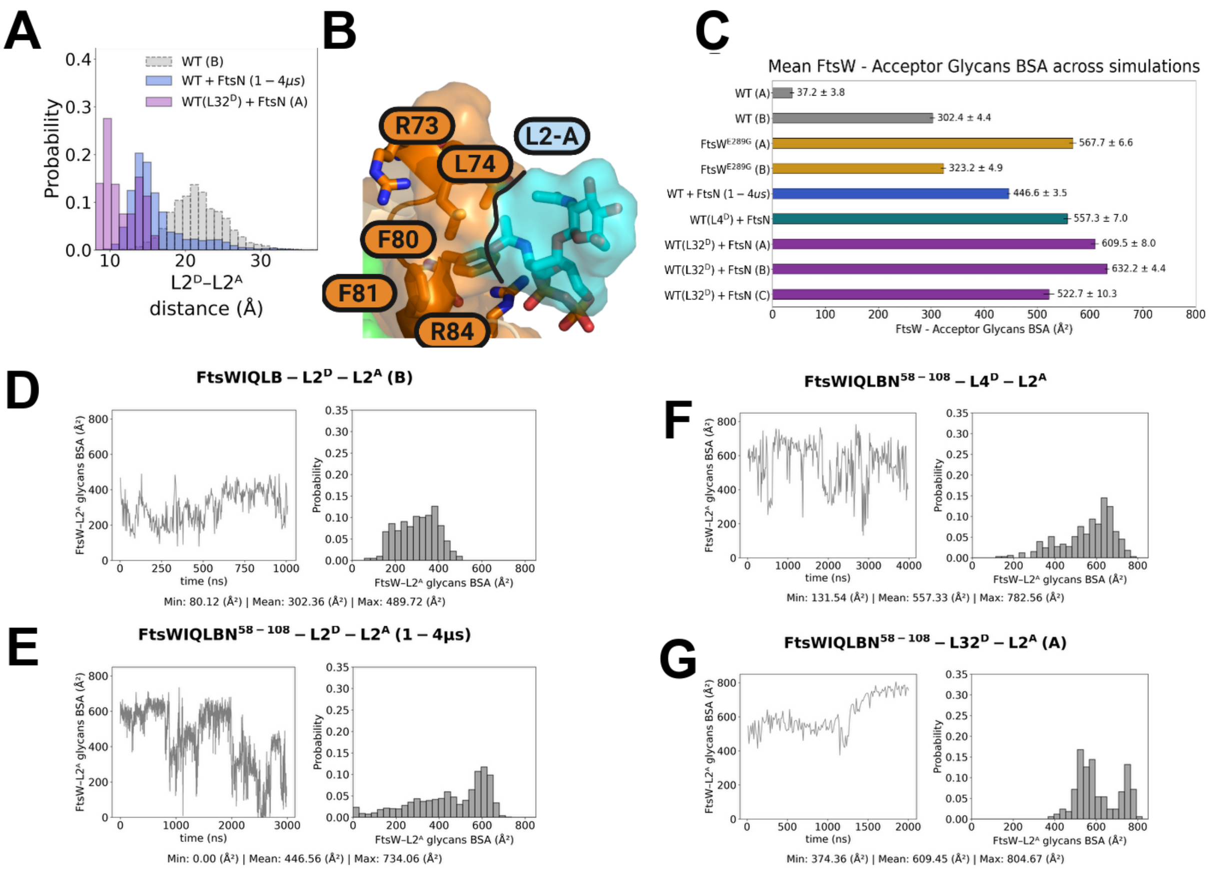


**Figure S17. Quantification of acceptor-site engagement by buried surface area (BSA).**

**(A)** Probability distributions of donor–acceptor separation (donor anomeric carbon C1 to acceptor O4) for the indicated simulations: WT FtsWIQLB–L2$^{D}$–L2$^{A}$ (replicate B; gray), activated FtsWIQLBN$^{58\text{-}108}$–L2$^{D}$–L2$^{A}$ (1–4 μs; blue), and processive-like FtsWIQLBN$^{58\text{-}108}$–L32$^{D}$–L2$^{A}$ (replicate A; purple).

**(B)** Structural illustration of the acceptor-site interface used for BSA analysis. The acceptor lipid II headgroup (L2-A; cyan) packs against FtsW (orange; representative residues indicated).

**(C)** Mean FtsW–acceptor glycan BSA averaged over each simulation condition (mean ± s.e.m. across frames), showing weak and intermittent acceptor engagement in WT trajectories (gray; WT-A and WT-B), enhanced engagement in the superfission FtsW(E289G) mutant (gold), and strong stabilization in activated complexes containing FtsN58–108 (blue) and in processive-like systems with extended donor chains (teal/purple).

For the simulation with the E289G mutation, increased BSA in replicate B is promoted by the R73-dependent peptide-stem dynamics described previously (**Fig. S15D**), and in replicate A it coincides with partial strengthening of these same clamp-associated contacts (**Fig. S18A, B**).

**(D–G)** Representative time traces (left) and corresponding probability distributions (right) of FtsW–acceptor glycan BSA for the indicated systems: **(D)** WT FtsWIQLB–L2D–L2A (replicate B), **(E)** FtsWIQLBN$^{58–108}$–L2$^{D}$–L2$^{A}$, **(F)** FtsWIQLBN$^{58–108}$–L4$^{D}$–L2$^{A}$, and **(G)** FtsWIQLBN$^{58–108}$–L32$^{D}$–L2$^{A}$ (replicate A). In each panel, sustained high-

BSA episodes reflect deep acceptor packing in the catalytic groove, whereas near-zero BSA corresponds to acceptor disengagement. BSA values are reported in $Å^2$.

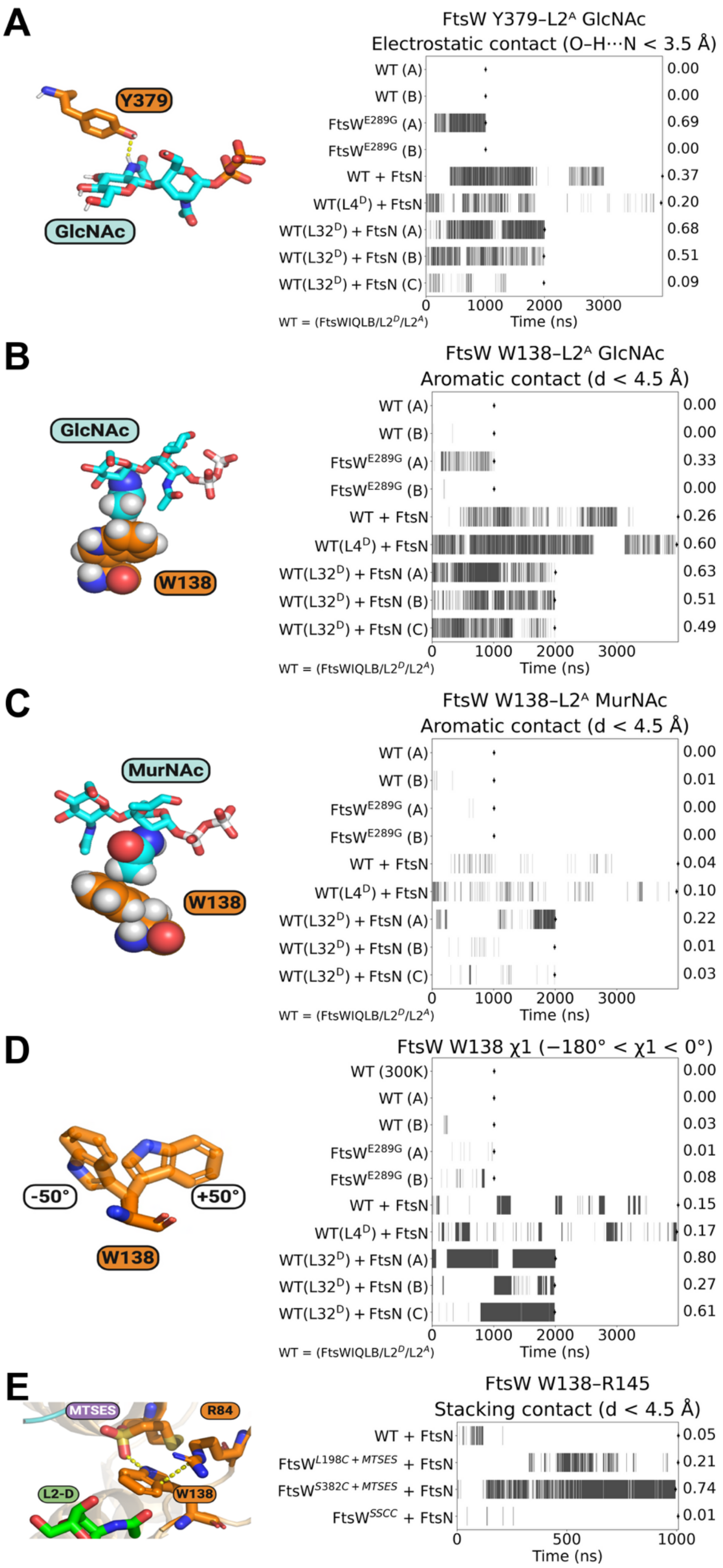
A
Y379
GlcNAc
FtsW Y379–L2ᴬ GlcNAc
Electrostatic contact (O–H···N < 3.5 Å)
WT (A) 0.00
WT (B) 0.00
FtsW$^{E289G}$ (A) 0.69
FtsW$^{E289G}$ (B) 0.00
WT + FtsN 0.37
WT(L4$^{D}$) + FtsN 0.20
WT(L32$^{D}$) + FtsN (A) 0.68
WT(L32$^{D}$) + FtsN (B) 0.51
WT(L32$^{D}$) + FtsN (C) 0.09
0 1000 2000 3000
Time (ns)
WT = (FtsWIQLB/L2$^{D}$/L2$^{A}$)
B
GlcNAc
W138
FtsW W138–L2ᴬ GlcNAc
Aromatic contact (d < 4.5 Å)
WT (A) 0.00
WT (B) 0.00
FtsW$^{E289G}$ (A) 0.33
FtsW$^{E289G}$ (B) 0.00
WT + FtsN 0.26
WT(L4$^{D}$) + FtsN 0.60
WT(L32$^{D}$) + FtsN (A) 0.63
WT(L32$^{D}$) + FtsN (B) 0.51
WT(L32$^{D}$) + FtsN (C) 0.49
0 1000 2000 3000
Time (ns)
WT = (FtsWIQLB/L2$^{D}$/L2$^{A}$)
C
MurNAc
W138
FtsW W138–L2ᴬ MurNAc
Aromatic contact (d < 4.5 Å)
WT (A) 0.00
WT (B) 0.01
FtsW$^{E289G}$ (A) 0.00
FtsW$^{E289G}$ (B) 0.00
WT + FtsN 0.04
WT(L4$^{D}$) + FtsN 0.10
WT(L32$^{D}$) + FtsN (A) 0.22
WT(L32$^{D}$) + FtsN (B) 0.01
WT(L32$^{D}$) + FtsN (C) 0.03
0 1000 2000 3000
Time (ns)
WT = (FtsWIQLB/L2$^{D}$/L2$^{A}$)
D
-50°
+50°
W138
FtsW W138 χ1 (−180° < χ1 < 0°)
WT (300K) 0.00
WT (A) 0.00
WT (B) 0.03
FtsW$^{E289G}$ (A) 0.01
FtsW$^{E289G}$ (B) 0.08
WT + FtsN 0.15
WT(L4$^{D}$) + FtsN 0.17
WT(L32$^{D}$) + FtsN (A) 0.80
WT(L32$^{D}$) + FtsN (B) 0.27
WT(L32$^{D}$) + FtsN (C) 0.61
0 1000 2000 3000
Time (ns)
WT = (FtsWIQLB/L2$^{D}$/L2$^{A}$)
E
MTSES
R84
L2-D
W138
FtsW W138–R145
Stacking contact (d < 4.5 Å)
WT + FtsN 0.05
FtsW$^{L198C+MTSES}$ + FtsN 0.21
FtsW$^{S382C+MTSES}$ + FtsN 0.74
FtsW$^{SSCC}$ + FtsN 0.01
0 500 1000
Time (ns)

**Figure S18. FtsN and long donor chains reinforce acceptor-site packing through clamp-associated contacts and a W138 rotameric switch**.

**(A)** Structural illustration and binarized trajectory of a hydrogen bond between FtsW Y379 and the acceptor GlcNAc N-acetyl group, quantified as a binarized O–H⋯N distance (with a threshold of $X < 3.5$ Å).

**(B)** Structural illustration of and binarized trajectory of methyl–aromatic packing between the W138 indole ring and the acceptor GlcNAc N-acetyl methyl group, quantified as proximity of the acetyl methyl carbon to the W138 ring centroid (with a threshold of $X < 4.5$ Å).

**(C)** Structural illustration of and binarized trajectory of W138 methyl–aromatic contact metric for the acceptor MurNAc N-acetyl group (with a threshold of $X < 4.5$ Å).

**(D)** Structural illustration of and binarized trajectory of the W138 $\chi1$ dihedral angle ($-180° < \chi1 < 0°$), reporting its conformational bias between outward- and inward-facing rotamers.

**(E)** Structural illustration and binarized trajectory of the W138–R145 stacking contact (distance < 4.5 Å) for the FtsN-bound $L2^D$–$L2^A$ reference simulation and the indicated engineered-variant trajectories; occupancies at right report the fraction of frames satisfying the contact criterion.

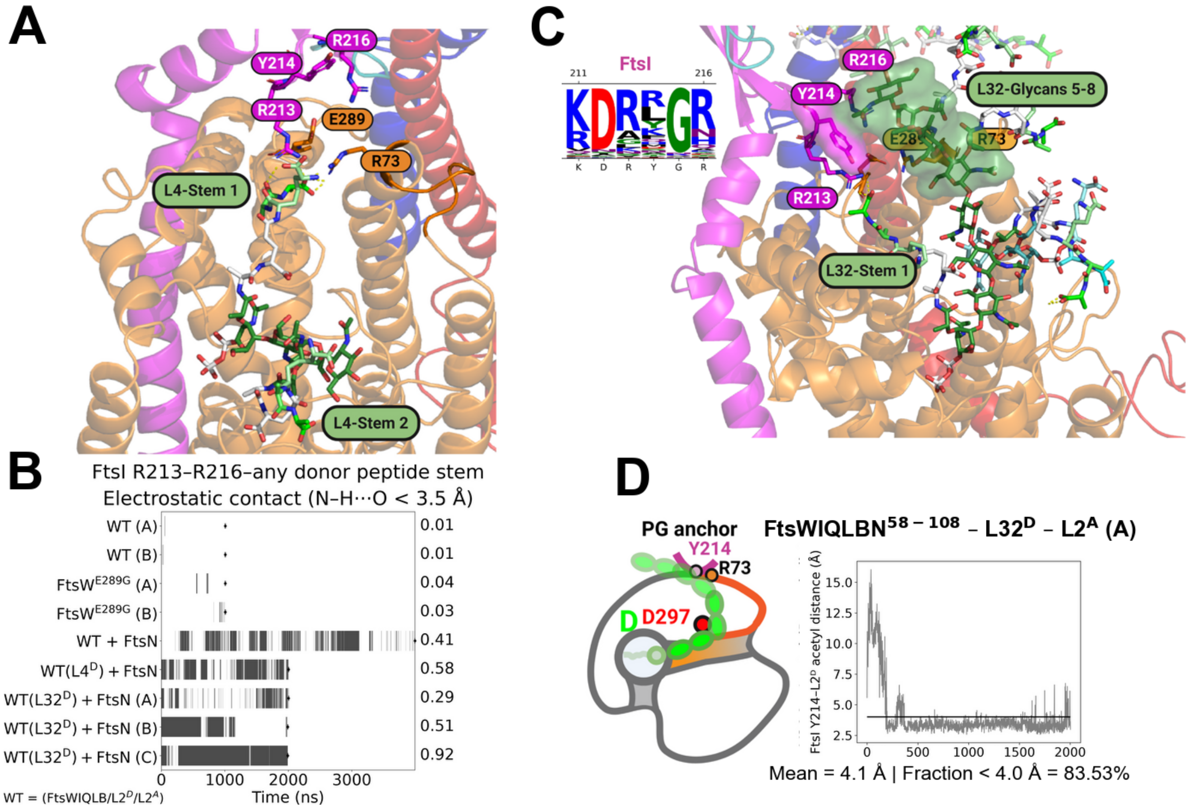


**Figure S19. Donor peptide-stem tethering to the FtsI anchor loop reveals a chain-length dependent Y214–acetyl contact.**

**(A)** Representative snapshot from the FtsN-bound L4 donor simulation showing donor peptide stems positioned near the FtsI anchor loop; selected anchor-loop residues (R213, Y214, R216) and nearby FtsW residues are labeled for reference.

**(B)** Binary contact occupancy over time for FtsI anchor-loop–donor-stem contacts (X < 3.5 Å; N–O), scored as snapshots in which either anchor-loop arginine (R213 or R216) contacted any donor stem acid. Black segments indicate snapshots meeting the criterion; values at right report the time-weighted fraction of snapshots in contact.

**(C)** Representative snapshot from the FtsN-bound L32 donor simulation highlighting the extended polymer (glycans 5–8) sampling the anchor-loop region. A logo plot for the FtsI sequence from amino acid 211 to 216, built from 3,859 FtsI sequences from distinct organisms, is included with a y-axis of sequence counts.

**(D)** Schematic and time series of donor acetyl proximity to Y214 in the L32 trajectory, computed as the minimum distance between any donor acetyl methyl carbon and any Y214 ring atom; the fraction below 4.0 Å indicates sustained acetyl–Y214 contact after the initial relaxation period.

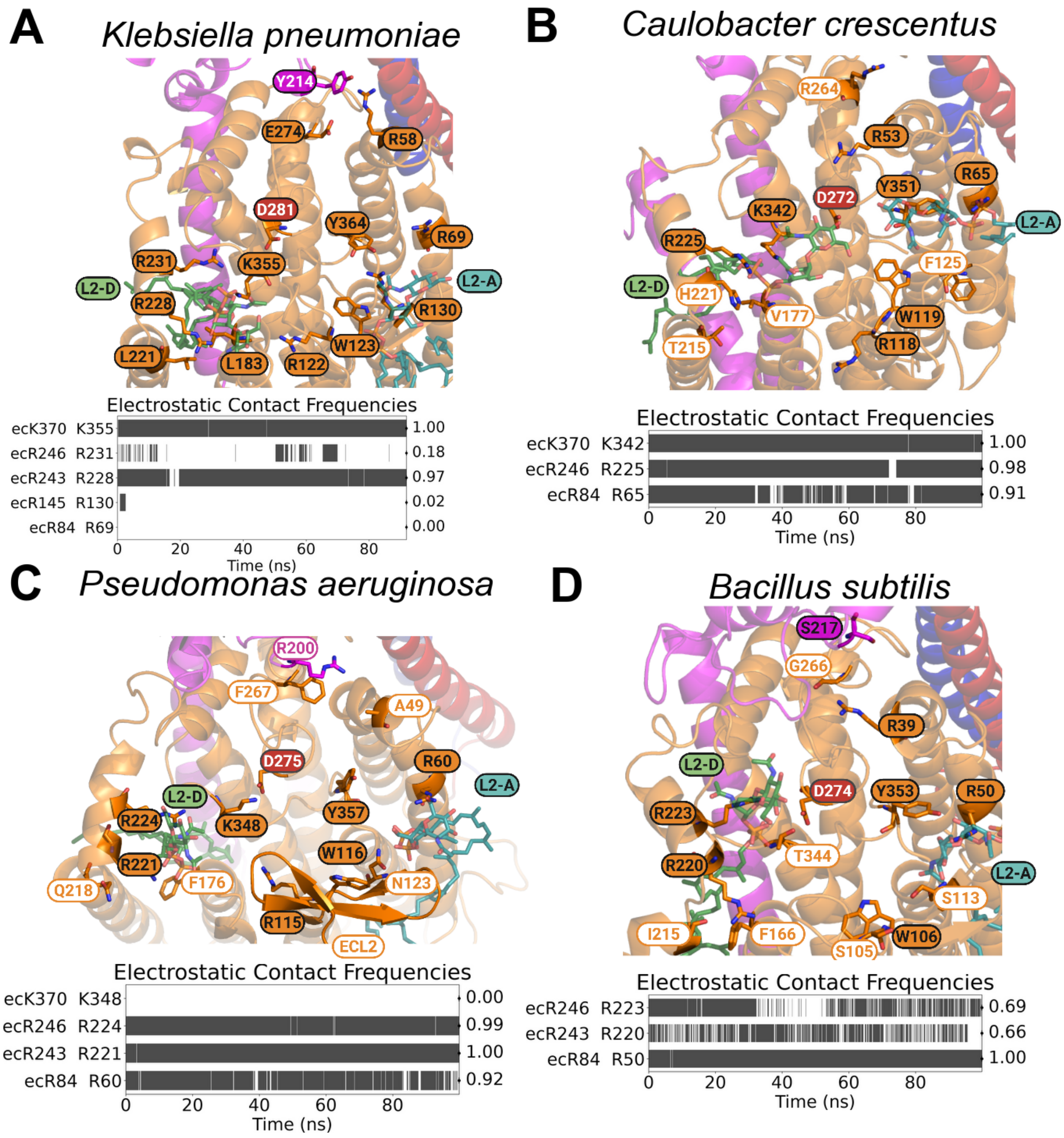


**Figure S20. Conserved lipid II coordination and electrostatic contact frequencies across FtsW homolog models.**

**(A–D)** Docked lipid II (L2) models for FtsW homologs from *Klebsiella pneumoniae* (**A**), *Caulobacter crescentus* (**B**), *Pseudomonas aeruginosa* (**C**), and *Bacillus subtilis* (**D**). Donor and acceptor substrates are labeled and colored ($L2^D$, green; $L2^A$, cyan). Residues that are identical to the E Coli sequence are colored orange with black text, and positions that encode a different residue are colored white with orange text. For each homolog, the electrostatic contact timeline below the structure reports frames from a short, unbiased MD trajectory in which the indicated residue–phosphate interaction satisfies the contact criterion (distance cutoff as indicated in the panel). Values at right report the overall fraction of frames meeting the criterion.

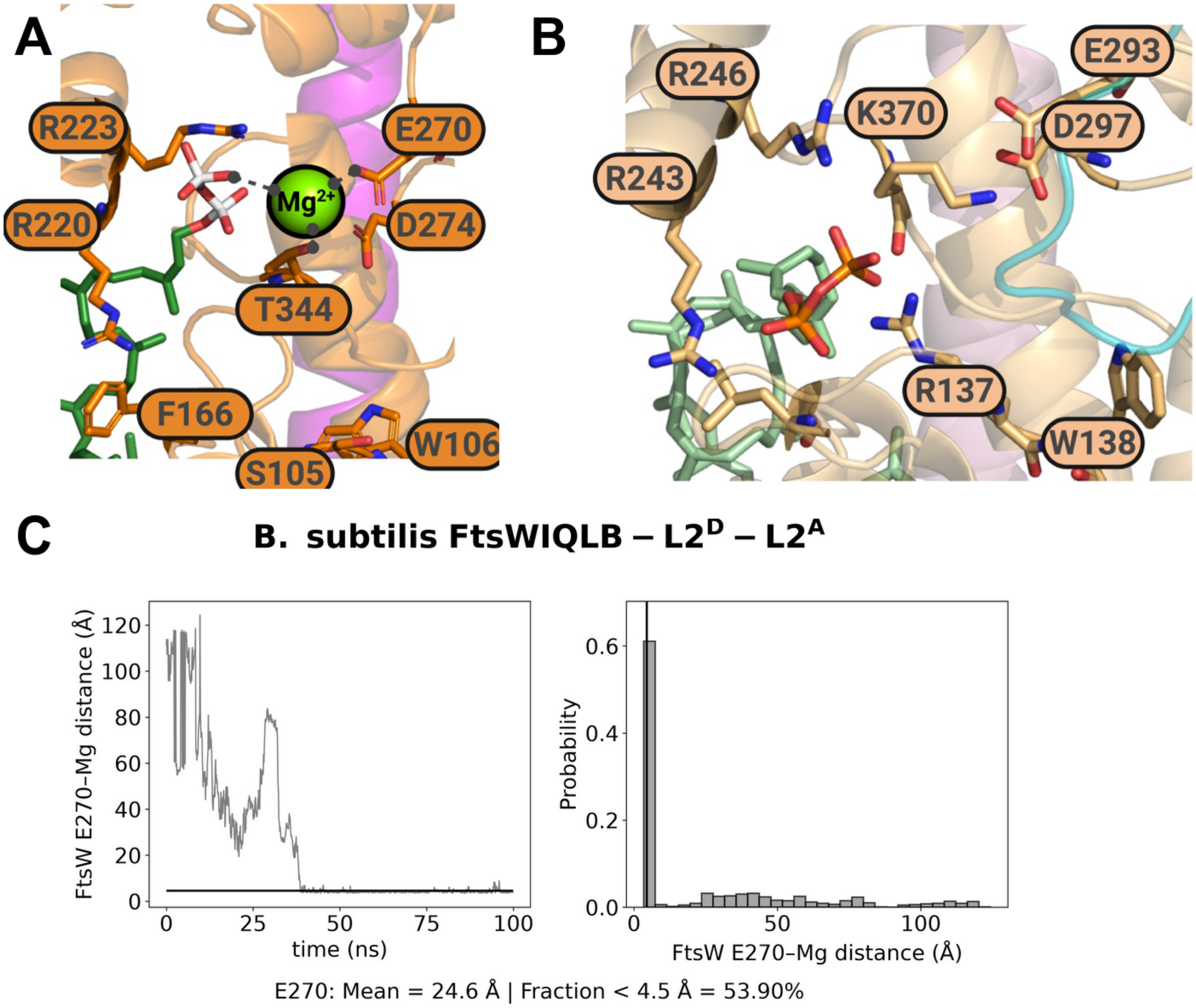


**Figure S21. Distinct Mg2+ coordination behavior in the donor site of *Bacillus subtilis* FtsW.**

**(A)** Donor-site view of the *B. subtilis* FtsW model with docked donor lipid II and Mg2+ highlighting proximity to the donor pyrophosphate and E270.

**(B)** Matched donor-site view from *E. coli* FtsW highlighting the Gram-negative K370 and R137 architecture.

**(C)** Time trace and distribution of the E270–Mg2+ distance in the *B. subtilis* trajectory, showing rapid localization of Mg2+ near E270 and sustained proximity over the remainder of the simulation.

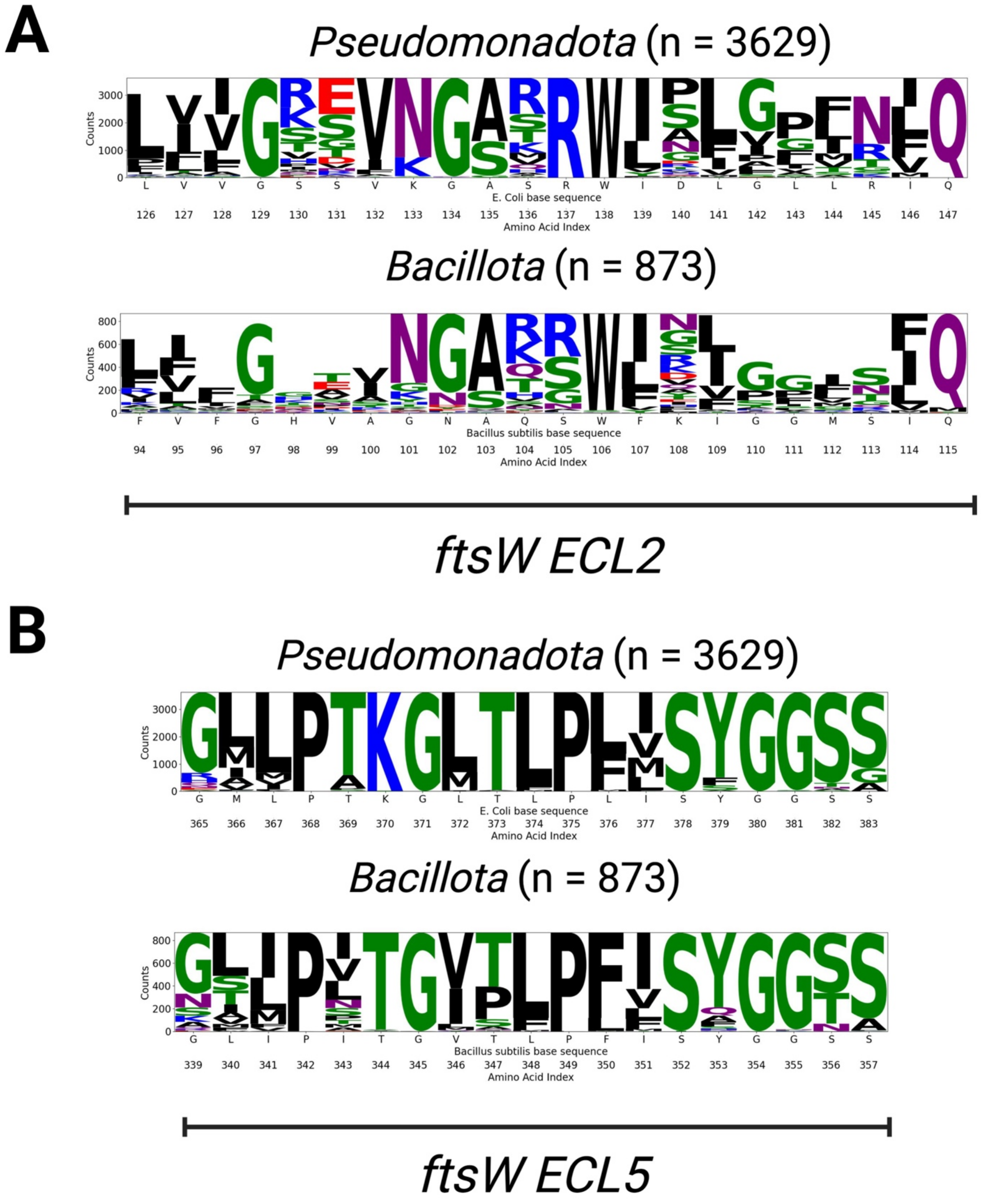


**Figure S22. Phylum-level conservation of ECL2 and ECL5 sequence features in FtsW.** Sequence logo representations of multiple sequence alignments of FtsW orthologs from the Pseudomonadota (n = 3,629) and Bacillota (n = 873) phyla.

**(A)** ECL2 region. Logos are shown for the segment corresponding to *E. coli* FtsW ECL2, with position indices shown relative to the phylum-specific reference sequence indicated beneath each logo (*E. coli* for Pseudomonadota; *Bacillus subtilis* for Bacillota).

**(B)** ECL5 region. Logos are shown for the segment corresponding to *E. coli* FtsW ECL5, displayed as in (A).

For all panels, letter height reflects residue conservation (counts), with the most conserved residues shown in larger font. Colors indicate residue class: basic (blue), acidic (red), polar uncharged (green), and hydrophobic (black).

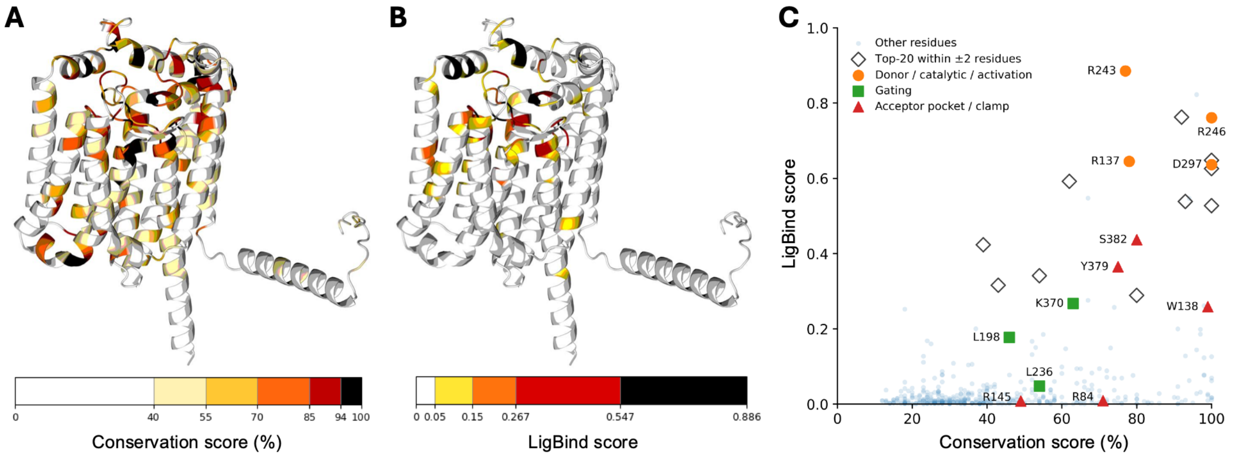


**Figure S23. Sequence conservation and predicted ligand-binding propensity across FtsW.**

**(A)** Sequence conservation mapped onto an ESMFold2 model of the 414-residue FtsW consensus sequence generated from the 6,859-sequence alignment used for Fig. S3. E. coli FtsW is used as the positional scaffold. Conservation is the frequency of the modal amino acid among sequences containing an amino acid at that position, with alignment gaps excluded.

**(B)** AiPP-LigBind scores for the consensus FtsW sequence mapped onto the same structural model.

**(C)** Correlation between the residue-level conservation and ligand binding propensity (AiPP-LigBind score) across the 414 consensus positions. Predefined mechanistic positions are grouped as donor/catalytic/activation residues (R137, R243, R246, and D297), gating residues (L198, L236, and K370), and acceptor-pocket/clamp residues (R84, W138, R145, Y379, and S382). Open diamonds indicate the Top-20 residues ranked by AiPP-LigBind that fall within ±2 positions of a predefined mechanistic site but are not themselves one of those sites. Residue numbering follows the E. coli FtsW positional scaffold.

**Supplementary Movie legends**

**Supplementary Movie 1.** Lipid entry through the FtsW TM6–TM7 gate. In a trajectory of the apo FtsWIQLB system, FtsW TM7 opens outward and a POPE lipid from the outer leaflet slithers through the opening between TM6 and TM7 toward the putative donor site. At ~333 ns one POPE tail partially enters the donor cavity, and FtsW R243 and R246 both contact the POPE phosphate head group.

**Supplementary Movie 2.** Acceptor-site dynamics in the FtsWIQLB-L2$^D$-L2$^A$ simulation. The GlcNAc moiety of the acceptor disaccharide flips down toward the catalytic groove, bringing the donor and acceptor reaction atoms closer together and closer to D297 (Fig. 1F).

**Supplementary Movie 3.** Metadynamics simulation of donor-site entry. L2 moves from the outer (D') to the inner (D) position as the C55 tail threads between TM6 and TM7, forming new electrostatic contacts with FtsW K370 and R246. The lipid tail crosses the gate by ~60 ns, followed by partial gate closure behind the tail at ~100 ns and complete cavity closure by ~150 ns (Fig. S6).

**Supplementary Movie 4.** Morph between the final snapshots of simulation 14 (WT-A) and simulation 17 (+FtsN) (Table 2), showing the rearrangements in the interactions between FtsW and the two bound L2 molecules in the FtsN-bound complex. The movie is an interpolation between two end-point structures and is not a continuous trajectory.

**Supplementary Movie 5.** Acceptor-site view of the superfission variant FtsW(E289G)IQLB-L2$^D$-L2$^A$ (1 μs, 310 K, 10 mM $Mg^{2+}$), first replicate, showing the donor L2 disaccharide flipping into the catalytic groove, shifted outer and inner gates, and increased flexibility of FtsW around E289 in ECL4.

**Supplementary Movie 6.** Second FtsW(E289G)IQLB-L2$^D$-L2$^A$ replicate, started from the same coordinates with randomized initial velocities, reproducing the shifted outer- and inner-gate geometries and the donor L2 disaccharide flipping into the catalytic groove.

**Supplementary Movie 7.** Acceptor-site binding in a WT FtsWIQLB-L2$^D$-L2$^A$ simulation (replicate WT-A). Acceptor binding is intermittent and is not stably maintained in the catalytic groove.

**Supplementary Movie 8.** Acceptor-site binding in the FtsWIQLB-FtsN58–108-L2$^D$-L2$^A$ simulation. In contrast to WT-A (Supplementary Movie 7), FtsN58–108 stabilizes acceptor binding in the catalytic groove.

**Supplementary Movie 9.** Acceptor site with a Lipid IV (L4) donor chain in the FtsWIQLB-FtsN58–108 system, including the extended 4 μs simulation. The short PG chain is largely unconstrained and explores the central cavity, making only transient contacts with ECL2, and acceptor binding remains highly dynamic.

**Supplementary Movie 10.** Acceptor site with a Lipid XXXII (L32) donor chain in the FtsWIQLB-FtsN58–108 system. The longer chain, docked like L4 at the start, forms a stable, extensive interface with FtsWI and the L2 acceptor.

**Supplementary Table 1.** *Escherichia coli* strains used in this study

| Strain Number | Genotype (description)[a b c] | Antibiotic resistance[d] | Reference or source |
|---|---|---|---|
| AP30 | BW25113 (Xiao lab derivative of BW25113) | None | Yang et al 2017[3] |
| AP82 | Stellar pIQ-carb-*ftsW*-Halotag (stellar transformed with plasmid from lightning QC mutagenesis, plasmid annotated as pAP82) | CarbR | This Study |
| AP183[e] | BW25113 Δ*ftsW*::*kan* pBAD33-*chlor-ftsW*$^+$-*sacB* pIQ-carb-*ftsW*-Halotag (XY304 transformed with pAP82) | KanR CarbR ChlorR | This Study |
| AP751 | Stellar pIQ-carb-*ftsW*(S130C)-Halotag (stellar transformed with plasmid from lightning QC mutagenesis, plasmid annotated as pAP751) | CarbR | This Study |
| AP753 | Stellar pIQ-carb-*ftsW*(S136C)-Halotag (stellar transformed with plasmid from lightning QC mutagenesis, plasmid annotated as pAP753) | CarbR | This study |
| AP754 | Stellar pIQ-carb-*ftsW*(S130C S136C)-Halotag (stellar transformed with plasmid from lightning QC mutagenesis, plasmid annotated as pAP754) | CarbR | This study |
| AP756[e] | BW25113 Δ*ftsW*::*kan* pBAD33-*chlor-ftsW*$^+$-*sacB* pIQ-carb-*ftsW*(S130C)-Halotag (XY304 transformed with pAP751) | KanR CarbR ChlorR | This Study |
| AP758[e] | BW25113 Δ*ftsW*::*kan* pBAD33-*chlor-ftsW*$^+$-*sacB* pIQ-carb-*ftsW*(S136C)-Halotag (XY304 transformed with pAP753) | KanR CarbR ChlorR | This Study |
| AP760[e] | BW25113 Δ*ftsW*::*kan* pBAD33-*chlor-ftsW*$^+$-*sacB* pIQ-carb-*ftsW*(S130C S136C)-Halotag (XY304 transformed with pAP754) | KanR CarbR ChlorR | This Study |
| AP764 | Stellar pIQ-carb-*ftsW*(S382C)-Halotag (stellar transformed with plasmid from lightning QC mutagenesis, plasmid annotated as pAP764) | CarbR | This study |
| AP766[e] | BW25113 Δ*ftsW*::*kan* pBAD33-*chlor-ftsW*$^+$-*sacB* pIQ-carb-*ftsW*(S382C)-Halotag (XY304 transformed with pAP764) | KanR CarbR ChlorR | This study |
| AP830 | Stellar pIQ-carb-*ftsW*(L198C) (stellar transformed with plasmid from lightning QC mutagenesis, plasmid annotated as pAP830) | CarbR | This study |
| AP832 | Stellar pIQ-carb-*ftsW*(K370C)-Halotag (stellar transformed with plasmid from lightning QC mutagenesis, plasmid annotated as pAP832) | CarbR | This study |
| AP852[e] | BW25113 Δ*ftsW*::*kan* pBAD33-*chlor-ftsW*$^+$-*sacB* pIQ-carb-*ftsW*(L198C) (XY304 transformed with pAP830) | KanR CarbR ChlorR | This study |

| | | | |
|---|---|---|---|
| AP874 | Stellar pIQ-carb-*ftsW*(L198C)-Halotag (stellar transformed with plasmid from lightning QC mutagenesis, plasmid annotated as pAP874) | CarbR | This study |
| AP877 | BW25113 *ftsW*(L198C) | None | Yang et al 2021[4] (XY495) |
| AP887 | BW25113 *ftsW*(L198C) pIQ-carb-*ftsW*(L198C)-Halotag (AP877 transformed with pAP874) | CarbR | This study |
| AP892 | BW25113 pIQ-carb-*ftsW*-Halotag (AP30 transformed with pAP82) | CarbR | This study |
| AP893 | BW25113 pIQ-carb-*ftsW*(S130C S136C)-Halotag (AP30 transformed with pAP754) | CarbR | This study |
| AP894 | BW25113 pIQ-carb-*ftsW*(S382C)-Halotag (AP30 transformed with pAP764) | CarbR | This study |
| AP895 | BW25113 pIQ-carb-*ftsW*(K370C)-Halotag (AP30 transformed with pAP832) | CarbR | This study |
| AP896 | BW25113 pIQ-carb-*ftsW*(L198C)-Halotag (AP30 transformed with pAP874) | CarbR | This study |
| XY304 | BW25113 Δ*ftsW*::*kan* pBAD33-*chlor-ftsW*$^+$*-sacB* | KanR CarbR | Yang et al 2021[4] |
| XY390 | Stellar pIQ-carb-*ftsW*$^+$ (plasmid pXY390) | CarbR | Yang et al 2021[4] |
| XY396[e] | BW25113 Δ*ftsW*::*kan* pBAD33-*chlor-ftsW*$^+$*-sacB* pIQ-carb-*ftsW*$^+$ (XY304 transformed with pXY390) | KanR CarbR ChlorR | Yang et al 2021[4] |

[a] Amino-acid Linker is annotated as: GSAGSAAGSGGS

[b] The pIQ plasmid carries a ColE1 origin of replication where *ftsW* mutant alleles are expressed from an IPTG-inducible promoter under LacI repression. The pBAD33 plasmid carries a p15A origin of replication and expresses *ftsW*$^+$ from an arabinose-inducible promoter, as described by Yang *et al.* (2021)[4].

[c] Site-directed mutagenesis was performed using the QuikChange (QC) Lightning Site-Directed Mutagenesis Kit (Agilent Technologies), see Supplementary Table 2. “Stellar” refers to a commercially available strain produced by Takara Bio (Cat #636763/ 636766) based on E. coli HST08 strain made Electrocompetent for storage of plasmids.

[d] Antibiotic resistance markers: KanR, kanamycin (10 µg/mL); CarbR, carbenicillin (60 µg/mL); ChlorR, Chloramphenical (25 µg/mL).

[e] To reduce possibility of suppressor mutations arising, these strains were plated and passaged in the indicated antibiotic concentrations under conditions relying on wild-type *ftsW*$^+$ allele from the pBAD33 plasmid using 0.2% arabinose.

| **Supplementary Table 2.** Plasmids used in this study | | | |
|---|---|---|---|
| **Plasmid number** | **Genotype (description)** [a b c] | **Antibiotic resistance**[d] | **Reference or source** |
| pAP82 | ColE1, bla lacI^Q1 PT5lac-ftsW-Halotag | CarbR | This study |
| pAP751 | ColE1, bla lacI^Q1 PT5lac-ftsW(S130C)-Halotag | CarbR | This study |
| pAP753 | ColE1, bla lacI^Q1 PT5lac-ftsW(S136C)-Halotag | CarbR | This study |
| pAP754 | ColE1, bla lacI^Q1 PT5lac-ftsW(S130C S136C)-Halotag | CarbR | This study |
| pAP764 | ColE1, bla lacI^Q1 PT5lac-ftsW(S382C)-Halotag | CarbR | This study |
| pAP830 | ColE1, bla lacI^Q1 PT5lac-ftsW(L198C) | CarbR | This study |
| pAP832 | ColE1, bla lacI^Q1 PT5lac-ftsW(K370C)-Halotag | CarbR | This study |

[a] Amino-acid Linker is annotated as: GSAGSAAGSGGS

[b] The pIQ plasmid carries a ColE1 origin of replication where *ftsW* mutant alleles are expressed from an IPTG-inducible promoter under LacI repression. The pBAD33 plasmid carries a p15A origin of replication and expresses *ftsW*$^{+}$ from an arabinose-inducible promoter, as described by Yang *et al*. (2021)[4].

[c] Site-directed mutagenesis was performed using the QuikChange (QC) Lightning Site-Directed Mutagenesis Kit (Agilent Technologies), see Supplementary Table 2. "Stellar" refers to a commercially available strain produced by Takara Bio (Cat #636763/ 636766) based on E. coli HST08 strain made Electrocompetent for storage of plasmids.

[d] Antibiotic resistance markers: KanR, kanamycin (10 μg/mL); CarbR, carbenicillin (60 μg/mL); ChlorR, Chloramphenical (25 μg/mL).

| **Supplementary Table 3.** Oligonucleotide primers used in this study[a] | | | |
|---|---|---|---|
| Primer | Sequence (5' to 3') | Template | Amplicon Product |
| **For construction of pAP82 (pIQ-carb-*ftsW*-Halotag)** | | | |
| JP1 | CCCAGATCCCGCAGCAGAGCCCGCACTTCCTCGTGAACCTCGTACAAACGC | pJM003[b] | pIQ-carb-*ftsW*-Halotag |
| JP2 | GGAAGTGCGGGCTCTGCTGCGGGATCTGGGGGATCCGAAATCGGTACTGGCT | | |
| **For construction of pAP751 (pIQ-carb-*ftsW(*S130C*)*-Halotag)** | | | |
| JP268 | AGATCGATCCAACGCGATGCACCTTTAACCGAGCAACCCACTACCAGGACGATCA | pAP82 | pIQ-carb-*ftsW(S130C)*-Halotag |
| JP272 | GCATCGCGTTGGATCGATCT | | |
| **For construction of pAP753 (pIQ-carb-*ftsW(*S136C*)*-Halotag)** | | | |
| JP270 | AAACCGAGATCGATCCAACGGCATGCACCTTTAACCGAGCTACCCACTAC | pAP82 | pIQ-carb-*ftsW(S136C)*-Halotag |
| JP273 | CGTTGGATCGATCTCGGTTT | | |
| **For construction of pAP754 (pIQ-carb-*ftsW(*S130C S136C*)*-Halotag)** | | | |
| JP271 | AAACCGAGATCGATCCAACGGCATGCACCTTTAACCGAGCAACCCACTACCAGGACGATC | pAP82 | pIQ-carb-*ftsW(S130C S136C)*-Halotag |
| JP273 | CGTTGGATCGATCTCGGTTT | | |
| **For construction of pAP764 (pIQ-carb-*ftsW(*S382C*)*-Halotag)** | | | |
| JP280 | GACATAATCAGTAAGCTGCAACCACCGTAACTGATCAGCG | pAP82 | pIQ-carb-*ftsW(S382C)*-Halotag |
| JP281X | AGCTTACTGATTATGTCGAC | | |
| **For construction of pAP830 (pIQ-carb-*ftsW*(L198C))** | | | |
| JP276 | AACACCACCACCGTACCACAGTCTGGCTGTGCCAGCAGTA | pAP82 | pIQ-carb-*ftsW(L198C)* |
| JP298 | TGTGGTACGGTGGTGGTGTTGTT | | |

[a] Lightning QC PCR (Agilent) was used to construct DNA used to transform into stellar cells.

[b] pJM003 Xiao lab plasmid contains pIQ-*ftsW*-$L_{GPG}$-Halotag with a Glycine-Proline Linker used as template for Lightning QC PCR.

**Supplementary Table 4: Cell lengths of FtsW-depletion strains complemented with ftsW+ (XY396) or ftsW(L198C) (AP852), grown in M9-glucose at 25 °C (Fig. S7A).**

| Genotype and condition | Replicate medians (µm) | Geometric mean ± s.e.m. (µm) | Pooled median (µm) | n repl. | n cells | Elongated cells, mean % (replicates) | Comparison: ratio of geometric means (95% bootstrap interval) | Result |
|---|---|---|---|---|---|---|---|---|
| ftsW+ +IPTG | 3.02, 2.72, 3.01 | 2.91 ± 0.10 | 2.92 | 3 | 184 | 0.6 (0.0, 1.9, 0.0) | reference | |
| ftsW+ +IPTG, +MTSES | 3.05, 3.24, 3.01 | 3.10 ± 0.07 | 3.09 | 3 | 162 | 3.6 (0.0, 10.7, 0.0) | vs ftsW+ +IPTG: 1.06 (0.966–1.20) | – |
| ftsW+ −IPTG | 5.4, 5.7, 17.5 | 8 ± 4 | 6.54 | 3 | 181 | 56.2 (36.9, 50.0, 81.7) | vs ftsW+ +IPTG: 2.78 (1.75–5.60) | * |
| ftsW(L198C) +IPTG | 4.1, 4.0, 3.5 | 3.9 ± 0.2 | 3.91 | 3 | 169 | 6.6 (7.8, 6.7, 5.2) | vs ftsW+ +IPTG: 1.33 (1.18–1.51) | * |
| ftsW(L198C) +IPTG, +MTSES | 18.0, 16.3, 29.1 | 20 ± 4 | 19.7 | 3 | 143 | 81.6 (73.0, 81.1, 90.6) | vs ftsW(L198C) +IPTG: 5.27 (2.56–8.27)<br>vs ftsW+ +IPTG, +MTSES: 6.59 (3.31–9.97) | *<br>* |
| ftsW(L198C) −IPTG | 38.8, 17.7, 18.5 | 23 ± 7 | 20.5 | 3 | 113 | 92.6 (94.4, 97.3, 86.2) | vs ftsW(L198C) +IPTG: 6.01 (4.06–9.48) | * |

Cell lengths in µm. Each biological replicate is summarised by its median; the geometric mean ± s.e.m. is calculated over the replicate medians and is the basis for the comparisons. Pooled median is the median of all cells in the group, imaging days pooled. Elongated cells: a cell was scored as elongated if longer than 5.84 µm, which is twice the pooled median of the ftsW+ +IPTG control group; the value is the mean of the replicate percentages, with the individual replicate percentages in parentheses. Comparison: ratio of the geometric means of replicate medians, test group / reference group, with the 95% interval of a two-level percentile bootstrap (cells resampled with replacement within each replicate, then replicates resampled with replacement; 40,000 iterations; seed 11). *, the interval excludes 1; –, the interval includes 1. This is descriptive and not a hypothesis test. n repl., number of independent biological replicates (n = 2 or 3). Values are given to three significant figures. +IPTG, complementing allele induced; +MTSES, MTSES added; −IPTG, complementing allele not induced (FtsW depleted).

**Supplementary Table 5: Single-molecule tracking population statistics for FtsW cysteine variants ± MTSES.**

| Metric | FtsW L198C-AP887 | FtsW L198C-AP887 + MTSES | FtsW S382C-AP766 | FtsW S382C-AP766 + MTSES | FtsW-AP826 (WT) |
|---|---|---|---|---|---|
| Trajectory segments (N) | 313 | 314 | 109 | 181 | 544 |
| Stationary (n) | 215 | 205 | 69 | 124 | 308 |
| Moving (n) | 98 | 109 | 40 | 57 | 236 |
| P(stationary) (%) | 68.7 ± 2.6 (215) | 65.3 ± 2.6 (205) | 63.3 ± 4.4 (69) | 68.5 ± 3.4 (124) | 56.6 ± 2.1 (308) |
| τ(stat) (s) | 46.2 ± 3.9 | 35.7 ± 4.1 | 45.5 ± 7.0 | 51.6 ± 6.6 | 13.8 ± 0.8 |
| P(slow-moving) (%) | 13.4 ± 1.9 (42) | 2.2 ± 0.8 (7) | 24.8 ± 4.1 (27) | 2.8 ± 1.2 (5) | 21.5 ± 1.7 (117) |
| v(sm) (nm/s) | 4.3 ± 0.6 | 3.3 ± 1.0 | 3.4 ± 0.3 | 3.3 ± 1.5 | 7.0 ± 0.3 |
| τ(sm) (s) | 74.7 ± 9.8 | 83.2 ± 37.2 | 53.5 ± 6.9 | 86.6 ± 36.7 | 29.0 ± 1.8 |
| PL(sm) (nm) | 139 ± 11 | 89 ± 10 | 153 ± 21 | 92 ± 12 | 182 ± 12 |
| P(fast-moving) (%) | 17.9 ± 2.1 (56) | 32.5 ± 2.6 (102) | 11.9 ± 3.2 (13) | 28.7 ± 3.3 (52) | 21.9 ± 1.7 (119) |
| v(fm) (nm/s) | 49.6 ± 6.1 | 46.6 ± 4.7 | 49.5 ± 16.4 | 29.8 ± 5.5 | 52.3 ± 3.6 |
| τ(fm) (s) | 5.9 ± 0.6 | 7.9 ± 0.8 | 6.5 ± 1.0 | 13.8 ± 2.5 | 5.9 ± 0.4 |
| PL(fm) (nm) | 210 ± 19 | 202 ± 14 | 227 ± 51 | 172 ± 22 | 250 ± 16 |

**Supplementary Table 6: Cell lengths of ftsW-Halo and ftsW(SSCC)-Halo strains grown in EZRDM at 30 °C, then on LB plates with 100 µM IPTG, without or with 10 mM DTT (Fig. S7B).**

| Genotype and condition | Replicate medians (µm) | Geometric mean ± s.e.m. (µm) | Pooled median (µm) | n repl. | n cells | Elongated cells, mean % (replicates) | Comparison: ratio of geometric means (95% bootstrap interval) | Result |
|---|---|---|---|---|---|---|---|---|
| ftsW-Halo −DTT | 2.38, 2.59, 2.39 | 2.45 ± 0.07 | 2.48 | 3 | 141 | 4.1 (2.4, 6.0, 4.0) | reference | |
| ftsW-Halo +DTT | 2.41, 2.98, 2.89 | 2.7 ± 0.2 | 2.76 | 3 | 150 | 8.0 (0.0, 6.0, 18.0) | vs ftsW-Halo −DTT: 1.12 (0.956–1.30) | – |
| ftsW(SSCC)-Halo −DTT | 5.88, 30.6, 8.95 | 12 ± 8 | 10.7 | 3 | 149 | 77.2 (62.0, 94.0, 75.5) | vs ftsW-Halo −DTT: 4.78 (2.46–10.8) | * |
| ftsW(SSCC)-Halo +DTT | 2.51, 5.5, 3.76 | 3.7 ± 0.8 | 3.65 | 3 | 150 | 30.0 (4.0, 64.0, 22.0) | vs ftsW(SSCC)-Halo −DTT: 0.319 (0.182–0.489)<br>vs ftsW-Halo +DTT: 1.36 (1.03–1.82) | *<br>* |

Cell lengths in µm. Each biological replicate is summarised by its median; the geometric mean ± s.e.m. is calculated over the replicate medians and is the basis for the comparisons. Pooled median is the median of all cells in the group, imaging days pooled. Elongated cells: a cell was scored as elongated if longer than 4.96 µm, which is twice the pooled median of the ftsW-Halo −DTT control group; the value is the mean of the replicate percentages, with the individual replicate percentages in parentheses. Comparison: ratio of the geometric means of replicate medians, test group / reference group, with the 95% interval of a two-level percentile bootstrap (cells resampled with replacement within each replicate, then replicates resampled with replacement; 40,000 iterations; seed 11). *, the interval excludes 1; –, the interval includes 1. This is descriptive and not a hypothesis test. n repl., number of independent biological replicates (n = 2 or 3). Values are given to three significant figures. −DTT, +DTT, without or with 10 mM DTT on the LB plate; SSCC, S130C S136C.

**Supplementary Table 7: Cell lengths of ECL2 cysteine variants in EZRDM at 37 °C, 50 µM IPTG, without DTT (Fig. S16D).**

| Genotype and condition | Replicate medians (µm) | Geometric mean ± s.e.m. (µm) | Pooled median (µm) | n repl. | n cells | Elongated cells, mean % (replicates) | Comparison: ratio of geometric means (95% bootstrap interval) | Result |
|---|---|---|---|---|---|---|---|---|
| BW25113 (untagged parent) | 4.4, 3.0, 4.0 | 3.8 ± 0.4 | 3.90 | 3 | 221 | 0.0 (0.0, 0.0, 0.0) | vs ftsW-Halo (AP183): 0.833 (0.773–0.906) | * |
| ftsW-Halo (AP183) | 5.0, 3.7, 5.0 | 4.5 ± 0.5 | 4.58 | 3 | 278 | 9.9 (25.9, 0.0, 3.9) | reference | |
| ftsW(S130C) (AP756) | 4.9, 3.6, 4.2 | 4.2 ± 0.4 | 4.31 | 3 | 267 | 1.9 (1.9, 2.4, 1.3) | vs ftsW-Halo (AP183): 0.931 (0.831–1.02) | – |
| ftsW(S136C) (AP758) | 6.2, 4.3, 5.0 | 5.1 ± 0.5 | 5.24 | 3 | 231 | 13.2 (30.6, 1.9, 7.0) | vs ftsW-Halo (AP183): 1.14 (1.00–1.30) | *† |
| FtsW(SSCC) (AP760) | 47.2, 22.3, 64.8 | 41 ± 12 | 40.9 | 3 | 217 | 95.0 (95.9, 89.1, 100.0) | vs ftsW-Halo (AP183): 9.07 (5.91–13.7) | * |

Cell lengths in µm. Each biological replicate is summarised by its median; the geometric mean ± s.e.m. is calculated over the replicate medians and is the basis for the comparisons. Pooled median is the median of all cells in the group, imaging days pooled. Elongated cells: a cell was scored as elongated if longer than 7.80 µm, which is twice the pooled median of the BW25113 control group; the value is the mean of the replicate percentages, with the individual replicate percentages in parentheses. Comparison: ratio of the geometric means of replicate medians, test group / reference group, with the 95% interval of a two-level percentile bootstrap (cells resampled with replacement within each replicate, then replicates resampled with replacement; 40,000 iterations; seed 11). *, the interval excludes 1; –, the interval includes 1. This is descriptive and not a hypothesis test. n repl., number of independent biological replicates (n = 2 or 3). Values are given to three significant figures. AP760 carries both substitutions (S130C and S136C); the single-cysteine strains AP756 and AP758 divide normally under the same conditions. †, the interval touches 1.0 and the call changes with the bootstrap seed; the figure marks these comparisons with a dashed bracket.

**Supplementary Table 8: Peptide identifications and sequence coverage for HaloTag-enriched FtsW-Halo variants.**

| Sample | Peptides identified | Coverage of FtsW-Halo fusion (%) |
|---|---|---|
| FtsW-Halo (WT) | 107 | 64.9 |
| FtsW(S382C)-Halo | 105 | 64.5 |
| FtsW(L198C)-Halo | 109 | 63.1 |
| FtsW(SSCC)-Halo | 28 | 24.7 |
| FtsW(S382C)-Halo + MTSES | 7 | 10.4 |
| FtsW(L198C)-Halo + MTSES | 19 | 21.4 |

Peptides identified and sequence coverage of the 721-residue FtsW-Halo fusion, from the analysis shown in Figure S10. Coverage is reported for the full fusion; the HaloTag portion was covered more completely than the FtsW portion in all samples.

### Supplementary Table 9: Cell lengths of FtsW-depletion strains complemented with ftsW-Halo (AP183) or ftsW(S382C)-Halo (AP766), grown in EZRDM at 37 °C (Fig. S7C).

| Genotype and condition | Replicate medians (µm) | Geometric mean ± s.e.m. (µm) | Pooled median (µm) | n repl. | n cells | Elongated cells, mean % (replicates) | Comparison: ratio of geometric means (95% bootstrap interval) | Result |
|---|---|---|---|---|---|---|---|---|
| ftsW-Halo +IPTG | 3.7, 5.8 | 5 ± 1 | 4.32 | 2 | 144 | 3.4 (0.0, 6.8) | reference | |
| ftsW-Halo +IPTG, +MTSES | 4.3, 6.0 | 5.1 ± 0.8 | 5.09 | 2 | 115 | 1.8 (0.0, 3.5) | vs ftsW-Halo +IPTG: 1.09 (0.943–1.26) | – |
| ftsW-Halo −IPTG | 9.5, 16.5 | 13 ± 3 | 11.9 | 2 | 109 | 77.5 (62.5, 92.5) | vs ftsW-Halo +IPTG: 2.70 (2.35–3.39) | * |
| ftsW(S382C) +IPTG | 5.6, 5.7, 6.5 | 5.9 ± 0.3 | 5.75 | 3 | 210 | 10.3 (2.4, 12.7, 15.9) | vs ftsW-Halo +IPTG: 1.22 (0.904–1.64) | – |
| ftsW(S382C) +IPTG, +MTSES | 13.5, 36.4, 28.2 | 24 ± 7 | 27.3 | 3 | 161 | 89.1 (73.1, 98.3, 96.0) | vs ftsW(S382C) +IPTG: 4.07 (2.57–6.46)<br>vs ftsW-Halo +IPTG, +MTSES: 4.38 (2.76–6.81) | *<br>* |
| ftsW(S382C) −IPTG | 15.7, 22.5 | 19 ± 3 | 19.5 | 2 | 114 | 93.9 (93.3, 94.4) | vs ftsW(S382C) +IPTG: 3.33 (2.45–4.47) | * |

Cell lengths in µm. Each biological replicate is summarised by its median; the geometric mean ± s.e.m. is calculated over the replicate medians and is the basis for the comparisons. Pooled median is the median of all cells in the group, imaging days pooled. Elongated cells: a cell was scored as elongated if longer than 8.63 µm, which is twice the pooled median of the ftsW-Halo +IPTG control group; the value is the mean of the replicate percentages, with the individual replicate percentages in parentheses. Comparison: ratio of the geometric means of replicate medians, test group / reference group, with the 95% interval of a two-level percentile bootstrap (cells resampled with replacement within each replicate, then replicates resampled with replacement; 40,000 iterations; seed 11). *, the interval excludes 1; –, the interval includes 1. This is descriptive and not a hypothesis test. n repl., number of independent biological replicates (n = 2 or 3). Values are given to three significant figures. +IPTG, complementing allele induced; +MTSES, MTSES added; −IPTG, complementing allele not induced (FtsW depleted). Groups with n = 2 replicates (the first three in the table and ftsW(S382C)-Halo −IPTG) are reported as measured; comparisons involving them rest on two replicates.

**Supplementary Table 10: Cell lengths of dominant-negative strains expressing ftsW-Halo alleles from pIQ plasmids in a wild-type ftsW background, grown in EZRDM at 37 °C without or with 100 µM MTSES (Fig. S9C).**

| Genotype and condition | Replicate medians (µm) | Geometric mean ± s.e.m. (µm) | Pooled median (µm) | n repl. | n cells | Elongated cells, mean % (replicates) | Comparison: ratio of geometric means (95% bootstrap interval) | Result |
|---|---|---|---|---|---|---|---|---|
| ftsW-Halo −MTSES | 5.21, 4.26, 5.19 | 4.9 ± 0.3 | 4.84 | 3 | 132 | 0.0 (0.0, 0.0, 0.0) | reference | |
| ftsW-Halo +MTSES | 4.95, 4.46, 5.19 | 4.9 ± 0.2 | 4.81 | 3 | 130 | 3.6 (8.6, 2.3, 0.0) | reference | |
| ftsW(SSCC)-Halo −MTSES | 6.57, 5.94, 7.28 | 6.6 ± 0.4 | 6.50 | 3 | 145 | 12.9 (3.0, 7.8, 27.9) | vs ftsW-Halo −MTSES: 1.35 (1.20–1.52) | * |
| ftsW(S382C)-Halo −MTSES | 6.01, 5.23, 5.76 | 5.7 ± 0.2 | 5.76 | 3 | 139 | 1.9 (0.0, 1.9, 3.8) | vs ftsW-Halo −MTSES: 1.16 (1.08–1.31) | * |
| ftsW(S382C)-Halo +MTSES | 5.93, 5.27, 6.13 | 5.8 ± 0.3 | 5.77 | 3 | 154 | 3.7 (9.5, 1.6, 0.0) | vs ftsW-Halo +MTSES: 1.19 (1.12–1.35) | * |
| ftsW(K370C)-Halo −MTSES | 73.0, 83.8, 66.0 | 74 ± 5 | 72.7 | 3 | 29 | 100.0 (100.0, 100.0, 100.0) | vs ftsW-Halo −MTSES: 15.2 (9.74–19.7) | * |
| ftsW(L198C)-Halo −MTSES | 8.99, 6.56, 9.74 | 8 ± 1 | 8.04 | 3 | 143 | 35.7 (35.3, 21.8, 50.0) | vs ftsW-Halo −MTSES: 1.71 (1.44–2.00) | * |
| ftsW(L198C)-Halo +MTSES | 9.86, 12.7, 11.8 | 11.4 ± 0.9 | 12.1 | 3 | 161 | 60.1 (53.1, 68.5, 58.6) | vs ftsW(L198C)-Halo −MTSES: 1.37 (1.00–2.11)<br>vs ftsW-Halo +MTSES: 2.34 (1.86–3.51) | *†<br>* |

Cell lengths in µm. Each biological replicate is summarised by its median; the geometric mean ± s.e.m. is calculated over the replicate medians and is the basis for the comparisons. Pooled median is the median of all cells in the group, imaging days pooled. Elongated cells: a cell was scored as elongated if longer than 9.69 µm, which is twice the pooled median of the ftsW-Halo −MTSES control group; the value is the mean of the replicate percentages, with the individual replicate percentages in parentheses. Comparison: ratio of the geometric means of replicate medians, test group / reference group, with the 95% interval of a two-level percentile bootstrap (cells resampled with replacement within each replicate, then replicates resampled with replacement; 40,000 iterations; seed 11). *, the interval excludes 1; –, the interval includes 1. This is descriptive and not a hypothesis test. n repl., number of independent biological replicates (n = 2 or 3). Values are given to three significant figures. All strains carry chromosomal wild-type ftsW and the indicated pIQ-ftsW-Halo allele (100 µM IPTG); −MTSES, +MTSES, without or with 100 µM MTSES; SSCC, S130C S136C. ftsW(K370C)-Halo was measured from 29 cells across three replicates because its filaments were extremely long and few. †, the interval touches 1.0 and the call changes with the bootstrap seed; the figure marks these comparisons with a dashed bracket.

**Supplementary Table 11:** Predefined mechanistic positions among top-ranked LigBind predictions in representative FtsW orthologs

| UniProt ID | Species | Top-15[a] | Top-20[a,b] |
|---|---|---|---|
| P0ABG4 | E. coli | 4/12 | 7/12 |
| Q9HW00 | P. aeruginosa | 5/12 | 5/12 |
| O07639 | B. subtilis | 5/12 | 5/12 |
| B8H092 | Caulobacter crescentus | 4/12 | 4/12 |
| A0A378ACG3 | K. pneumoniae | 4/12 | 7/12 |
| **Combined** | **Five proteins** | **22/60** | **28/60** |
| **Combined percentage** | | **36.7%** | **46.7%** |

[a] *A position was counted at Top-k when its LigBind rank was less than or equal to k*

[b] *Across the five orthologs, 28 of 60 predefined positions occurred among the Top-20 predictions. For reference, uniform random selection of 20 residues from each sequence would include approximately 3 of the 60 positions on average. This comparison is descriptive only.*

**Supplementary Table 12:** Family-wide recurrence of AiPP-LigBind predictions at predefined FtsW mechanistic positions.

| Position | n[a] | Exact Top-20 (%)[b] | Adjacent-only ±2 (%)[c] |
|---|---|---|---|
| R137 | 6839 | 99.23 | 15.59 |
| R243 | 6845 | 83.14 | 90.05 |
| R246 | 6819 | 99.68 | 0.26 |
| D297 | 6852 | 93.16 | 94.02 |
| L198 | 6845 | 24.92 | 99.93 |
| L236 | 6264 | 0.05 | 0.21 |
| K370 | 6848 | 37.82 | 94.44 |
| R84 | 6825 | 0.00 | 0.04 |
| W138 | 6838 | 20.25 | 0.06 |
| R145 | 3982 | 0.00 | 43.47 |
| Y379 | 6843 | 46.95 | 99.91 |
| S382 | 6840 | 57.27 | 99.08 |

[a] Here, n is the number of orthologs containing an unambiguous residue at the corresponding aligned position.

[b] Exact recurrence is the percentage of these orthologs in which that residue ranked among the Top-20 predictions.

[c] Adjacent-only recurrence reports a Top-20 non-mechanistic residue within ±2 positions in primary sequence after excluding all predefined mechanistic residues.
